# Single-cell type analysis of wing premotor circuits in the ventral nerve cord of *Drosophila melanogaster*

**DOI:** 10.1101/2023.05.31.542897

**Authors:** Erica Ehrhardt, Samuel C Whitehead, Shigehiro Namiki, Ryo Minegishi, Igor Siwanowicz, Kai Feng, Hideo Otsuna, FlyLight Project Team, Geoffrey W Meissner, David Stern, James W. Truman, David Shepherd, Michael H. Dickinson, Kei Ito, Barry J Dickson, Itai Cohen, Gwyneth M Card, Wyatt Korff

## Abstract

To perform most behaviors, animals must send commands from higher-order processing centers in the brain to premotor circuits that reside in ganglia distinct from the brain, such as the mammalian spinal cord or insect ventral nerve cord. How these circuits are functionally organized to generate the great diversity of animal behavior remains unclear. An important first step in unraveling the organization of premotor circuits is to identify their constituent cell types and create tools to monitor and manipulate these with high specificity to assess their functions. This is possible in the tractable ventral nerve cord of the fly. To generate such a toolkit, we used a combinatorial genetic technique (split-GAL4) to create 195 sparse transgenic driver lines targeting 196 individual cell types in the ventral nerve cord. These included wing and haltere motoneurons, modulatory neurons, and interneurons. Using a combination of behavioral, developmental, and anatomical analyses, we systematically characterized the cell types targeted in our collection. In addition, we identified correspondences between the cells in this collection and a recent connectomic data set of the ventral nerve cord. Taken together, the resources and results presented here form a powerful toolkit for future investigations of neuronal circuits and connectivity of premotor circuits while linking them to behavioral outputs.

## Introduction

For animals to survive and reproduce, they must perform precisely controlled movements under the direction of the nervous system. Most of these motor behaviors in insects are facilitated by the vertebrate spinal cord analog, the ventral nerve cord (VNC). The VNC receives and processes sensory information and is involved in generating most of the locomotor actions that underlie fly behaviors such as walking (Bidaye et al., 2014, Tuthill and Wilson, 2016, Chen et al., 2018, Howard et al., 2019), grooming (Seeds et al., 2014), escape (King and Wyman, 1980, Trimarchi and Schneiderman, 1995, Card and Dickinson, 2008, von Reyn et al., 2014), flight (Dickinson and Muijres, 2016, Schnell et al., 2017, O’Sullivan et al., 2018, Namiki et al., 2022), courtship (Clyne and Miesenbock, 2008, von Philipsborn et al., 2011, Shirangi et al., 2013, Shiozaki et al., 2024, Lillvis et al., 2024), and copulation (Crickmore and Vosshall, 2013, Pavlou et al., 2016).

The *Drosophila* VNC (Figure 1A,B), a set of fused ganglia that contain motoneurons for muscles that move the wings, legs, and other body parts, is a promising model in which to investigate premotor circuits at the single neuron level because it is numerically tractable (on the order of 23,000 neurons (Takemura et al., 2023) and amenable to manipulation using genetic tools. Premotor circuits in the VNC receive sensory input from peripheral organs in the wings, legs, halteres, and abdomen (Tsubouchi et al., 2017) and descending commands from the brain (Namiki et al., 2018, Shiozaki et al., 2024). Within the microcircuits of the VNC, these multimodal inputs are transformed into patterns of motoneuron activity that produce adaptive motor actions. VNC neurons also transmit information to the brain via ascending pathways (Tsubouchi et al., 2017). Understanding the neuronal basis of motor behavior is contingent upon a detailed understanding of the neuronal circuitry that makes up the VNC; however, despite recent advances in both functional recording (Chen et al., 2018, Shiozaki et al., 2024) and connectomic mapping of this region (Kuan et al., 2020, Azevedo et al., 2024, Cheong et al., 2023, Lesser et al., 2023, Takemura et al., 2023, Marin et al., 2023, Shiozaki et al., 2024, Lillvis et al., 2024), relatively little is known about the functional organization of VNC local circuitry (Venkatasubramanian and Mann, 2019).

**Figure 1:**
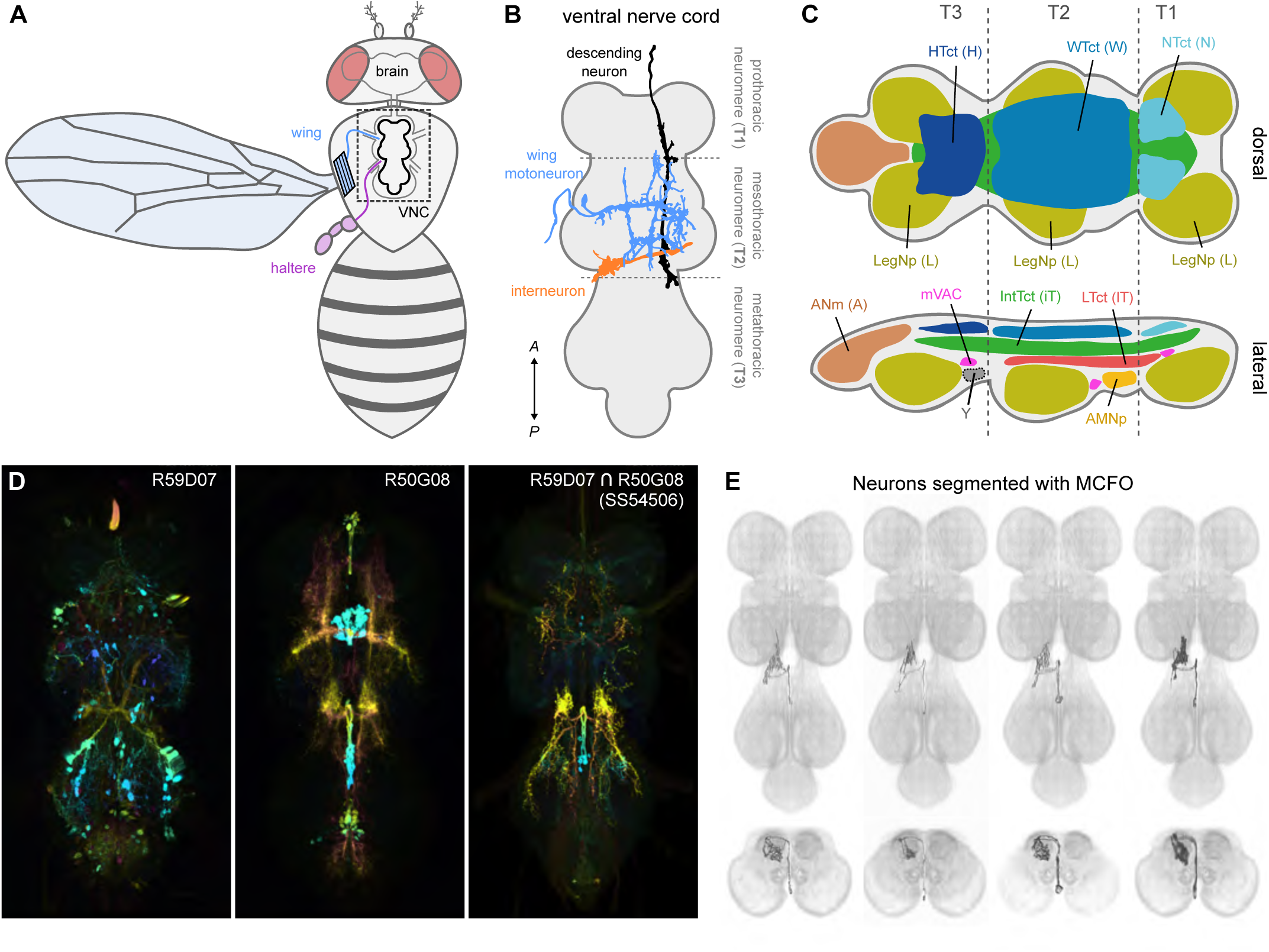
Isolating neurons in the ventral nerve cord (VNC). (**A**) The fly central nervous system (gray) with the ventral nerve cord (VNC) highlighted in black. Illustrations of wing motoneuron and haltere sensory afferent shown in blue and purple, respectively. (**B**) Top-down (dorsal) view of VNC showing example neuron types: wing motoneuron (blue), descending neuron (black), and haltere-to-wing neuropil interneuron (orange). Boundaries between the pro-, meso-, and metathoracic neuromeres—i.e. T1, T2, and T3—are also shown (gray dotted lines and labels). (**C**) Schematic of VNC neuropils. Abbreviations used: T1 (prothoracic segment), T2 (mesothoracic segment), T3 (metathoracic segment), VAC (ventral association center), mVAC (medial ventral association center), and AMNp (accessory mesothoracic neuropil). (**D**) Example usage of the split-GAL4 technique for narrowing driver line expression profile (white) in the VNC (orange). R59G07 (**D_1_**) and R50G08 (**D_2_**) are used to drive half of the GAL4 transcription factor in SS54506 (**D_3_**), resulting in a sparse expression pattern. (**E**) Multiple interneurons segmented from the expression pattern of SS54506 using multicolor flip-out (MCFO).

For flying insects, such as the fruit fly, *Drosophila melanogaster*, wing behaviors comprise some of the most fascinating examples of motor control. In flight, subtle adjustments to wing motions can have large aerodynamic consequences that allow flies to rapidly evade predators (Muijres et al., 2014), effectively forage in complex environments (van Breugel and Dickinson, 2014), and even traverse distances as large as ∼15 km in a single flight bout (Coyne et al., 1982, Leitch et al., 2020). On the ground, male flies’ tightly patterned and subtle small wing vibrations create a species-specific courtship song necessary to attract mates (Ewing, 1979). Control of behaviors for a given appendage, such as the wing, must use the same limited set of motoneurons and muscles (Dickinson and Tu, 1997, Lindsay et al., 2017, O’Sullivan et al., 2018), but may produce very different motions. For example, flight requires large, synchronized sweeping movements of both wings, whereas song is generated via small vibrations of a single wing. Moreover, flight and song occur in entirely different behavioral contexts. Just how premotor microcircuits in the VNC generate distinct, context-dependent patterns of motor activity using the same set of motoneurons is not well understood (O’Sullivan et al., 2018).

To execute these wing behaviors, flies make use of a highly specialized wing motor system, whose constituent muscles can be broadly separated into two distinct categories: the large, asynchronous power muscles and the small, synchronous control muscles. The power muscles comprise two antagonistic sets of muscle fibers per side, which fill the thorax and provide the primary drive for wing movement. While they receive innervation from motoneurons, the power muscles are stretch-activated, and contract at a frequency determined by the resonant properties of the insect’s thorax rather than the spike rate of motoneuron inputs (Boettiger, 1960, Pringle, 1949, Hürkey et al., 2023). In contrast, the smaller control muscles contract on a 1:1 basis with motoneuron spikes and drive more subtle changes to wing movement.

The set of neurons responsible for local—i.e. within-VNC—control of this wing motor system includes motoneurons, non-motor efferent neurons, and interneurons. Motoneurons receive input from premotor neurons in the nerve cord, extend axons through nerves that exit the central nervous system, and terminate in the periphery, where they innervate muscles. They thus comprise the final link between neuronal activity and motion, and any dissection of premotor circuits should be anchored by an analysis of motoneuron function. Motoneurons are also the easiest neuron type to identify within a given motor circuit, as they can be characterized by their muscle of termination. Yet for many common behaviors, exactly which motoneurons are activated and with what dynamic pattern is still unknown. This is especially true for behaviors involving appendages that must be moved through complex biomechanical articulations, such the fly wing hinge, and in very different kinematic patterns to produce different behaviors, such as flapping versus singing.

In addition to excitatory motoneurons, Dipteran wing muscles are innervated by mesothoracic ventral unpaired median (T2VUM) neurons, non-motor efferent neuromodulatory cells that play a crucial role in coordinated wing behaviors (Sadaf et al., 2015, Schlurmann and Hausen, 2003). These T2VUM cells are members of a broader category of largely efferent neurons called ventral unpaired median (VUM) neurons, so named for their somata, which cluster along the midline of the VNC in the ventral cortex, and their bilaterally symmetric neurite projections (Schlurmann and Hausen, 2003). While a recently identified VUM cell in *Drosophila* has been shown to be dopaminergic (Sadaf et al., 2015), octopaminergic VUM neurons innervating wing muscles have been identified in several other insect species, including *Calliphora* (blowflies) and *Schistocerca* (desert locusts), and may be a general feature of the neuronal circuitry controlling insect wings (Schlurmann and Hausen, 2003, Stocker et al., 2018).

Interneurons are a broad class of cells that reside entirely in the VNC and constitute much of the circuitry required for executing and controlling motor behaviors. The functional roles of VNC interneurons are, however, largely unexplored, in part because they are difficult to access and individually address (Venkatasubramanian and Mann, 2019). Thus, transgenic driver lines that target such VNC interneurons will be essential to unraveling the functional organization of the local VNC circuits that give rise to complex wing behaviors (Shiozaki et al., 2024, Lillvis et al., 2024).

To facilitate systematically identifying and visualizing the constituent neurons in the local premotor circuits of the VNC, we applied recent advances in combinatorial genetic techniques (Figure 1D) (Jenett et al., 2012, Luan et al., 2006) to generate a large-scale collection of transgenic flies that can be used to target individual VNC neurons for both visualization and *in vivo* manipulation as has been done in other regions of the fly nervous system (Gao et al., 2008, Aso et al., 2014, Wolff et al., 2015, Wu et al., 2016, Namiki et al., 2018, Taisz et al., 2023, Meissner et al., 2025). The VNC can be broadly subdivided into dorsal and ventral regions: dorsal VNC neuropils control the neck, wings, and halteres in the execution of behaviors like flight and courtship song, while ventral VNC neuropils control the legs for behaviors like walking and grooming (Figure 1C) (Court et al., 2020). Here we focus on the dorsal VNC neuropils associated with wing behaviors. We created 195 cell-type specific *Drosophila melanogaster* split-GAL4 driver lines targeting motoneurons, the modulatory ventral unpaired median neurons (VUMs), and interneurons in this half of the VNC (Figure 1D,E) (Court et al., 2020, Namiki et al., 2018). Ascending and sensory neurons that innervate the VNC are beyond the scope of this effort. This library should enable researchers to probe the premotor circuits controlling the rich set of behaviors requiring wing, neck, or haltere coordination, such as flight or courtship.

To help catalyze the use of this library, we performed a combination of behavioral assays and quantitative anatomical analyses to characterize the dorsal VNC organization functionally. Using reagents targeting wing motoneurons, we mapped individual motoneuron manipulations to specific behavioral phenotypes in both flight and courtship song behavioral assays. Using light microscopy and multi-color flip out (MCFO) single-cell labeling (Figure 1E), in tandem with reagents targeting dorsal VNC neurons, we segmented 196 cells and annotated both their developmental origins (Doe, 1992, Truman et al., 2010) and their sites of input and output. We then used this information to identify matches for the segmented cells in the *Drosophila* male VNC connectome (MANC; (Takemura et al., 2023)), facilitating connectivity analyses of these neurons. Together, our results lay the groundwork for a basic functional architecture of the neuronal circuitry controlling wing movements and provide an important resource for future investigations of the neural substrates underlying motor behavior.

## Results

### Creation of a library of split-GAL4 lines

We used the split-GAL4 system to generate a collection of driver lines with sparse expression in dorsal VNC neurons (Figure 1D,E; Figure 1—figure supplements 1-5). In the split-GAL4 system, two different gene expression regulatory elements each drive the expression of one domain of the GAL4 transcription factor (the activation domain, AD, or DNA binding domain, DBD), such that only cells expressing both domains produce a functional GAL4 protein and transcribe the gene of interest that is downstream of the GAL4 binding target, the Upstream Activation Sequence (UAS). This combinatorial approach can produce sparse, even cell-type specific, expression patterns. To produce driver lines with expression in only a single neuronal cell type, we screened the publicly available database of Janelia Gen1 GAL4 lines (Pfeiffer et al., 2008, Pfeiffer et al., 2010, Jenett et al., 2012, Meissner et al., 2023) for pairs of lines that had expression in the same cell type using an automated protocol (Otsuna et al., 2018). From this search, we identified and performed 1,658 AD/DBD candidate intersections, which produced 195 combinations with sufficiently sparse expression to use to make homozygous transgenic lines for inclusion in our collection (Figure 1—figure supplements 1-5).

We identified 196 unique dorsal VNC cell types (Table 1) targeted by these 195 stabilized driver lines (Table 2). These included 12 wing power muscle motoneurons (in 16 driver lines), 16 wing control muscle motoneurons (in 35 driver lines), 5 haltere muscle motoneurons (in 15 driver lines), 5 ventral unpaired median (VUM) neurons (in 8 driver lines), 72 intrasegmental interneurons primarily innervating dorsal neuropil regions (in 64 driver lines) and 85 intersegmental interneurons primarily innervating the dorsal neuropils (in 81 driver lines). While generally sparse, many of our driver lines contained multiple cell types. We used the multi-color flip out (MCFO) stochastic labeling technique (Nern et al., 2015) to visualize different cell types expressed in a single driver in different colors, and we used this high-resolution, single-cell image data to create a database of their morphologies (Figure 1E; Figure 1—figure supplements 1-5).

**Table 1:** List of neurons identified in the study. The number of cell bodies, cell body locations and numbers, hemilineages of origin, areas of arborizations, existence of axon-like projections, the driver lines that label the cells and alternative names previously used in the literature are listed for all the neuron types. Loc.: location of the cell bodies: 1, 2, 3, A, associated with T1, T2, T3 or Abdominal ganglia; a, anterior; p, posterior, v, ventral; d, dorsal, m, medial; l, lateral;. No.: most plausible number of the cells of that type based on the comparison of the driver line labeling samples; actual numbers in the VNC may differ. For unpaired cells, the total number of cells is given in italics; for paired cells, the number of cells per side is given in plain text. Lineage: most plausible cell lineage based on the comparison of the neuroblast labeling: emb, embryonic; abd, abdominal. Areas of arborizations: first character represents arborization ipsilateral to the soma, second character represents arborization contralateral to the soma. A, axon terminals, D, dendritic arbors, M, intermixed input and output sites, P, input and output sites partitioned in separate areas within the same hemi-neuropil. Lower case letters, only small or sparse arborizations. Dash, no arborizations on that side of the VNC. N, neck neuropil; W, wing neuropil; X, accessory metathoracic neuropil (AMN); H, haltere neuropil; iT1, iT2, iT3, intermediate tectulum in the T1, T2 and T3 neuromeres; lT1, lT2, lT3, lower tectulum in the T1, T2 and T3 neuromeres; L1, L2, L3; leg neuropils in the T1, T2 and T3 neuromeres; A, abdominal neuromeres; Ext; external projection towards outside of the VNC. Seg: +, existence of a section of elongated fiber(s) that connect different segregated masses of arborizations. Specific drivers: the lines that label only one or a few types of neurons that are clearly segregated. Other drivers: the lines that also label various other cells. The annotations of innervation in this table do not describe the branching patterns of the neurons, which may also affect how neurons process information. Therefore, to understand the structure of these neurons, this table must be combined with the images of each neuron shown in the three views in the figures, or in the stacks that will be provided via Virtual Fly Brain. *References in Table 1:* *1 Ikeda, K. (1977) Flight motor innervation of a flesh fly. In: Hoyle G. editor. Identified Neurons and Behavior of Arthropods. Springer. pp. 357-358* *2 Koenig, J.H., & Ikeda, K. (1980). Neural interactions controlling timing of flight muscle activity in Drosophila. J. exp. Biol. 87, 121-136.* *3 Harcombe, E.S., & Wyman, R.J. (1977). Output pattern generation by Drosophila flight motoneurons. Journal of Neurophysiology, 40(5), 1066-1077.* *4 Schlurmann, M., & Hausen, K. (2007). Motoneurons of the flight power muscles of the blowfly Calliphora erythrocephala: Structures and mutual dye coupling. The Journal of Comparative Neurology, 500, 448-464.* *5 Duch, C., Bayline, R.J., & Levine, R.B. (2000). Postembryonic development of the dorsal longitudinal flight muscle and its innervation in Manduca sexta. The Journal of Comparative Neurology, 422, 1-17* *6 Duch, C., & Levine, R.B. (2000). Remodeling of membrane properties and dendritic architecture accompanies the postembryonic conversion of a slow into a fast motoneuron. The Journal of Neuroscience, 20(18), 6950-6961.* *7 Kondoh, Y., & Obara, Y. (1982). Anatomy of motoneurones innervating mesothoracic indirect flight muscles in the silkmoth Bombyx mori. J. exp. Biol., 98, 23-37* *8 Trimarchi, J.R., & Schneiderman, A.M. (1995). Flight initiations in Drosophila melanogaster are mediated by several distinct motor patterns. J Comp Physiol A 176, 355-364.* *9 Trimarchi, J. R., & Schneiderman, A. M. (1994). The motor neurons innervating the direct flight muscles of Drosophila melanogaster are morphologically specialized. Journal of Comparative Neurology, 340(3), 427–443. <u>https://doi.org/10.1002/cne.903400311</u>* *10 Strausfeld, N.J., Bassemir, U., Singh, R.N., & Bacon, J.P. (1984). Organization principles of outputs from Dipteran brains. J. Insect. Physiol., 30(1), 73-79.* *11 Fayyazuddin, A., & Dickinson, M.H. (1996). Haltere afferents provide direct, electrotonic input to a steering motor neuron in the blowfly, Calliphora. The Journal of Neuroscience, 16(16), 5225-5232.* *12 von Philipsborn, A. C., Liu, T., Yu, J. Y., Masser, C., Bidaye, S. S., & Dickson, B. J. (2011). Neuronal Control of Drosophila Courtship Song. Neuron, 69(3), 509–522. https://doi.org/10.1016/j.neuron.2011.01.011* *13 Schlurmann, M., & Hausen, K. (2003). Mesothoracic Ventral Unpaired Median (mesVUM) Neurons in the Blowfly Calliphora erythrocephala. Journal of Comparative Neurology, 467(3), 435–453. <u>https://doi.org/10.1002/cne.10930</u>* *14 Kennedy, T., & Broadie, K. (2018). Newly Identified Electrically Coupled Neurons Support Development of the Drosophila Giant Fiber Model Circuit. eNeuro, 5(6), ENEURO.0346-18.2018. https://doi.org/10.1523/ENEURO.0346-18.2018* *15 Trimarchi, J.R., & Murphey, R.K. (1997). The shaking-B^2^ mutation disrupts electrical synapses in a flight circuit in adult Drosophila. J. Neurosci., 17, 4700-4710.* *16 Strausfeld, N.J., & Seyan, H.S. (1985). Convergence of visual, haltere, and prosternal inputs at neck motor neurons of Calliphora erythrocephala. Cell Tissue Res, 240, 601-615.* *17 Hengstenberg, R. (1991). Gaze control in the blowfly Calliphora: a multisensory, two-stage integration process. Seminars in the Neurosciences, 3, 19-29.*

**Table 2:** List of driver lines in the study. For each driver line, the cell categories, cell types, VNC expression quality rating and brain expression quality rating are given. Cell categories provide an overview of which cells are present: Bi, bilateral; Uni, unilateral; MN, motoneuron; LN, local interneuron; IN, intersegmental interneuron; VUM, ventral unpaired median neuron; VPM, ventral paired median neuron. Cell types list the names of cells identified in each driver line. VNC and Brain ratings award a grade of A to driver lines with no off-target expression, A/B for driver lines with off-target expression in one or two cells, B for up to five off-target cell types, C for more than five off-target cell types.

Though we targeted dorsal VNC neurons, our screen serendipitously produced several lines sparsely targeting ventral VNC neurons, including at least one set of leg motoneurons in the mesothoracic segment (T2) of the VNC (SS34789), several primary leg sensory neurons (SS40865, SS41034, SS44081), abdominal motor neurons (SS41029, SS48240, SS48247), and a prosternal sensory neuron (SS47204, SS48287).

Split-GAL4 lines from this collection are available to the public and can be found at splitgal4.janelia.org and driver lines are available at the Bloomington Stock Center or by request.

### Wing motoneurons

To provide tools for analyzing motor circuits and probing motoneuron function, we sought to produce a comprehensive set of driver lines that targeted each of the motoneurons for the wing, the primary appendage driven by dorsal VNC microcircuits. We created 46 driver lines that covered 12/12 of the motoneurons innervating the power muscles (Figures 2-3) and 16/21 of the motoneurons innervating the control muscles (Figures 4-9) with varying levels of specificity.

**Figure 2:**
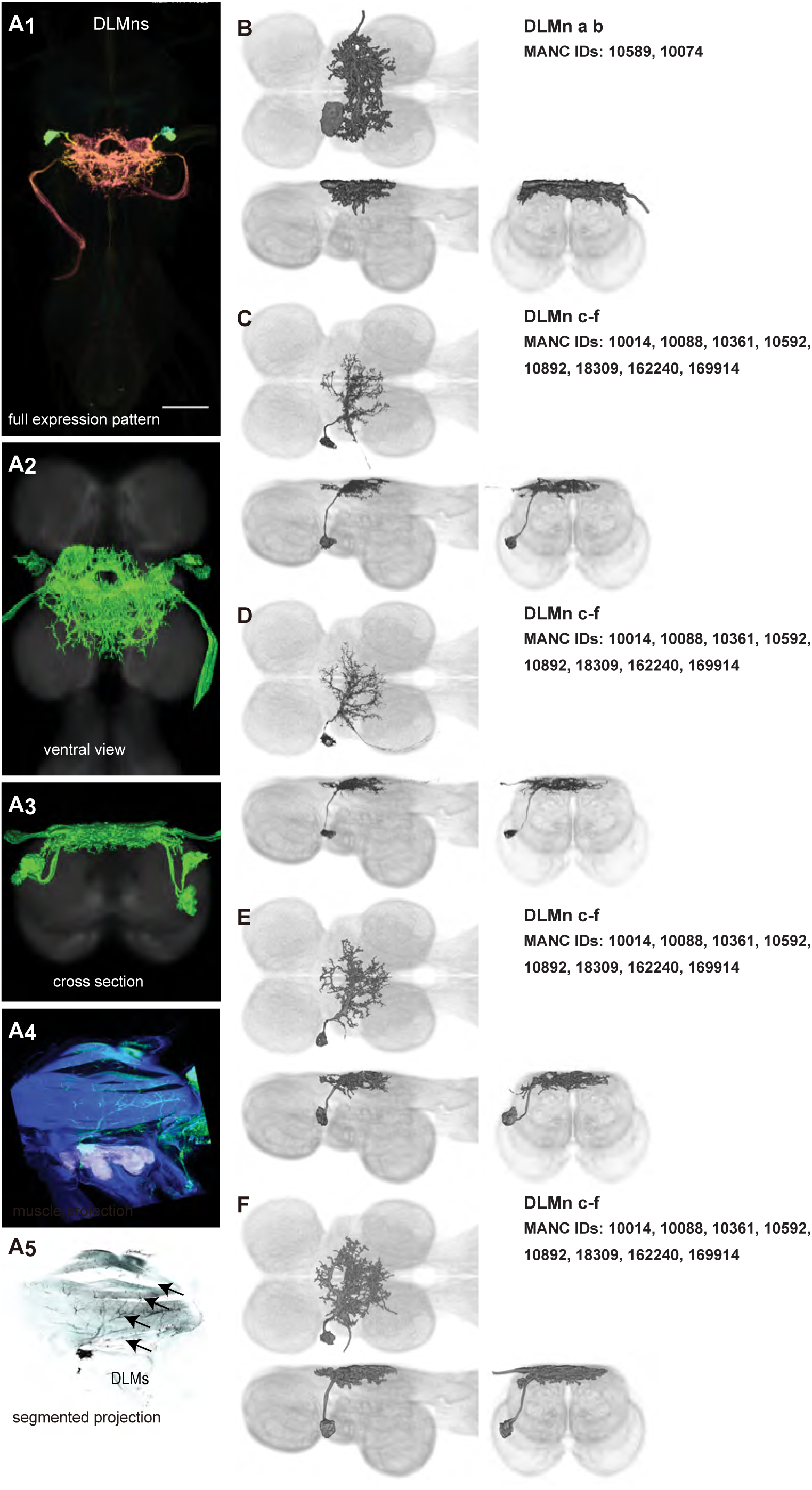
Morphology of DLM power muscle motoneurons. (**A_1_**) Color MIP of full expression pattern of a split line targeting DLMns, SS44039, crossed with pJFRC51-3xUAS-Syt::smGFP-HA in su(Hw)attP1; pJFRC225-5xUAS-IVS-myr::smGFP-FLAG in VK00005, aligned to the JRC 2018 VNC Unisex template. Scale bar in A_1_ represents 50 μm. (**A_2_**) Segmented images of DLM wing motoneurons in upright VNCs. VNCs were aligned to the JRC 2018 Unisex template. (**A_3_**) Transverse views of the segmented neurons shown in A_2_. (**A_4_**) Images of the muscles and their motoneuron innervation in thoraxes stained with phalloidin conjugated with Alexa 633. Phalloidin staining is shown in blue, GFP in green, and nc82 (Bruchpilot, to label the nervous system) in gray. (**A_5_**) Segmented muscle images. (**B-F)** Multicolor flipout (MCFO) was used to separate the power motoneurons, isolating individual cells where possible. Males were used for all motoneuron images.

**Figure 3:**
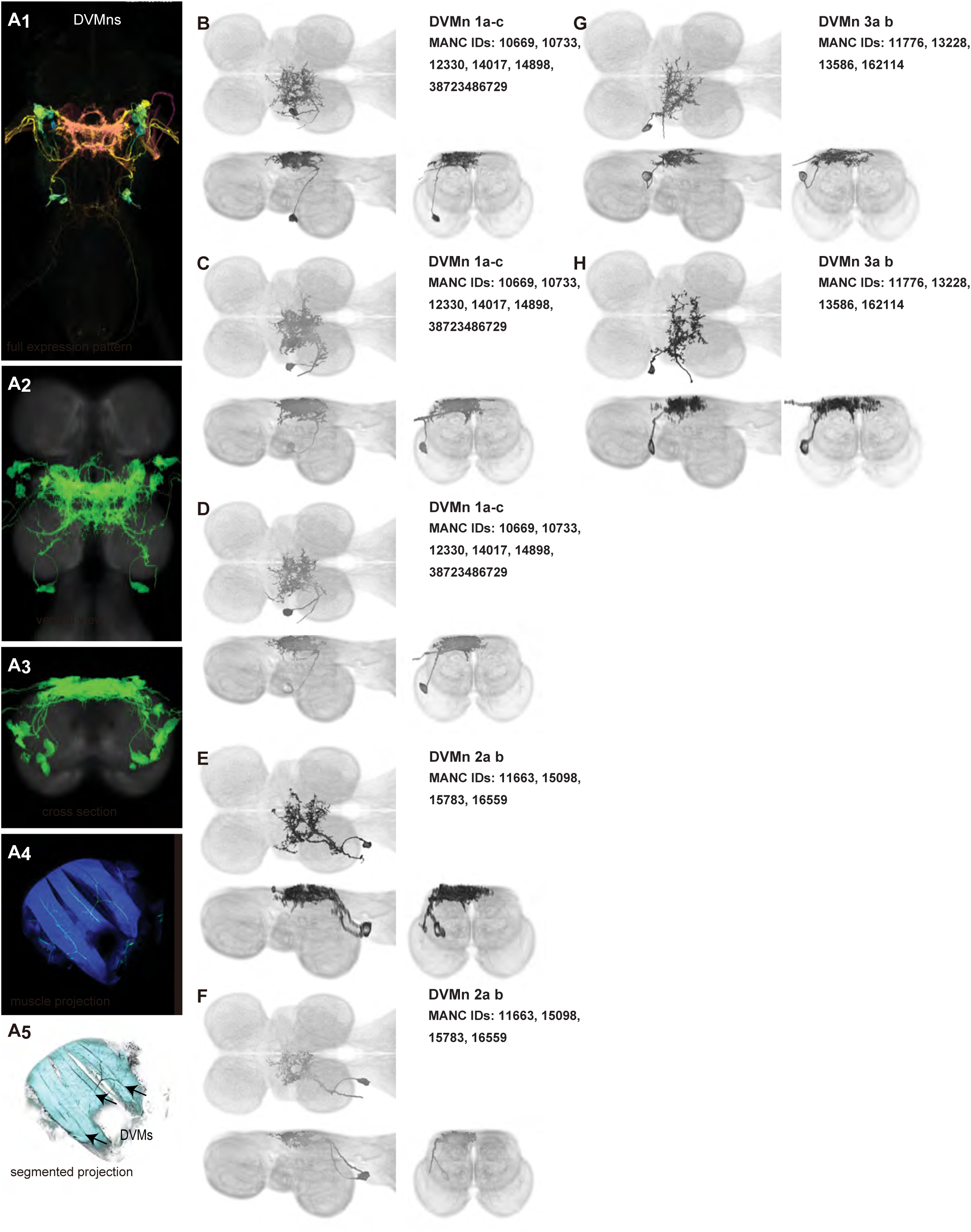
Morphology of DVM power muscle motoneurons. (**A_1_**) Color MIP of full expression pattern of a split line targeting DVMns, SS31950, crossed with pJFRC51-3xUAS-Syt::smGFP-HA in su(Hw)attP1; pJFRC225-5xUAS-IVS-myr::smGFP-FLAG in VK00005, aligned to the JRC 2018 VNC Unisex template. Scale bar in A_1_ represents 50 μm. (**A_2_**) Segmented images of DVM wing motoneurons in upright VNCs. VNCs were aligned to the JFC 2018 Unisex template. (**A_3_**) Transverse views of the segmented neurons shown in A_2_. (**A_4_**) Images of the muscles and their motoneuron innervation in thoraxes stained with phalloidin conjugated with Alexa 633. Phalloidin staining is shown in blue, GFP in green, and nc82 (Bruchpilot, to label the nervous system) in gray. (**A_5_**) Segmented muscle images. (**B-H)** Multicolor flipout (MCFO) was used to separate the power motoneurons, isolating individual cells where possible. Males were used for all motoneuron images.

**Figure 4:**
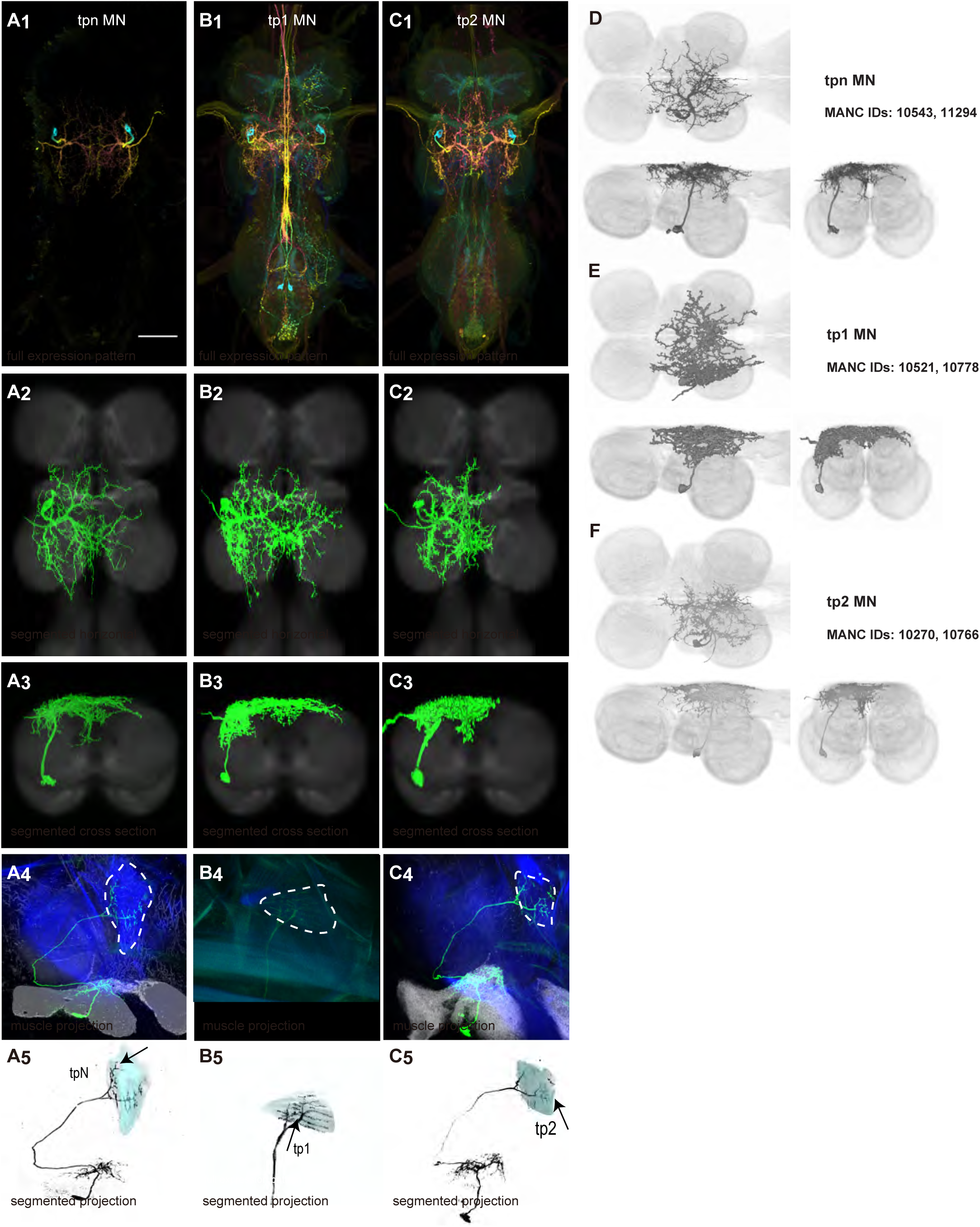
Morphology of tergopleural wing steering muscle motoneurons targeted by our sparse split lines. (**A-C_1_**) Color MIPs of full expression patterns of the split lines (respectively SS51528, SS41052, SS47120), crossed with pJFRC51-3xUAS-Syt::smGFP-HA in su(Hw)attP1; pJFRC225-5xUAS-IVS-myr::smGFP-FLAG in VK00005, aligned to the JRC 2018 VNC Unisex template. Scale bar in A_1_ represents 50 μm. (**A-C_2_**) Segmented images of tergopleural wing motoneurons in upright VNCs. Multicolor flipout (MCFO) was used to separate left and right neurons. VNCs were aligned to the JFC 2018 Unisex template. (**A-C_3_**) Transverse views of the segmented neurons shown in A-C_2_. (**A-C_4_**) Images of the muscles and their motoneuron innervation in thoraxes stained with phalloidin conjugated with Alexa 633. Phalloidin staining is shown in blue, GFP in green, and nc82 (Bruchpilot, to label the nervous system) in gray. (**A-C_5_**) Segmented muscle images. (**D-F**) segmented images of steering motoneurons including side views. Males were used for all motoneuron images except tpN, as our tpN line lacked expression in males.

#### Power muscle motoneurons

We generated 16 driver lines targeting the two classes of motoneurons that each innervate a wing power muscle type: dorsal longitudinal motoneurons (DLMns) and dorsoventral motoneurons (DVMns), which innervate the dorsal longitudinal muscles (DLMs) and dorsoventral muscles (DVMs), respectively. The morphology and innervation patterns of these power motoneurons are shown in Figures 2-3. The DLMns (Figure 2) comprise five motoneurons, with DLMna/b innervating the two dorsal DLM fibers and DLMnc-f innervating the other four DLM fibers (Coggshall, 1978). The DVMns (Figure 3) comprise seven motoneurons, each innervating a single DVM fiber, labeled DVMn1a-c, DVMn2a-b, and DVMn3a-b (Schlurmann and Hausen, 2007, Trimarchi and Schneiderman, 1995).

These motoneuron lines are varyingly selective for power muscle type, and can be categorized as being either 1) solely selective for DLMns (no strong DVMn expression: SS31541, SS31561, SS44039, and SS44056), 2) largely selective for DVMns (expression in one or more DLMns: SS41068 and SS49797), or 3) exhibiting mixed expression in both DLMns and DVMns (SS31543, SS31950, SS31997, SS37294, SS37295, SS40765, SS40772, SS40989, SS43980, SS44060). To confirm the identity of each set of motoneurons, we determined which muscles they innervated by co-staining the expression pattern of each split-GAL4 line with GFP and the muscles with phalloidin (Figures 2A, 3A). In separate experiments, we used MCFO to isolate individual power muscle motoneurons within our split-GAL4 lines and manually segment them where possible, creating digital meshes of individual neurons (or, where they could not be separated, bilateral pairs) to better characterize their morphology within the central nervous system (Figures 2B-F, 3B-H).

DLMnc-f are nearly identical and their somata form a closely apposed cluster (Figure 2C-F). The DLMns could not be separated using split-GAL4 driver lines; every line with any DLMn expression had expression in all DLMns. Due to the nature of our technique, muscle images and high-quality VNC images could not be obtained from the same MCFO preparation. Thus, we were unable to identify which DLMn neuron in our MCFO data corresponded to which specific DLM fiber and we therefore refer to these as a group, DLMnc-f. In contrast, DLMna/b, which innervates two DLM fibers, was definitively identified based on its morphology within the VNC, including its sizeable dorsal soma and contralateral axon (Figure 2B). The DLMns arborize extensively in both ipsi- and contralateral wing neuropils, where they receive all their inputs (Table 1).

Among the DVMns (Figure 3), soma positions are comparatively unique and their correspondence to DVM innervated has been established (Schlurmann and Hausen, 2007). The muscle DVM 1 is innervated by three motoneurons (DVMn1a-c, Figure 3B-D), whose somata are connected to their dendritic arbors by a neurite that projects posteriorly away from the soma before bending back to the anterior wing neuropil. The muscle DVM 2 is innervated by two motoneurons with posterior somata (DVMn2a-b, Figure 3E-F). In our MCFO preparations, DVMn2a never had expression on its own, so it could not be segmented and appears in the same image as DVMn1b (Figure 3E). The muscle DVM 3 is innervated by two motoneurons with anterior somata (DVMn3a-b, Figure 3G-H). Relative to the DLMns, the DVMns exhibit less contralateral arborization, and they extend ventral projections into the intermediate tectulum (Figure 3—figure supplement 1).

#### Control muscle motoneurons

We generated 35 driver lines that targeted the wing control motoneurons. 19 of these driver lines are expressed in a single motoneuron cell type (Figures 4-9). These single neuron driver lines included motoneurons to both indirect control muscles (tp1, tp2, tpN, ps1, tt; Figures 4 and 5) and direct steering muscles (i1, i2, hg1, hg2, hg3; Figures 7 and 9). The remaining 16 driver lines expressed in more than one motoneuron type but collectively covered 16/21 wing control motoneurons. Some MNs in these multiple-MN driver lines could be identified and segmented (b1, b2, b3, iii1, iii3, iii4; Figures 6 and 8). As with the power muscles, we determined the identities of these motoneurons by co-staining for both driver line expression and muscle tissue (e.g. Figure 4A) and isolated individual motoneurons using MCFO (e.g. Figure 4D). The steering motoneurons covered in our collection innervate a mixture of both tonic (b1, b3, i2, iii3) and phasic (b2, i1, iii1, iii4, hg1, hg2, hg3) muscles, the two functionally distinct categories of direct steering muscles (Lindsay et al., 2017).

**Figure 5:**
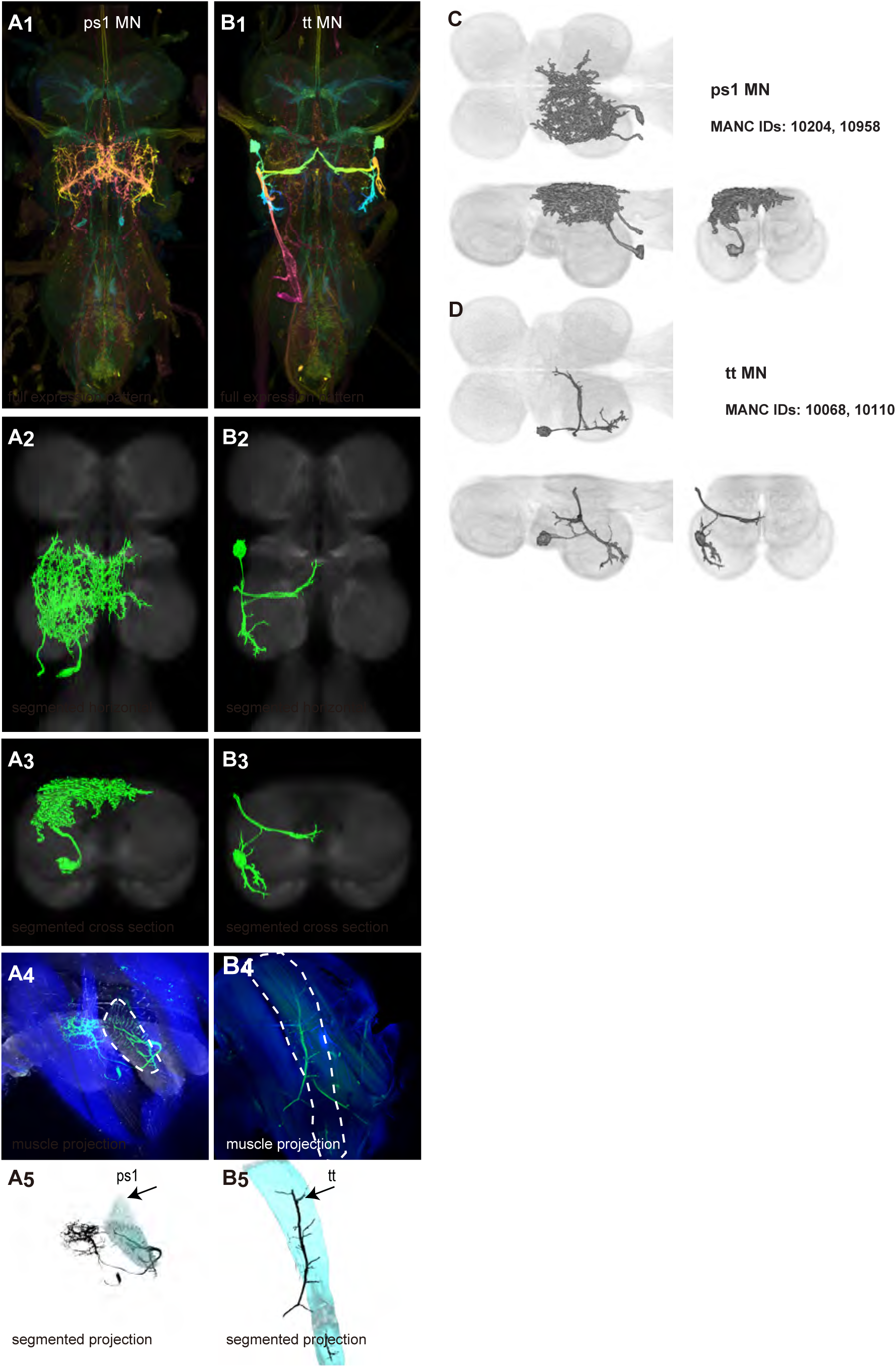
Morphology of other indirect wing steering muscle motoneurons targeted by our sparse split lines. (**A-B_1_**) Color MIPs of full expression patterns of the split lines (SS47204, SS47125), crossed with pJFRC51-3xUAS-Syt::smGFP-HA in su(Hw)attP1; pJFRC225-5xUAS-IVS-myr::smGFP-FLAG in VK00005, aligned to the JRC 2018 VNC Unisex template. Scale bar in A_1_ represents 50 μm. (**A-B_2_**) Segmented images of other indirect wing motoneurons in upright VNCs. Multicolor flipout (MCFO) was used to separate left and right neurons. VNCs were aligned to the JFC 2018 Unisex template. (**A-B_3_**) Transverse views of the segmented neurons shown in A-B_2_. (**A-B_4_**) Images of the muscles and their motoneuron innervation in thoraxes stained with phalloidin conjugated with Alexa 633. Phalloidin staining is shown in blue, GFP in green, and nc82 (Bruchpilot, to label the nervous system) in gray. (**A-B_5_**) Segmented muscle images. (**C-D**) segmented images of steering motoneurons including side views. Males were used for all motoneuron images.

**Figure 6:**
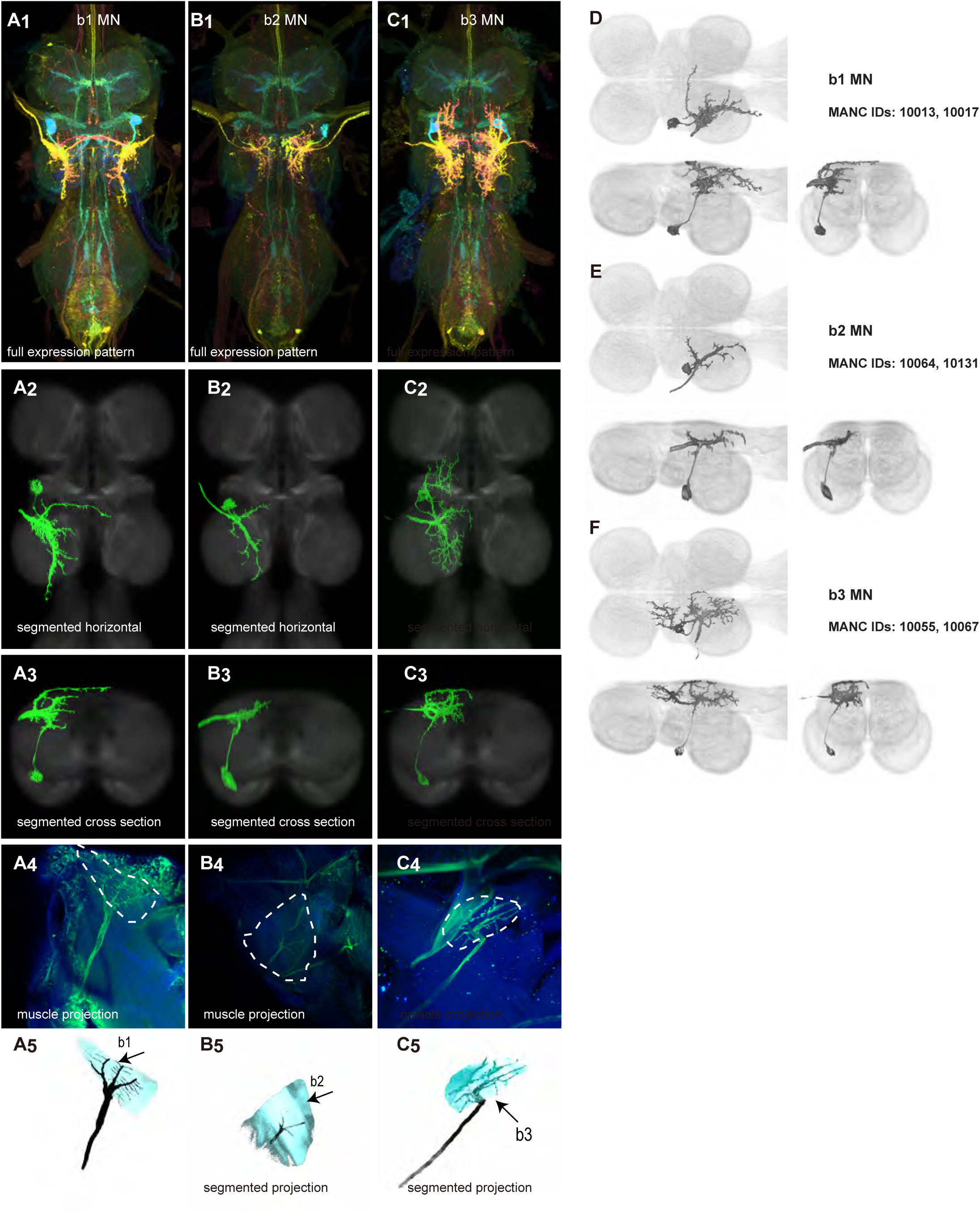
Morphology of basalar wing steering muscle motoneurons targeted by our sparse split lines. (**A-B_1_**) Color MIPs of full expression patterns of the split lines (SS40980, SS45772, SS45779), crossed with pJFRC51-3xUAS-Syt::smGFP-HA in su(Hw)attP1; pJFRC225-5xUAS-IVS-myr::smGFP-FLAG in VK00005, aligned to the JRC 2018 VNC Unisex template. Scale bar in A_1_ represents 50 μm. (**A-B_2_**) Segmented images of basalar wing motoneurons in upright VNCs. Multicolor flipout (MCFO) was used to separate left and right neurons. VNCs were aligned to the JFC 2018 Unisex template. (**A-B_3_**) Transverse views of the segmented neurons shown in A-B_2_. (**A-B_4_**) Images of the muscles and their motoneuron innervation in thoraxes stained with phalloidin conjugated with Alexa 633. Phalloidin staining is shown in blue, GFP in green, and nc82 (Bruchpilot, to label the nervous system) in gray. (**A-B_5_**) Segmented muscle images. (**C-D**) segmented images of steering motoneurons including side views. Males were used for all motoneuron images.

**Figure 7:**
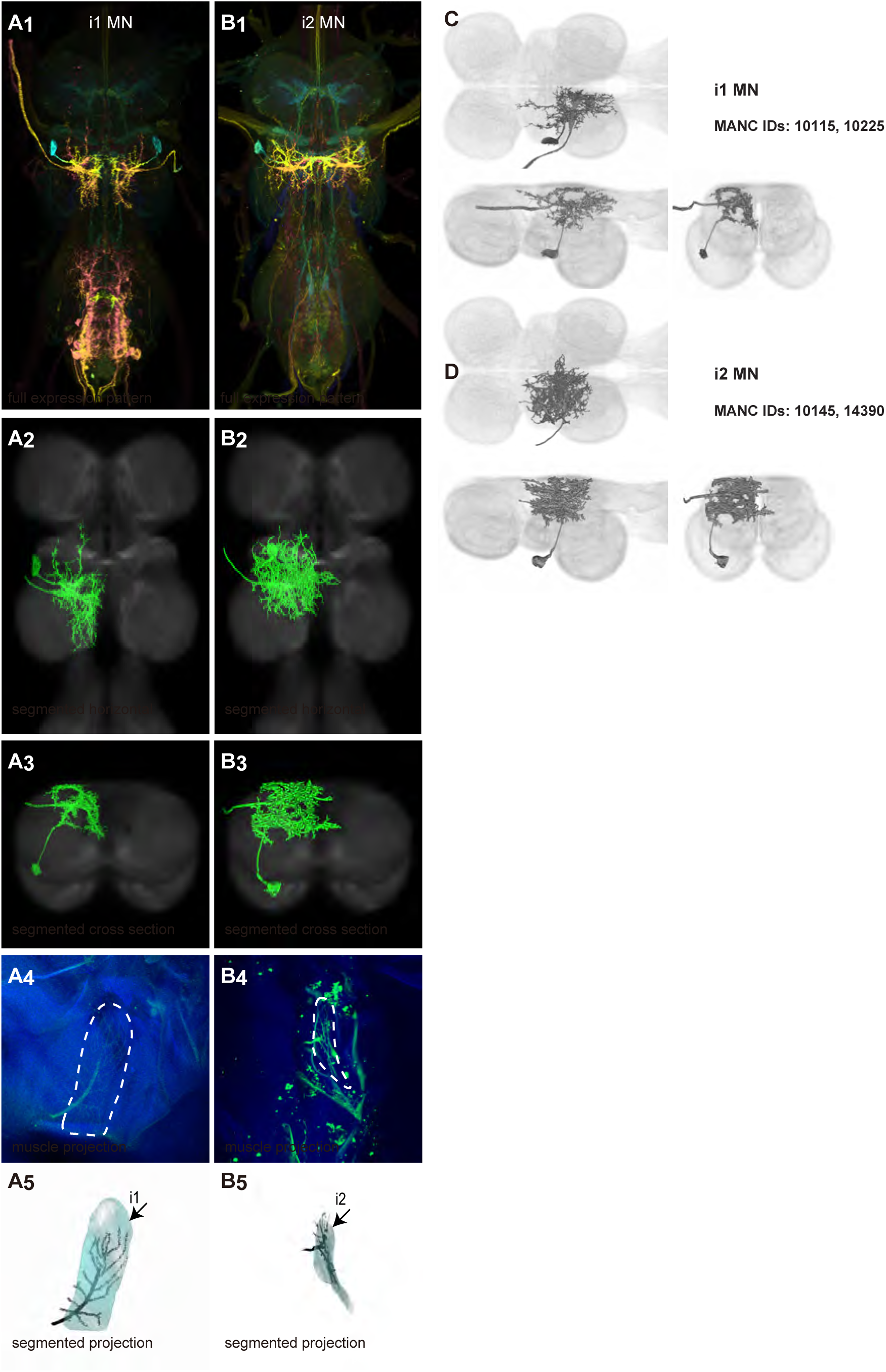
Morphology of first axillary wing steering muscle motoneurons targeted by our sparse split lines. (**A-B_1_**) Color MIPs of full expression patterns of the split lines (SS41039, SS45782), crossed with pJFRC51-3xUAS-Syt::smGFP-HA in su(Hw)attP1; pJFRC225-5xUAS-IVS-myr::smGFP-FLAG in VK00005, aligned to the JRC 2018 VNC Unisex template. Scale bar in A_1_ represents 50 μm. (**A-B_2_**) Segmented images of first axillary wing motoneurons in upright VNCs. Multicolor flipout (MCFO) was used to separate left and right neurons. VNCs were aligned to the JFC 2018 Unisex template. (**A-B_3_**) Transverse views of the segmented neurons shown in A-B_2_. (**A-B_4_**) Images of the muscles and their motoneuron innervation in thoraxes stained with phalloidin conjugated with Alexa 633. Phalloidin staining is shown in blue, GFP in green, and nc82 (Bruchpilot, to label the nervous system) in gray. (**A-B_5_**) Segmented muscle images. (**C-D**) segmented images of steering motoneurons including side views. Males were used for all motoneuron images.

**Figure 8:**
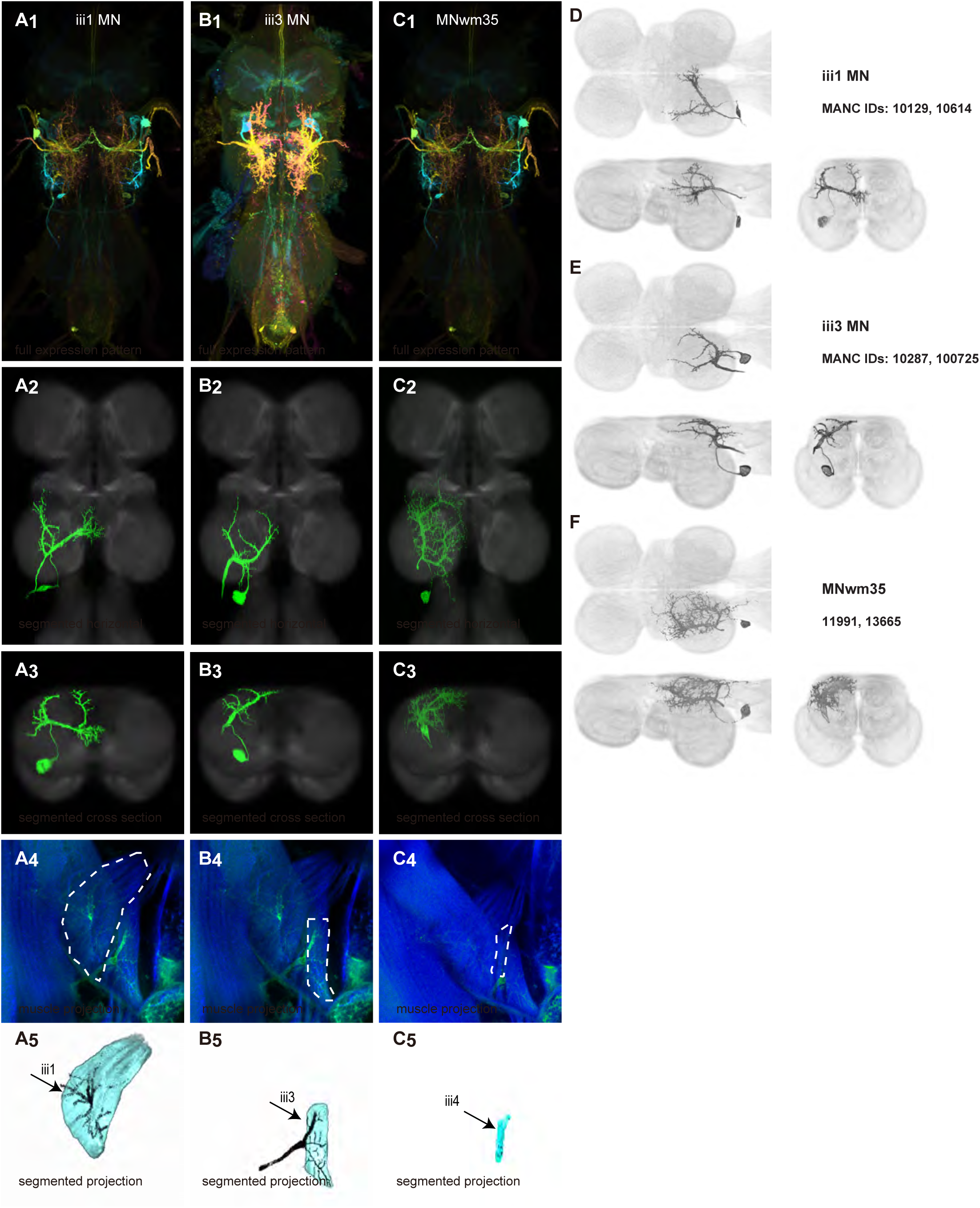
Morphology of third axillary wing steering muscle motoneurons targeted by our sparse split lines. (**A-B_1_**) Color MIPs of full expression patterns of the split lines (SS41027, SS45779, SS41027), crossed with pJFRC51-3xUAS-Syt::smGFP-HA in su(Hw)attP1; pJFRC225-5xUAS-IVS-myr::smGFP-FLAG in VK00005, aligned to the JRC 2018 VNC Unisex template. Scale bar in A_1_ represents 50 μm. (**A-B_2_**) Segmented images of third axillary wing motoneurons in upright VNCs. Multicolor flipout (MCFO) was used to separate left and right neurons. VNCs were aligned to the JFC 2018 Unisex template. (**A-B_3_**) Transverse views of the segmented neurons shown in A-B_2_. (**A-B_4_**) Images of the muscles and their motoneuron innervation in thoraxes stained with phalloidin conjugated with Alexa 633. Phalloidin staining is shown in blue, GFP in green, and nc82 (Bruchpilot, to label the nervous system) in gray. (**A-B_5_**) Segmented muscle images. (**C-D**) segmented images of steering motoneurons including side views. Males were used for all motoneuron images.

The control motoneurons mainly have their dendrites in the wing neuropil ipsilateral to their soma and axon (Figure 9—figure supplement 1). Unlike the power MNs, which have almost all their dendrites in the anterior wing neuropil, some control MNs also receive input in the posterior wing neuropil, intermediate or lower tectulum, neck neuropil and haltere neuropil. Morphologically, the control MNs can be classified into unilateral neurons that mainly arborize in the wing neuropil (b1, iii3, and iii4 MNs; Figure 9—figure supplement 1A), bilateral neurons that mainly arborize in the wing neuropil (tp1, tp2, tpN, ps1, hg2, and hg3 MNs; Figure 9— figure supplement 1B), neurons that arborize approximately equally in the wing neuropil and intermediate tectulum (b3, i1, i2, and hg1 MNs; Figure 9—figure supplement 1C), and neurons that have more innervation in the intermediate and lower tectulum than in the wing neuropil (tt, b2, and iii1 MNs; Figure 9—figure supplement 1D). The tergotrochanter muscle motoneuron (tt MN) was unique among control motoneurons in that it had no wing neuropil innervation. Previous studies have demonstrated that this motoneuron is primarily used during jumping, and is responsible for extending the middle legs during escape(Trimarchi and Schneiderman, 1993). Similar to the tt MN, the b2 and iii1 MNs also had some dense arborization in the tectulum and less arborization in the wing neuropil.

**Figure 9:**
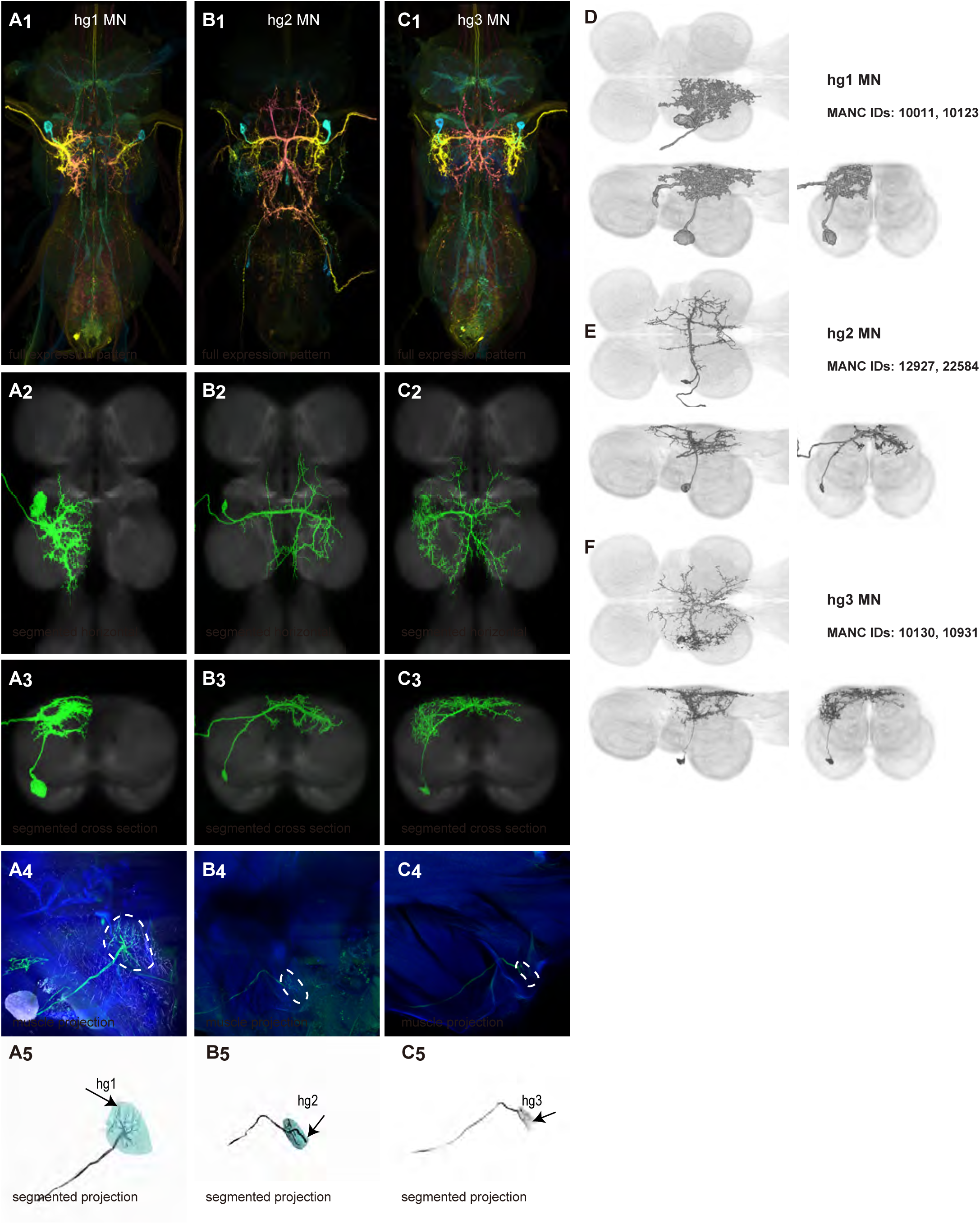
Morphology of fourth axillary wing steering muscle motoneurons targeted by our sparse split lines. (**A-C_1_**) Color MIPs of full expression patterns of the split lines (SS32023,SS37253, SS49039), crossed with pJFRC51-3xUAS-Syt::smGFP-HA in su(Hw)attP1; pJFRC225-5xUAS-IVS-myr::smGFP-FLAG in VK00005, aligned to the JRC 2018 VNC Unisex template. Scale bar in A_1_ represents 50 μm. (**A-C_2_**) Segmented images of fourth axillary wing motoneurons in upright VNCs. Multicolor flipout (MCFO) was used to separate left and right neurons. VNCs were aligned to the JFC 2018 Unisex template. (**A-C_3_**) Transverse views of the segmented neurons shown in A-C_2_. (**A-C_4_**) Images of the muscles and their motoneuron innervation in thoraxes stained with phalloidin conjugated with Alexa 633. Phalloidin staining is shown in blue, GFP in green, and nc82 (Bruchpilot, to label the nervous system) in gray. (**A-C_5_**) Segmented muscle images. (**D-F**) segmented images of steering motoneurons including side views. Males were used for all motoneuron image.

### Functional experiments using wing motoneuron split lines in tethered flight and song

To demonstrate the utility of our driver lines and to provide a functional baseline for future analyses of premotor circuits, we used the motoneuron driver lines described above to probe the roles of both power and control wing muscles in the execution of two distinct wing behaviors: flight and courtship song. Previous work suggests that these behaviors are coordinated by overlapping sets of muscles (Heide and Götz, 1996, Lindsay et al., 2017, O’Sullivan et al., 2018). Our collection of driver lines and quantitative behavioral assays allow us to confirm and expand upon these previous results to further understand how wing motoneurons participate in flight and song.

To complement existing data, we used our most sparse-expressing motoneuron lines to quantify the phenotypes of single muscle activation during flight or inactivation in courtship. Because the motoneuron-to-muscle relationship is one-to-one for the motoneurons we tested, we interpret our results as activating or silencing individual muscles. In flight we evaluated the roles of eight muscles, including the two different sets of power muscles (DLM, DVM) and six control muscles (tp2, ps1, i1, i2, hg1, hg2). In courtship we studied a subset of wing motoneurons (DVM, tp2, ps1, i2, hg1, hg2, hg3). As a genetic control, we tested flies using “empty” split-GAL4 lines with the same genetic background but minimal GAL4 expression (SS01062, for flight studies and SS01062 and SS01055 for courtship) (Namiki et al., 2018).

Previous studies indicated that activity in specific wing control muscles acts to make small changes in wing kinematics that have large aerodynamic consequences for controlling the fly’s flight trajectory (Lindsay et al., 2017, Whitehead et al., 2022, Melis et al., 2024). Here we used our lines to drive expression of a red-shifted channelrhodopsin (CsChrimson; (Klapoetke et al., 2014)) in single motoneurons, allowing us to optogenetically activate specific motoneurons, and hence wing muscles, during tethered flight and quantify what effect this had on the details of wing kinematics. Using an automated video tracking algorithm (“Kinefly,” see Methods) we quantified the activation-induced changes in four wing kinematic parameters: wingbeat frequency, back and forward deviation of the wings, and stroke amplitude (Figure 10A-C).

**Figure 10:**
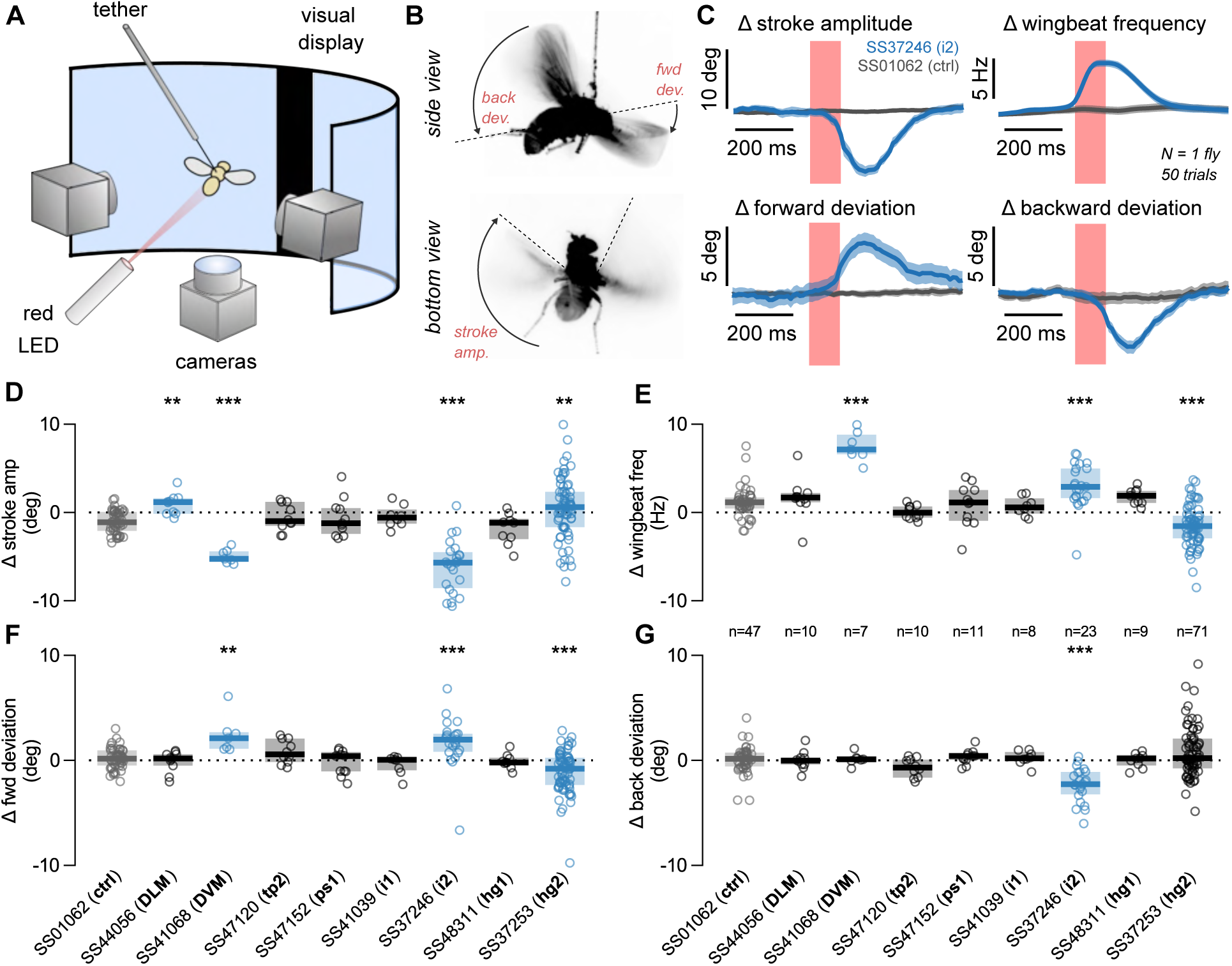
Optogenetic stimulation of wing motor neurons evokes changes to flight kinematics. (**A**) Schematic of tethered flight measurement apparatus. Tethered flies are positioned in front of a display screen that presents both open- and closed-loop visual stimuli. The fly is illuminated near-infrared LEDs and filmed by three cameras recording at 100 fps. An additional red LED (617 nm) provides optogenetic stimulation in 100 ms pulses. Stills from two of the three cameras recording the tethered fly in flight. Top panel (side view) illustrates the forward (fwd) and backward (back) deviation angles; bottom panel (bottom view) illustrates the stroke amplitude. (B) Averaged wing kinematics for two flies undergoing 50 repetitions of a closed loop trial with 100 ms optogenetic pulse. Dark lines and envelopes show the mean and 95% confidence interval, respectively. Traces in blue correspond to a i2-GAL4>UAS-CsChrimson fly; traces in gray show an example genetic control, SS01062>UAS-CsChrimson, where SS01062 is an empty split Gal4 line. (**D-G**) Statistics across flies for the four kinematic variables shown in (C): stroke amplitude (D), wingbeat frequency (E), forward deviation angle (F), and backward deviation angle (G). Open circles show per-fly measurements; bars and horizontal lines show interquartile range and population median, respectively. The number of flies per genotype is shown between E and G. Significance is determined via Wilcoxon rank sum test with Bonferroni correction (***, p<0.001; **, p<0.01; *, p<0.05).

We found that activation of one type of power muscle, the DLMs, had a relatively minor effect on the wingstroke: we measured a small, but significant, increase in stroke amplitude, with no other significant changes to wing kinematic parameters (Figure 10D-G, p<0.05, Wilcoxon rank sum test compared to control genotype). In contrast, activation of the other power muscle type, the DVMs, generated a large behavioral response, decreasing stroke amplitude and increasing both forward deviation angle and wingbeat frequency (Figure 10D-G). The activation of the control muscle i2 had the strongest effect in our screen, with a phenotype similar to DVM activation with the addition of a decrease in backward wing deviation (Figure 10D-G). Activation of hg2 produced changes in three of our four wing kinematic parameters, but the mean value of the parameters changed very little; rather, the variance we observed between flies increased substantially relative to genetic control flies (Figure 10D-G). Activation of the motoneurons for the remaining muscles (tp2, ps1, i1, hg1) did not lead to any observable phenotypes. This was a surprising result for tp2, which has previously been implicated as critical for flight and correct wing posture (O’Sullivan et al., 2018). One interpretation of our result is that, to engage the correct wing posture during flight, the tp2 muscle is maximally active and hence further motoneuron activation does not alter wing kinematics.

To investigate the role of wing muscles in courtship song, we silenced individual motoneurons in male flies using our motoneuron-specific lines to drive either the inwardly-rectifying potassium channel Kir2.1 (UAS-Kir2.1) or tetanus neurotoxin light chain (UAS-TNTe) to hyperpolarize the neurons or block synaptic transmission, respectively (Baines et al., 2001, Sweeney et al., 1995). Motoneuron-silenced males were individually housed before introduction into a recording chamber with a 1-day old virgin female, where we recorded 30 minutes of courtship song (Figure 11A) (Arthur et al., 2013). Because experiments with both silencing reporters produced similar results, Figure 11 and Figure 11—figure supplement 1 show data from only Kir2.1 flies.

**Figure 11:**
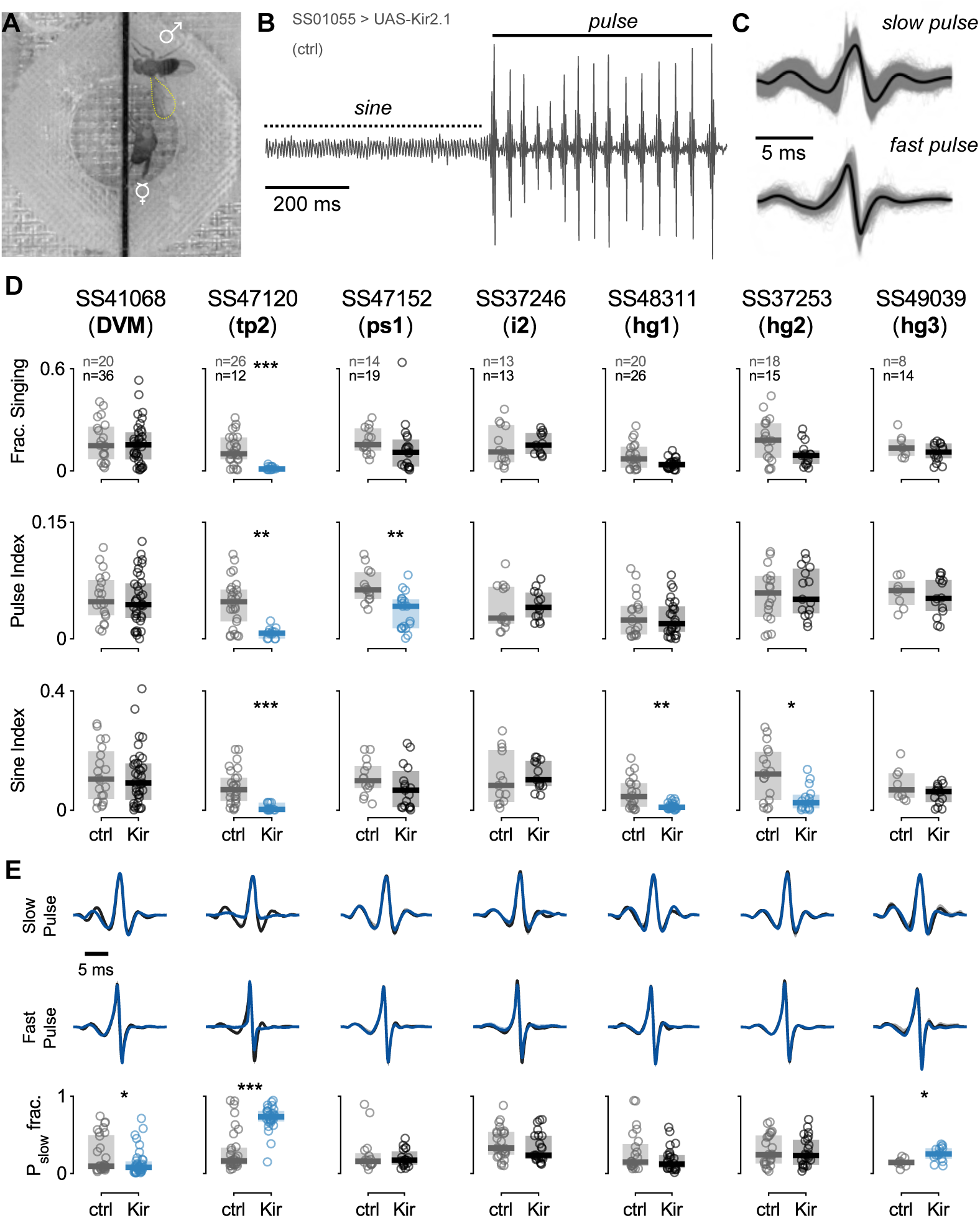
Chronic silencing of wing motoneurons results in courtship song deficits. (**A**) Image from a typical courtship assay, showing a male (above) extending its left wing to sing to a female (below). (**B**) Example trace of song recording from a control group fly (SS01055>UAS-Kir2.1). Bouts of sine and pulse song are labelled on the left and right of the trace, respectively. (**C**) Example slow (top) and fast (bottom) pulse modes for a single fly. Thick black lines show the mean pulse shape; thin gray lines show individual pulses. (**D**) Courtship song statistics for motoneuron driver lines crossed to UAS-Kir2.1. Rows show different song parameters: total fraction of the trial spent singing (top), fraction of song spent singing pulse mode song (middle), and fraction of song spent singing sine mode song (bottom). Open circles show per-fly measurements; bars and horizontal lines show interquartile range and population median, respectively. The number of flies per genotype is given in D, with the sample sizes for control and experimental groups on the top and bottom, respectively. (**E**)€ Analysis of pulse type. The top and middle and rows show waveforms for slow and fast pulse modes, respectively, for each genotype in C (blue), overlaid onto control (dark gray). Thick line shows the grand mean across flies; envelope gives 95% confidence interval for mean from bootstrap. Bottom row shows the fraction of pulses that are classified as slow for each genotype. Significance assigned using the Wilcoxon rank sum test and Fisher’s exact test (***, p<0.001; **, p<0.01; *, p<0.05).

The song of *Drosophila melanogaster* contains two primary components: sine song and pulse song (Figure 11B). We analyzed the fraction of time during the assay in which flies within a given genotype produced song (Figure 11D, top), as well as the fraction of the song devoted to sine and pulse song, denoted sine index and pulse index, respectively (Figure 11D, middle and bottom). We found that silencing tp2 affected nearly all aspects of song production, silencing ps1 decreased production of pulse song, and silencing hg1 and hg2 decreased production of sine song (*p*<0.05, Figure 11D), consistent with previous findings (Shirangi et al., 2013, O’Sullivan et al., 2018).

To expand upon these and previous results, we further analyzed the details of pulse song production. Recent studies uncovered two distinct categories of pulses: fast and slow (Figure 11C) (Clemens et al., 2018). Figure 11E shows averaged slow and fast pulse shapes for each genotype (blue, top and middle) in our experiments compared to control flies (gray, top and middle). Across genotypes, we compared the relative proportion of slow pulse types among the total number of pulses produced and found that DVM-silenced flies produced significantly fewer slow-mode pulses, while hg3-silenced flies produced significantly more slow-mode pulses (*p*<0.05; Figure 11E). Flies with the tp2 motoneuron silenced showed a significant increase in fast pulses produced, but this may be an epiphenomenon related to the low number of total pulses produced by these flies. Additional quantified song traits, including inter-pulse interval, sine song frequency, and transition probabilities between song types, were all consistent with the findings above; namely, tp2-silenced flies exhibited differences for both song types, hg1 and hg2 flies exhibited specific differences in sine song frequency, and ps1 flies exhibited differences in pulse song timing (Figure 11—figure supplement 1A). Notably, i2-silenced flies also showed a statistically significant decrease in inter-pulse interval; however, further testing would be required to firmly establish this result.

### Haltere motoneurons

Each haltere—a mechanosensory organ derived from the ancestral hindwing (Figure 1A)—is associated with its own power muscle, hDVM, and six steering muscles (Dickerson et al., 2019). We generated a total of 15 driver lines that targeted the haltere motoneurons (Figures 12 and 13). Three of the haltere motoneurons could be both segmented and unambiguously identified: hDVM, hi1, and hi2 (Figure 12). We also identified the hb1 and hb2 motoneurons in driver line SS47195, along with hi1 MN (Figure 13B). We segmented these cells using MCFO (Figure 13D,E), but were unable to determine which cell innervated hb1 and which innervated hb2. We thus named the cells hb1/2a and hb1/2b. SS36076 had expression in hiii3 MN, but we were unable to segment any image of this motoneuron. Collectively, these haltere motoneurons fall into two broad morphological categories: unilateral intersegmental cells that arborize in the haltere, wing, and neck neuropils (hb1/2 MNs; Figure 13—figure supplement 1A) and bilateral cells that largely arborize in the haltere neuropil (hDVMn, hi1/2 MNs; Figure 13—figure supplement 1B).

**Figure 12:**
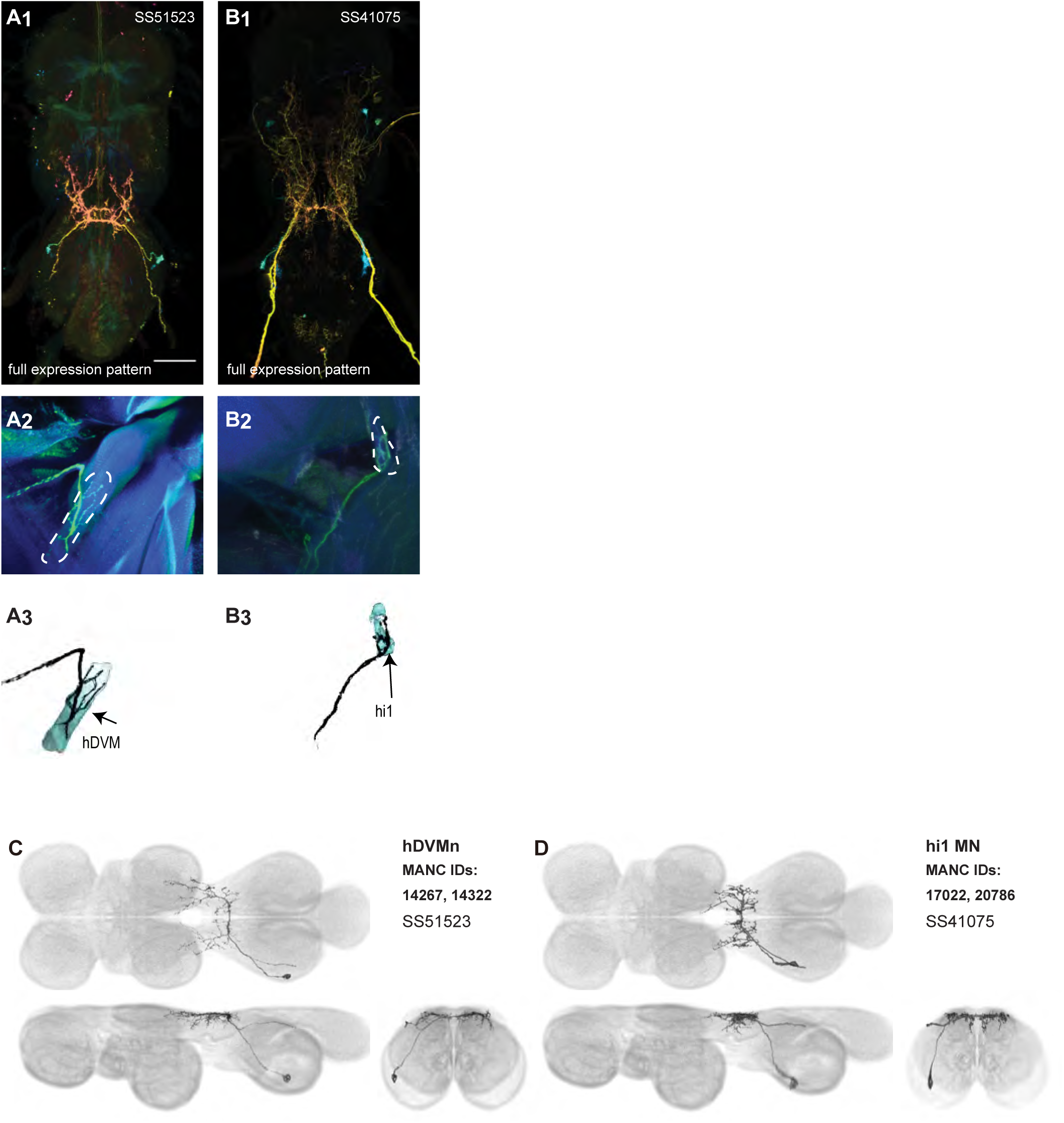
Morphology of haltere motoneurons targeted by our sparse split lines. (**A-B_1_**) Color MIPs of full expression patterns of the split lines (SS51523, SS41075), crossed with pJFRC51-3xUAS-Syt::smGFP-HA in su(Hw)attP1; pJFRC225-5xUAS-IVS-myr::smGFP-FLAG in VK00005, aligned to the JRC 2018 VNC Unisex template. Scale bar in A_1_ represents 50 μm. (**A-B_2_**) Images of the muscles and their motoneuron innervation in thoraxes stained with phalloidin conjugated with Alexa 633. Phalloidin staining is shown in blue, GFP in green, and nc82 (Bruchpilot, to label the nervous system) in gray. (**A-B_3_**) Segmented muscle images. (**C-D**) segmented images of haltere motoneurons.

**Figure 13:**
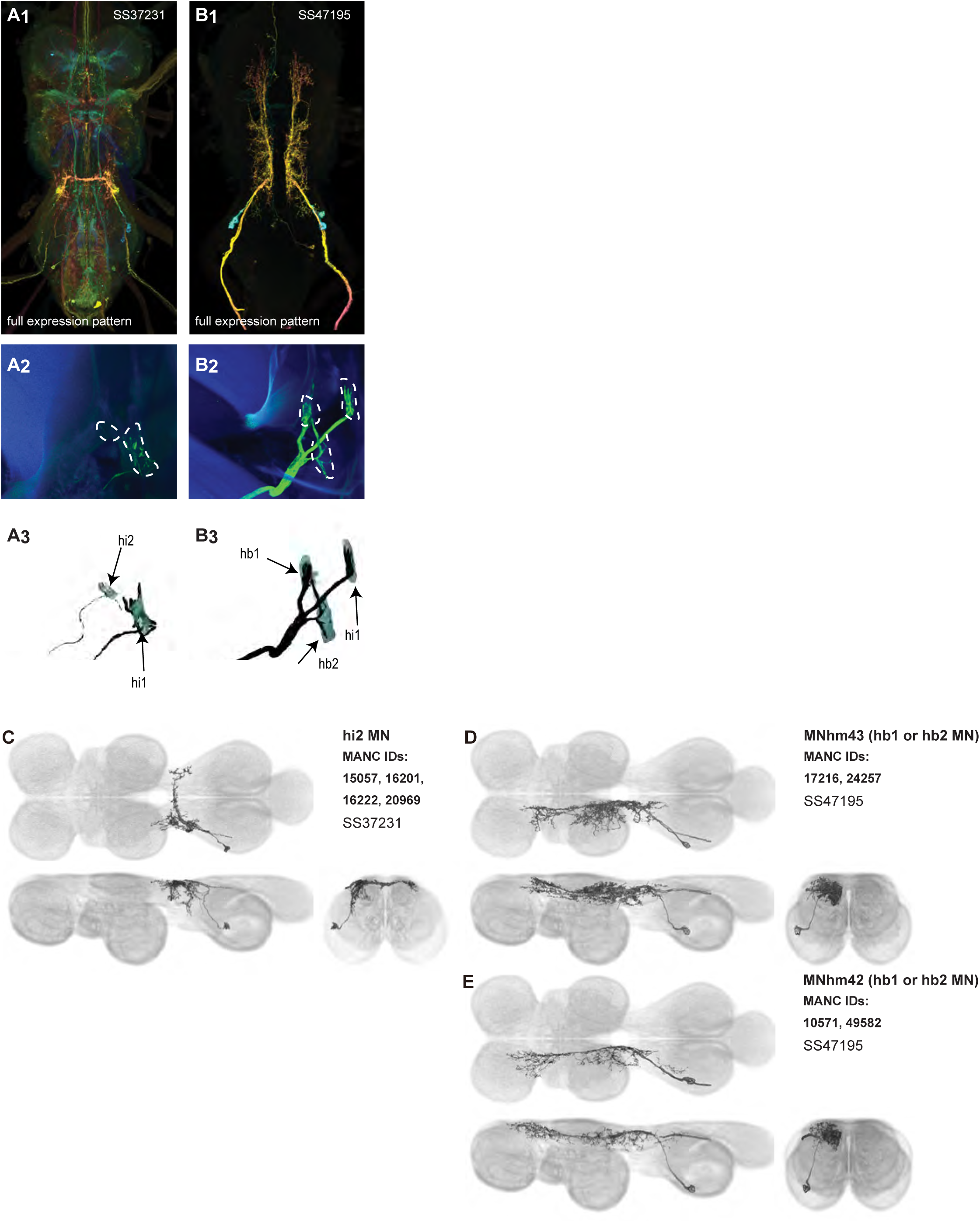
Morphology of haltere motoneurons targeted by our broad split lines. (**A-B_1_**) Color MIPs of full expression patterns of the split lines (SS37231, SS47195), crossed with pJFRC51-3xUAS-Syt::smGFP-HA in su(Hw)attP1; pJFRC225-5xUAS-IVS-myr::smGFP-FLAG in VK00005, aligned to the JRC 2018 VNC Unisex template. Scale bar in A_1_ represents 50 μm. (**A-B_2_**) Images of the muscles and their motoneuron innervation in thoraxes stained with phalloidin conjugated with Alexa 633. Phalloidin staining is shown in blue, GFP in green, and nc82 (Bruchpilot, to label the nervous system) in gray. (**A-B_3_**) Segmented muscle images. (**C-E**) segmented images of haltere motoneurons.

### VUM neurons

We created four driver lines targeting mesothoracic ventral unpaired median (T2VUM) neurons (Figure 14A-D). We identified the muscles innervated by these T2VUM cells as above by using our split-GAL4 lines to express the fluorescent reporter GFP and staining hemithoraces of these flies with phalloidin to visualize the muscles (Figure 14A-D). Two of our T2VUM splits, SS40867 and SS40868, showed innervation of the DLMs and a single DVM (Figure 14A-B), a pattern matching the blowfly mesVUM-MJ (Schlurmann and Hausen, 2003). This putative mesVUM-MJ homolog was the cell with the strongest expression in these two splits. Note that we created both SS40867 and SS40868 using the tdc2-AD split half, i.e. the activation domain corresponding to tyrosine decarboxylase 2 (TDC2), the enzyme that synthesizes tyramine, the precursor of octopamine. This suggests that the cells targeted by both SS40867 and SS40868 are octopaminergic, including the mesVUM-MJ, consistent with previous findings in blowflies (Schlurmann and Hausen, 2003).

**Figure 14:**
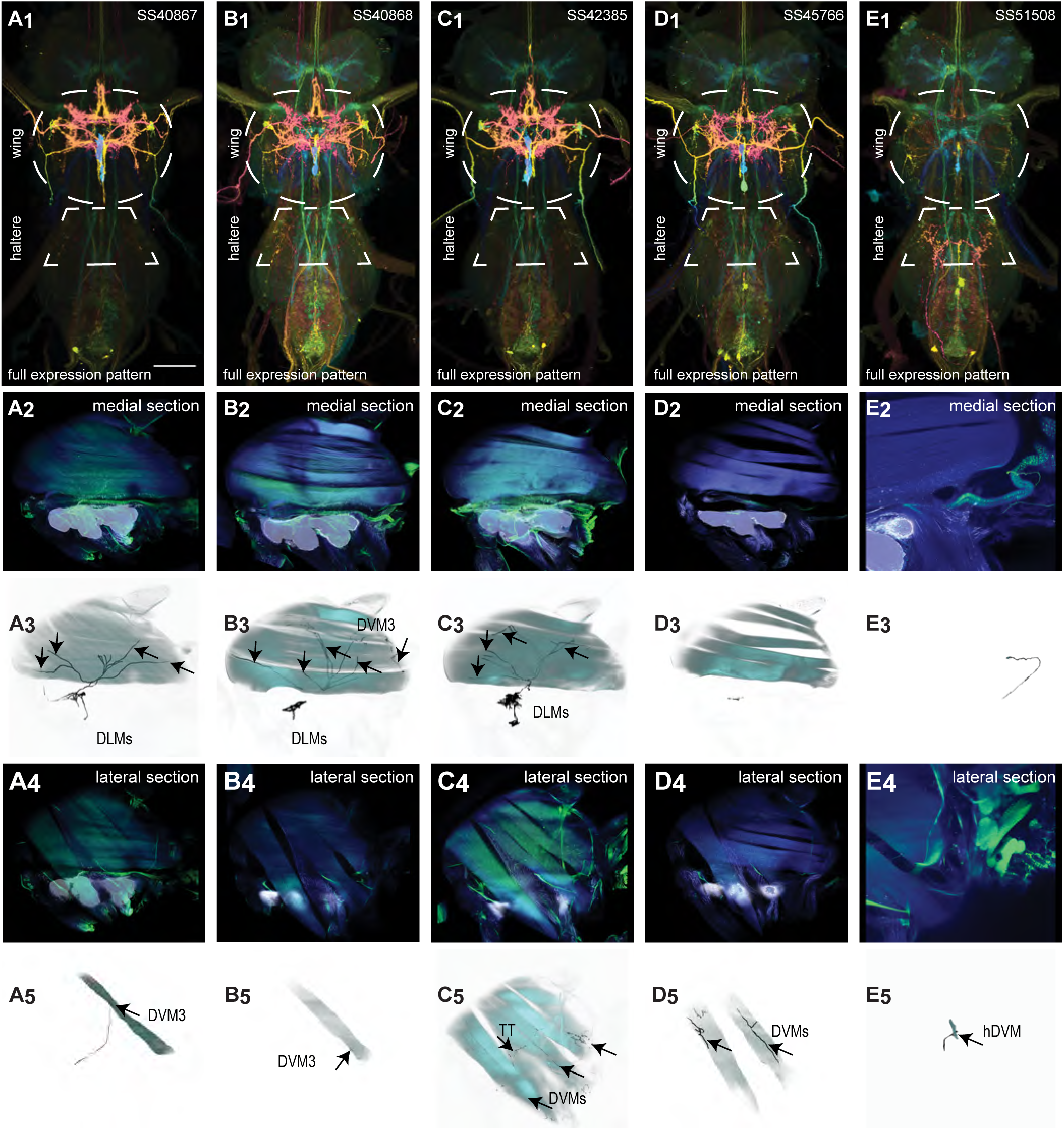
Morphology of VUMs in our split lines. (**A_1_-E_1_**) Color MIPs of the full expression pattern of each split (SS40867, SS40868, SS42385, SS45766, SS51508) in the VNC of males crossed with pJFRC51-3xUAS-Syt::smGFP-HA in su(Hw)attP1; pJFRC225-5xUAS-IVS-myr::smGFP-FLAG in VK00005, aligned to the JRC 2018 VNC Unisex template. Scale bar in A_1_ represents 50 μm. Wing and haltere neuropils are indicated by dashed white outlines. (**A-E_2_**) Medial views of the muscles and their motoneuron innervation in thoraxes stained with phalloidin conjugated with Alexa 633. Phalloidin staining is shown in blue, GFP in green, and nc82 (Bruchpilot, to label the nervous system) in gray. (**A-E_3_**) Segmented medial muscle images. (**A-E_4_**) Lateral views of the muscles and their motoneuron innervation. (**A-E_5_**) Segmented lateral muscle images. Males were used for all images.

A third T2VUM driver line, SS42385, shows innervation of the DLMs, all DVMs, and the tergotrochanter (tt) muscle. This split may have the same mesVUM-MJ homolog that is present in SS40867 and SS40868, as well as homologs to mesVUM-TT (T2VUM targeting the tergotrochanter muscle), mesVUM-PM (T2VUM targeting all power muscles), and a T2VUM which targets specifically DVMs. The final T2VUM line, SS45766, shows innervation of only DVMs and not DLMs (Figure 14D). Another line, SS51508, targeted a single VUM neuron whose soma was not in T2 but in the posterior VNC at the most anterior abdominal segment (Figure 14E) and whose dendrites mainly innervate the haltere neuropil. Phalloidin imaging revealed that the VUM targeted in SS51508 innervates a haltere muscle, which may be hDVM, the haltere’s serial homolog of DVM (Figure 14E). Note, the SS51508 split line is not completely sparse — it also expresses in a pair of cell bodies in the posterior abdominal ganglion, and we observed a dangling axon in our phalloidin preparation, which could be a severed abdominal motoneuron.

We used MCFO to obtain images of individual neurons targeted by our VUM split lines, shown in Figure 15. MCFO often has stronger expression than the GFP reporter line that we used for phalloidin double-labeling experiments, so although the SS40867, SS40868 and SS45766 lines appear sparse in Figure 14A-E, MCFO suggests that each of these splits actually has expression in multiple T2VUM neurons (Figure 15G). We did not create any split lines targeting VUMs in T1 or T3. Our splits do not target all T2VUMs; *Drosophila* has at least three more T2VUMs that did not appear in the MCFO of our splits. Figure 15G lists the T2VUMs that we can identify in the VNC MCFO images of each of our splits (black) and the muscle fiber innervation that was observed in phalloidin experiments with our splits (red). Oddly, SS42385 had fewer T2VUMs identified in MCFO but innervated more muscles than SS40867. This may be due to leaky expression in MCFO that causes labeling of cells that do not normally appear in the full expression pattern, or it could be due to weak GFP labeling resulting in DVM and tt innervation being too dim to see in phalloidin experiments.

**Figure 15:**
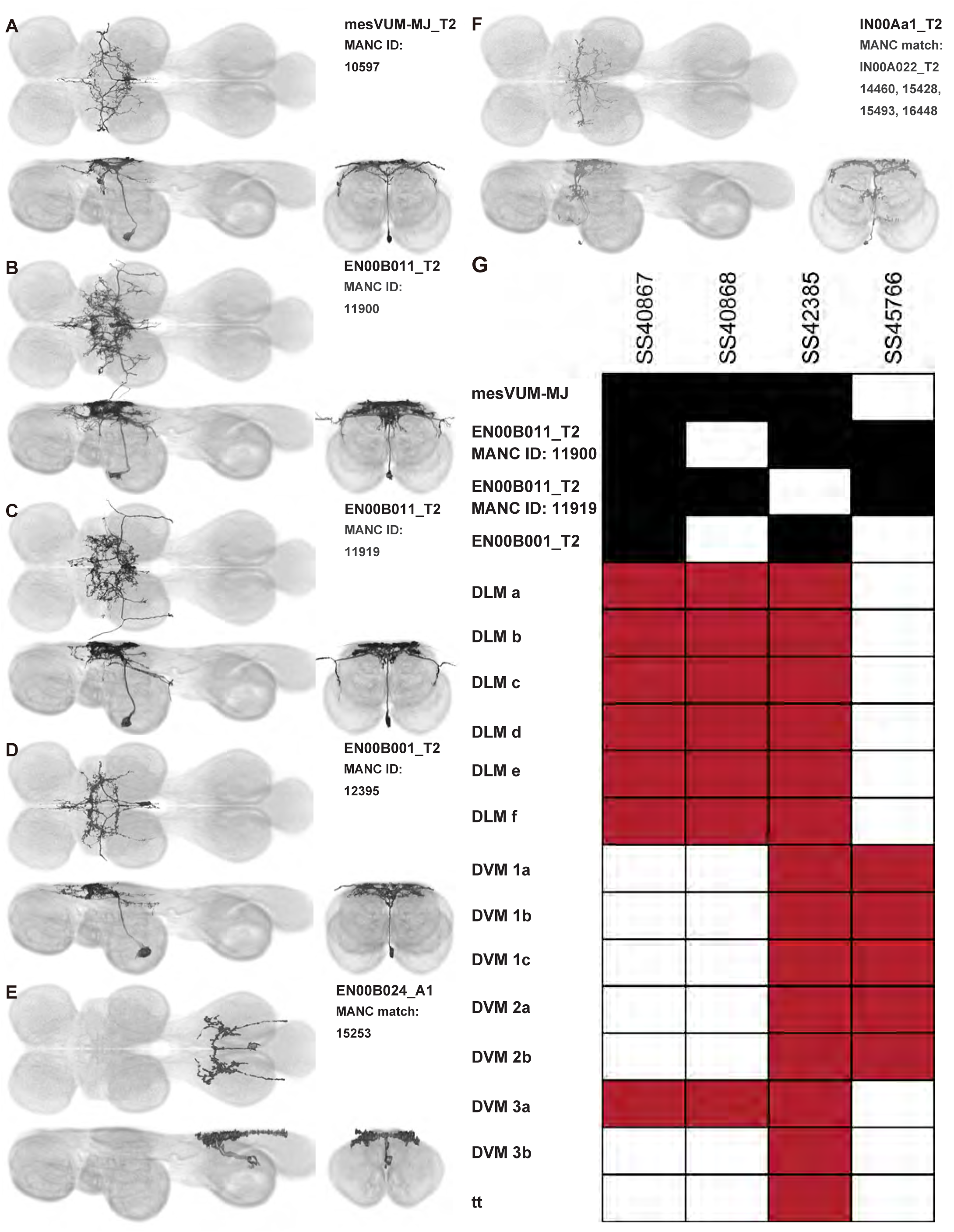
Morphology of individual VUMs and a VPM in our splits, and a table of expression in our T2VUM splits. Segmented multicolor flipout images aligned to the JRC 2018 VNC Unisex template. The driver lines were SS40867, SS46645, SS42385, SS40867, SS51508, SS48268 respectively.

In addition to the VUMs, we also observed a mesothoracic ventral paired median (VPM) neuron in driver line SS48268 (Figure 15F, single cell segmented from this pair). This interneuron, which we call IN00Aa1_T2, has similar wing neuropil innervation to the VUMs.

### Interneuron splits and MCFO images of individual interneurons

We generated 143 driver lines targeting a total of 157 dorsal VNC interneurons, cells that are upstream of efferent motoneurons and VUMs and that may play crucial roles in executing and coordinating wing behaviors. Figure 16 and Figure 16—figure supplements 1-7 show segmented images of intrasegmental interneurons, whose processes are confined to a single neuromere; Figure 17 and Figure 17—figure supplements 1-8 show segmented images of intersegmental interneurons, which connect two or more neuromeres.

**Figure 16:**
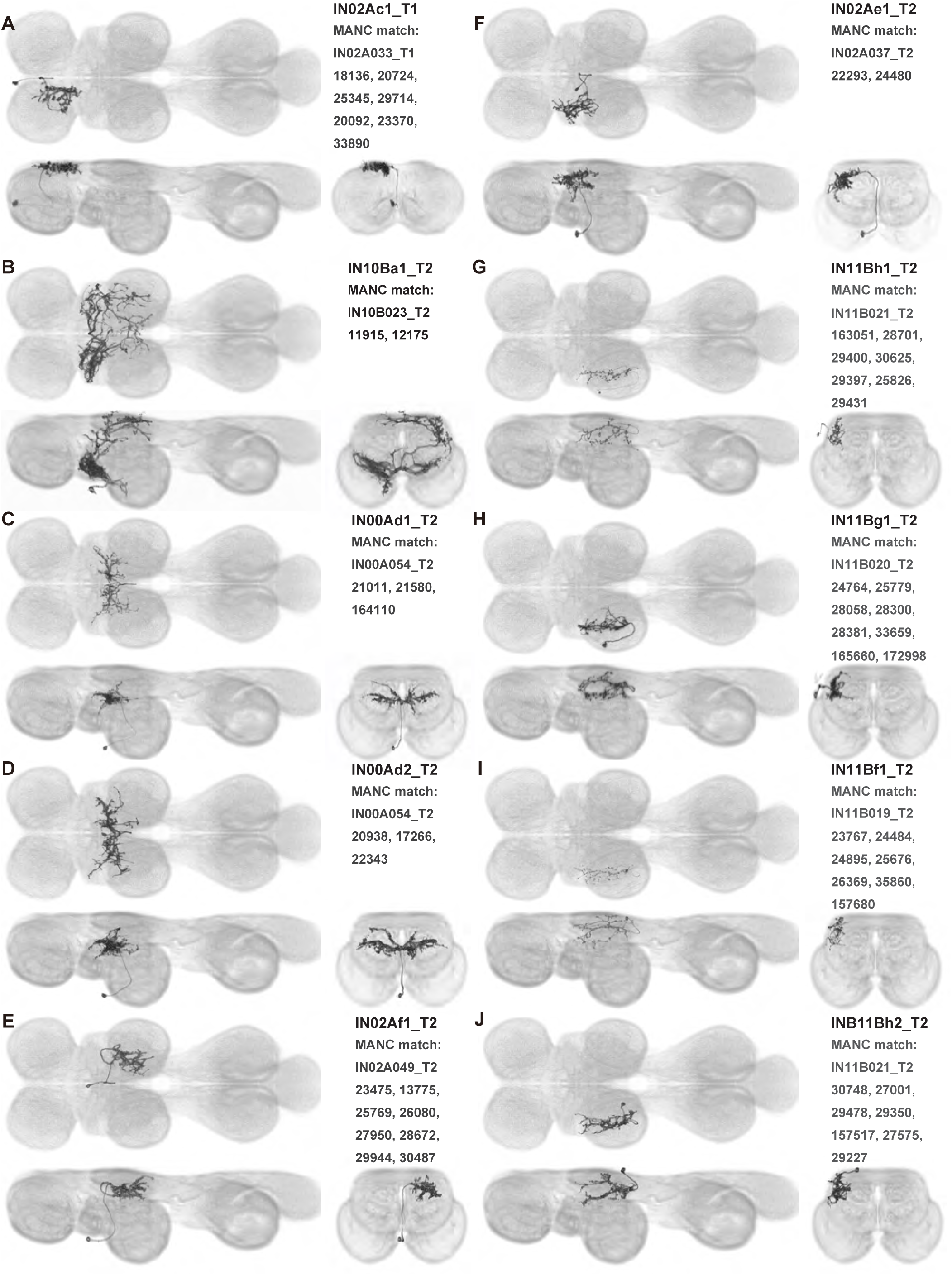
Morphology of identified intrasegmental interneurons. MCFO was used to separate interneurons. Each neuron is shown from three views—horizontal (top left), sagittal (bottom left), and transverse (bottom right)— along with neuron name and hemilineage identity (top right). Images taken from preparations of male flies, segmented to isolate individual neurons, and aligned to the JRC 2018 Unisex VNC template. The driver lines used are: SS44314, SS43546, SS36094, SS36094, SS44314, SS31472, SS49042, SS33409, SS49042, SS25511.

**Figure 17:**
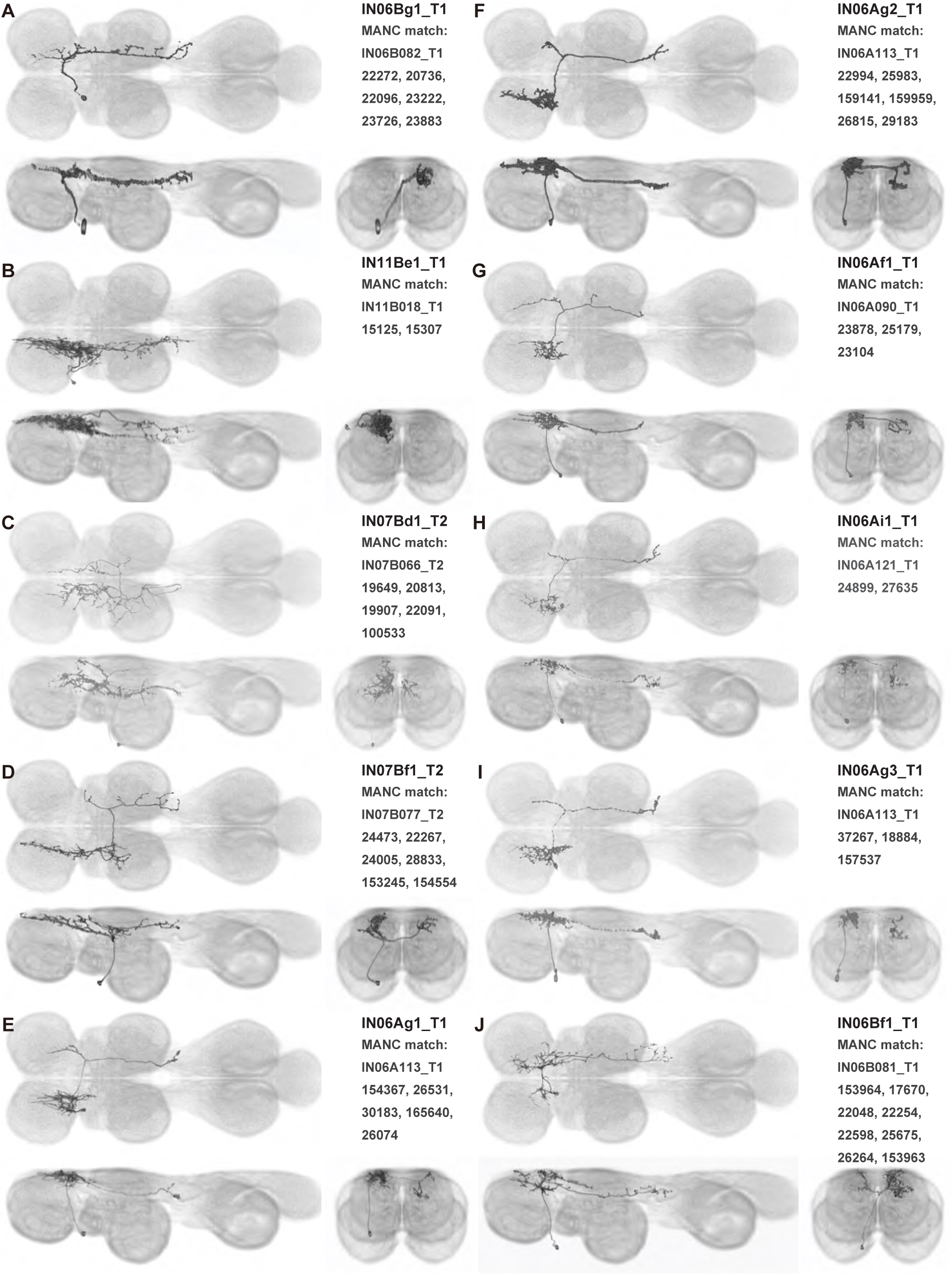
Morphology of identified intersegmental interneurons. MCFO was used to separate interneurons. Each neuron is shown from three views—horizontal (top left), sagittal (bottom left), and transverse (bottom right)— along with neuron name and hemilineage identity (top right). Images taken from preparations of male flies, segmented to isolate individual neurons, and aligned to the JRC 2018 Unisex VNC template. The driver lines used are: SS54495, SS48619, SS60603, SS31899, SS32400, SS54480, SS49807, SS54495, VT014604 (no single cell images were produced from the split GAL4 driver lines targeting this cell, SS31309, SS30330, SS54480, SS54474 or SS54495, so a generation 1 GAL4 image was used), SS25553.

#### Hemilineage identification

To categorize the interneurons in our collection, we identified the developmental origin, i.e. hemilineage, of each of our cells. Neurons belonging to the same hemilineage share similar projection patterns (Lacin and Truman, 2016, Shepherd et al., 2019), neurotransmitter profiles (Lacin et al., 2019), and even functional significance (Harris et al., 2015, Shepherd et al., 2019)—akin to the functional populations arising from common stem cells in the vertebrate spinal cord (Grillner and Jessell, 2009). Each interneuron in our collection was manually annotated for hemilineage type based on the morphology of hemilineage cell types in the adult fly VNC as described in (Shepherd et al., 2019) (Table 1). For example, the IN02Ac1_T1 neuron (Figure 16A) is a member of the 2A hemilineage—i.e. it arises from an A daughter cell of neuroblast 2—in the prothoracic neuromere of the VNC (t1), and thus was assigned the hemilineage classification “2A t1.”

#### Identifying interneuron input and output sites in VNC neuropil regions

To characterize the broad connectivity patterns of the interneurons within our collection, we identified the VNC neuropils (Figure 1C) in which each interneuron had sites of either input or output (Table 1). We used two methods to distinguish presynaptic (input) and postsynaptic (output) terminals in each of our interneurons: manual annotation based on neurite morphology and imaging of terminals labeled with synaptotagmin, a synaptic vesicle-specific protein. For the former, we distinguished between smooth or varicose neurite terminals and labeled these as input or output sites, respectively (Namiki et al., 2018). For the latter, we labeled terminals that co-stained with synaptotagmin as presynaptic (output) and those without co-staining as postsynaptic (input). These two approaches produced consistent results, allowing us to determine the VNC neuropils in which each interneuron had input and output sites (Table 1).

#### Identifying targeted interneurons in the VNC connectome

To further analyze the connectivity of interneurons in our collection, we used segmented, single-cell MCFO images to identify corresponding cells in the male VNC connectome (MANC; (Takemura et al., 2023)). We used a combination of manual annotation and color depth maximum intensity projection (CDM) search (Otsuna et al., 2018, Clements et al., 2024) to obtain matches between light-level images and electron microscopy (EM) volumes, using morphology and hemilineage identity as the primary search criteria. Figure 18 and Figure 18—figure supplements 1-16 compares the morphologies of the skeletonized light level image and EM volume for all segmented interneurons in our collection. Figure 18— figure supplements 17-20 and Figure 18—figure supplement 21 show matches obtained for the MNs and VUMs in our collection, respectively, with the former taken from previous connectome studies (Cheong et al., 2023). The matches for all cell types are also listed in Table 4. We rated the confidence of our MANC matches for each light microscopy morphology type on a scale of 1-3, where 3 indicates the highest confidence (Table 4).

**Figure 18:**
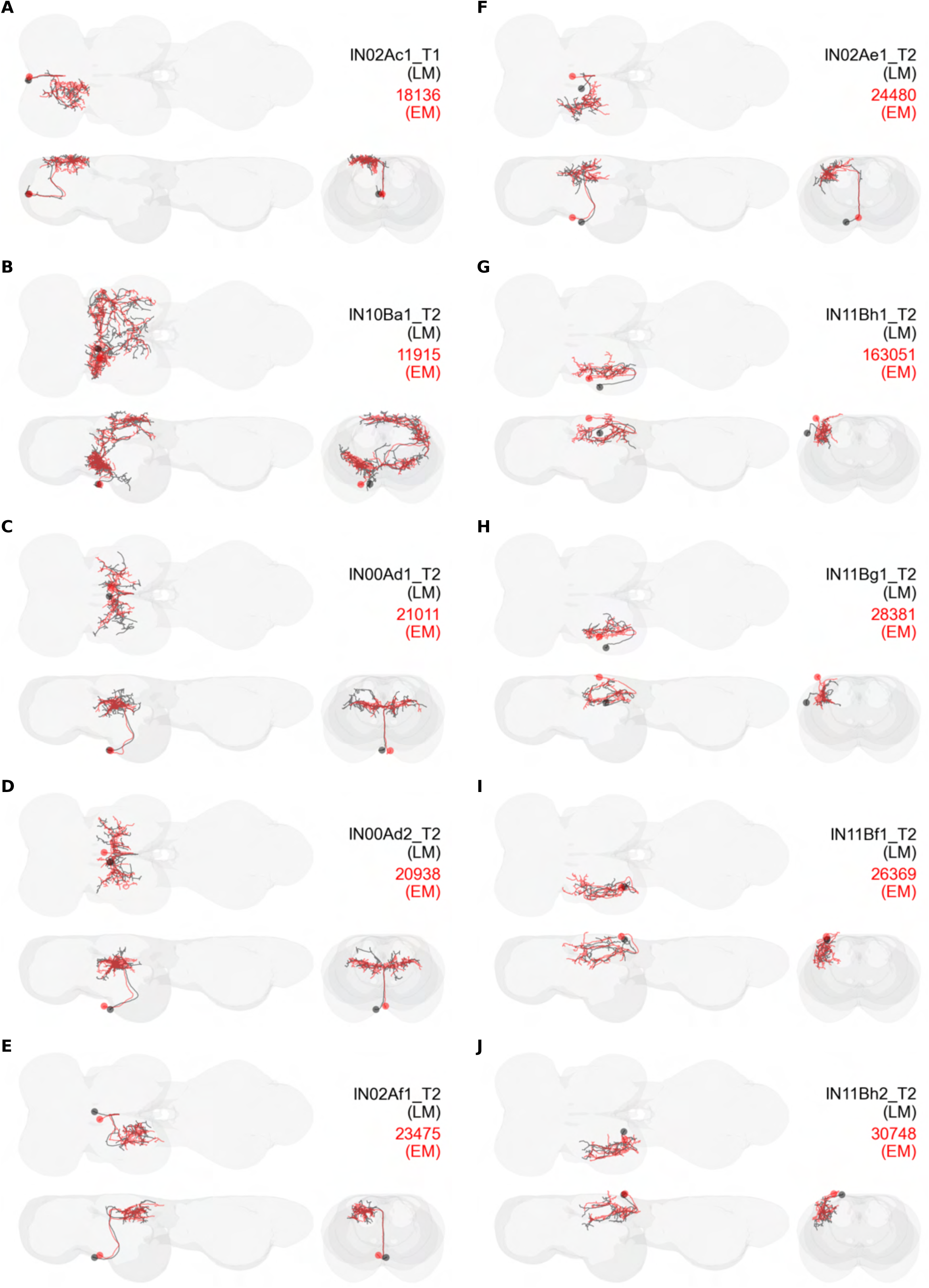
Comparison of interneuron single-cell light microscopy images and MANC electron microscopy matches. Panels (**A**-**I**) each correspond to a single cell type, showing the skeletonized light microscopy (LM) image (black) and the skeleton of the best match obtained from electron microscopy data (red; MANC connectome). The labels in each panel give the neuron name according to our nomenclature (black) and the body ID for the cell in MANC (red).

#### Naming convention

To name the cell types that we defined based on light microscopy images, we followed a modified version of the format for the MANC cell type names. The first two characters indicate whether a cell is an interneuron (IN) or efferent neuron (EN; for VUMs). The next three characters identify the hemilineage, e.g. “02A” for the 2A hemilineage. The next character is a lowercase letter of the alphabet, used to indicate cell type. Cell types were based on the putative MANC matches, so two segmented light microscopy images that both match with cells of a single type in the MANC dataset would be indexed by the same letter. Next, we include an additional integer index to distinguish between cells assigned to the same MANC type, and, finally, an indicator of the cell neuromere, e.g. “_T2” for mesothoracic neuromere. For example, a neuron named IN02Ac1_T1 would be an interneuron from the 2A lineage that is the first cell belonging to the MANC cell type we denoted “c,” and its soma is found in the prothoracic neuromere.

We used a variation of the MANC nomenclature to distinguish between a label from our dataset and one from the MANC. Further, we used both an alphabetic and numeric index for our cell names due to the ambiguity arising from light microscopy images of neurons with subtle differences in morphology. Such images would often map to a single MANC cell type, but could correspond either to different cells, at a level of variation below which the MANC project defines cell types, or different examples of a single cell with developmental variation. Thus, whereas the alphabet letter corresponds to putative cell type from the MANC dataset, the integer index corresponds to morphological variants from our imaging dataset.

In five cases, our light microscopy morphology variants seemed to be particularly uncertain, because two light microscopy variants matched not only the same MANC cell type but the same left-right pair of neurons. These cases are particularly likely to be developmental variation rather than distinct cell types. We marked these variants with asterisks.

### Sexual dimorphism in neuron morphology

Most of the interneurons that we observed were the same in males and females, but some interneurons are sexually dimorphic (Figure 19). We created four splits targeting neurons from hemilineage 17A with somata in T2 (SS40783, SS42438, SS42439 and SS42498). The cells labeled in these splits appear to be subsets of a fruitless-expressing cell type called dMS2 (von Philipsborn et al., 2011). We used MCFO to dissect the separate dMS2 and other 17A t2 cells in the male, resulting in nine segmented dMS2 cell types (dMS2a1_T2-dMS2a9_T2; Figure 16—figure supplements 1, 3, and 4, Figure 17—figure supplement 5). In the line SS40783, six pairs of dMS2 cells are present in males, whereas only four are present in females, a difference that is presumably related to sexual dimorphism in these neurons. Further, the male expression pattern in SS40783 includes an anterior medial arborization which is absent in females. This male-specific medial arborization is present among all four of our dMS2 split lines and may be due to either changes in neuron morphology or differences in cell counts.

**Figure 19:**
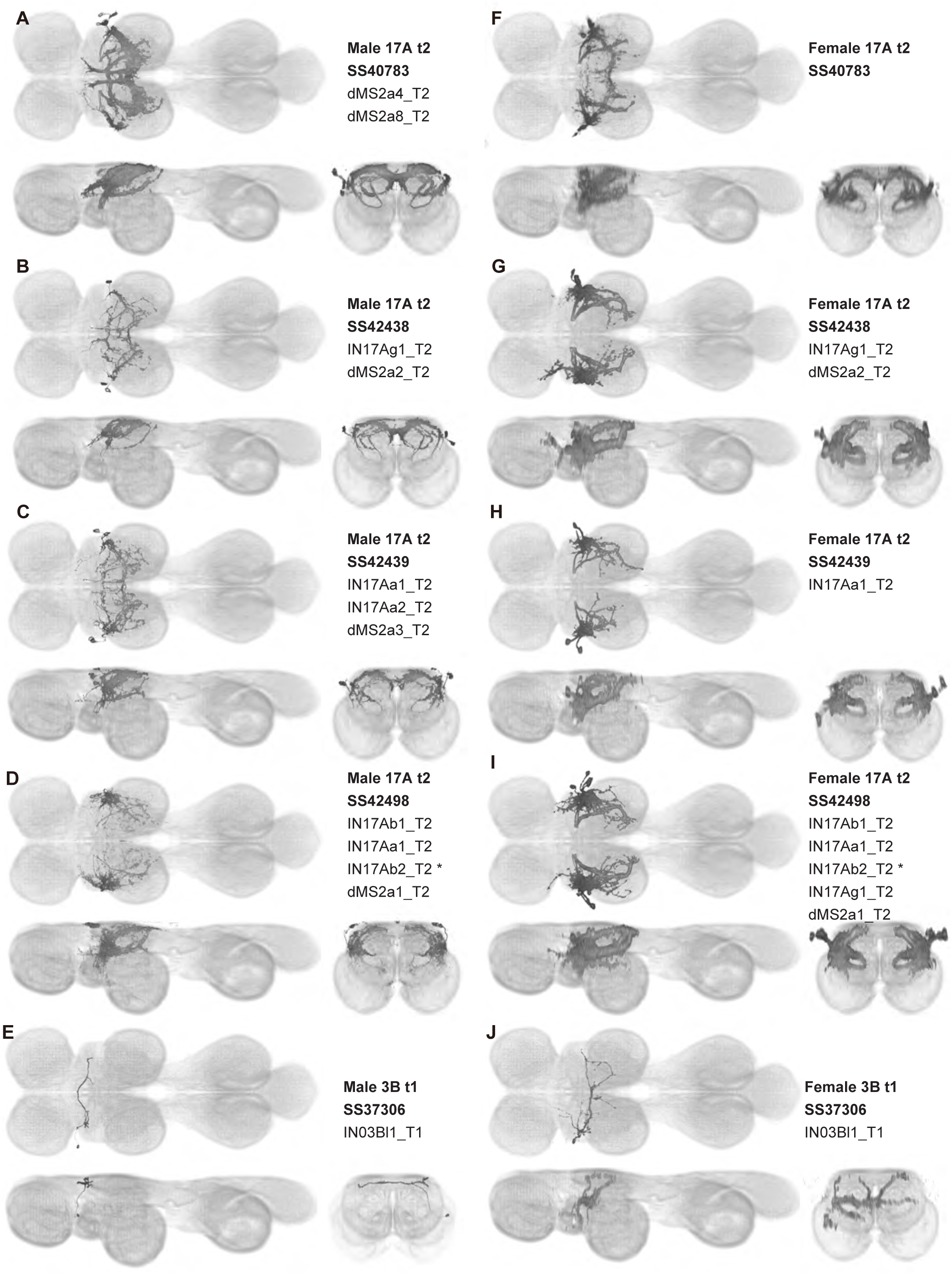
Sexually dimorphic anatomy in split lines targeting two hemilineages. **A** shows the male expression pattern of a split targeting 3B t1, crossed with pJFRC51-3xUAS-Syt::smGFP-HA in su(Hw)attP1; pJFRC225-5xUAS-IVS-myr::smGFP-FLAG in VK00005, not aligned **B-E** show the male expression patterns of 17A t2 in four different split lines, crossed with pJFRC51-3xUAS-Syt::smGFP-HA in su(Hw)attP1; pJFRC225-5xUAS-IVS-myr::smGFP-FLAG in VK00005**. F-J** show the female expression patterns of these same lines, crossed with UAS-CsChrimson. All images are aligned to the JRC 2018 VNC template.

We also observed sexual dimorphism in SS37306, a line targeting IN03Bl1_T1, a single cell in the hemilineage 3B t1 that arborizes in T2. In the female, IN03Bl1_T1has extensive neurites in T2 that innervate both ventral and dorsal neuropils, and it crosses the midline in the intermediate tectulum. In contrast, the male IN03Bl1_T1 has only sparse arborizations in the dorsal-most layer VNC and crosses the midline in the neck neuropil. The neurites of IN03Bl1_T1 are also more posterior in the female than in the male, with female and male neurites more heavily innervating the wing and neck neuropils, respectively.

## Discussion

In this study, we generated a library of cell-type-specific driver lines targeting the dorsal neuropils of the fly VNC and then systematically characterized the function of a subset of these lines with activation and silencing experiments. We produced 195 sparse split-GAL4 driver lines (Table 2) that drive expression in a wide range of dorsal VNC cell types, including wing motoneurons, ventral unpaired median neurons, and local VNC interneurons. Using stochastic labeling, we obtained single-cell images for the expression patterns of these driver lines in order to elucidate their unique morphology, input and output sites, hemilineage origin, and putative connectivity.

This driver line toolkit compliments existing resources (O’Sullivan et al., 2018, Lillvis et al., 2024) and will allow future studies to visualize, manipulate, and study the individual neurons in the VNC, providing a powerful probe for investigating the physiology and function of VNC circuits. In addition, because we matched the neurons in our driver lines to existing connectomic data, our toolkit will allow analyses of these neurons in the broader context of the networks in which they function. We envision the characterization of dorsal VNC neurons provided in this study will serve as a guide for future investigations that seek to elucidate the detailed function of VNC circuitry.

### Quantification of behavior using driver lines

Our collection of sparse driver lines, combined with the sophisticated genetic toolkit for *Drosophila* (Simpson and Looger, 2018, Venken et al., 2011), offers the opportunity to investigate the consequences of targeted neuronal manipulations on the broad range of behaviors controlled through the VNC. To understand the role of individual neurons in wing motor control, we used driver lines that target wing motoneurons and probed the connection between individual wing muscle activity and its role in the performance of both tethered flight (Figure 10) and courtship song (Figure 11), similar to findings reported in previous studies (Lindsay et al., 2017, O’Sullivan et al., 2018).

In tethered flight, we found that optogenetic activation of i2, hg2, DLM, and DVM motoneurons all evoked measurable changes to wing kinematics. These effects are largely consistent with previous studies of flight muscle physiology; however, there are some subtle differences. For instance, while our finding that activation of the steering muscle motoneuron i2 decreases stroke amplitude is consistent with previous imaging studies, we do not observe a very similar effect upon activation of the i1 motoneuron, as the same studies would predict (Lindsay et al., 2017, Melis et al., 2024). Moreover, while increases in the firing rate of both DLM and DVM power muscle motoneurons is linked to increases in wingbeat frequency (Gordon et al., 2006), we only observe a statistically significant effect upon DVM activation. Such inconsistencies may be due to differences in activation versus functional recording experiments, or potentially off-target driver line expression. In combination with these functional recording experiments, our findings provide a basis for further activation screens that may be used to probe upstream flight control circuits in the VNC.

In the case of courtship, we found that silencing the tp2 motoneuron affected all aspects of song, while silencing of the ps1 motoneuron and two hg motoneurons (hg1 and hg2), affected only pulse and sine song, respectively. We did not observe substantial changes to song structure upon silencing of the hg3 MN, a somewhat surprising result given the effects of silencing on the hg1 and hg2 MNs; however, hg3 MN silencing did result in a small but statistically significant decrease in the fraction of slow versus fast pulses. These findings are largely consistent with previous studies of the wing motor system (Shirangi et al., 2013, O’Sullivan et al., 2018), although one such study using a similar methodology reported decreases in pulse song singing upon silencing of the hg1, hg2, and i2 motoneurons, which we do not observe in our data(O’Sullivan et al., 2018). Such discrepancies may be due to the differences in genetic reagents used to carry out experiments. More broadly, our results are consistent with recent connectomic and functional analyses of the VNC courtship song circuit, which shows distinct pathways for pulse and sine song that each preferentially target different wing motoneurons (Lillvis et al., 2024).

### Homologous wing power motoneurons in other insects

Our high-resolution, segmented images of *Drosophila* wing motoneurons facilitate morphological comparisons across insect species. Whereas the morphology of the DLMns of *Drosophila* has previously been investigated using horseradish peroxidase injection (Coggshall, 1978, Sun and Wyman, 1997), the morphology of DVMns in *Drosophila* had never been described. We found that the power MNs, including DVMns, of *Drosophila* are nearly identical to those reported in *Calliphora* (Schlurmann and Hausen, 2007). The most visible difference in DLMn morphology between *Drosophila* and *Calliphora* is that the medial dendrites of DLMna/b extend further both anteriorly and posteriorly in *Calliphora*, while the medial dendrites in *Drosophila* are restricted to a narrower area on the anterior-posterior axis. Thus, the power MNs of *Calliphora* are scaled up but have extremely similar features to *Drosophila* power MNs. Future morphological data from a wider range of fly species will shed light on how broadly this anatomical conservation of power motoneurons extends throughout the Diptera.

Power muscles and motoneurons associated with the front wing have been described in more distantly related insects, particularly in Lepidoptera, making some comparisons across orders possible (Table 3). In moths, as in *Drosophila*, the power muscles comprise two orthogonally arranged muscle sets, the DLMs and DVMs. Despite differences in the number and arrangement of these muscles, the motoneurons innervating power muscles in flies and moths share substantial homology, and the dendritic arbors of these power MNs are located predominantly in the dorsal mesothoracic nerve cord (Agee and Orona, 1988, Ando et al., 2011, Duch et al., 2000, Kondoh and Obara, 1982).

**Table 3:** Previous power muscle and motoneuron terminology used in the literature. The terminology used to name dorsolongitudinal (DLM) and dorsoventral (DVM) power muscles, their muscle fibers or their motoneurons in 105 previous articles from 1970 to present is summarized. The fly DLM comprises five muscle fibers, which have been assigned been named by some studies in order from ventral to dorsal (V to D) and by others from dorsal to ventral (D to V). The terminology of this study is described in the top row. *References in Table 3:* *1. Levine, J.D., & Wyman, R.J. (1973). Neurophysiology of flight in wild-type and a mutant Drosophila. Proc. Nat. Acad. Sci. USA, 70(4), 1050-1054* *2. Ewing, A.W. (1977) The neuromuscular basis of courtship song in Drosophila: The role of the indirect flight muscles. J. comp. Physiol. 119, 249-265.* *3. Fernandes, J.J., Celniker, S.E., & VijayRaghavan, K. (1996). Development of the indirect muscle attachment sites in Drosophila: Role of the PS integrins and the stripe gene. Developmental Biology 176, 166-184.* *4. Ewing, A.W. (1979). The neuromuscular basis of courtship song in Drosophila: The role of the direct and axillary wing muscles. J. Comp. Physiol. 130, 87-93.* *5. Edgecomb, R.S., Ghetti, C., & Schneiderman, A.M. (1993). Bendless alters thoracic musculature in Drosophila. Journal of Neurogenetics 8(4), 209-219. DOI: 10.3109/01677069309083449* *6. Sandstrom, D.J., Bayer, C.A., Fristrom, J.W., & Resifo, L.L. (1997). Broad-Complex transcription factors regulate thoracic muscle attachment in Drosophila. Developmental Biology, 181, 168-185.* *7. Sandstrom, D.J., & Restifo, L.L. (1999). Epidermal tendon cells require Broad Complex function for correct attachment of the indirect flight muscles in Drosophila melanogaster. Journal of Cell Science 112, 4051-4065.* *8. Rivlin, P.K., Gong, A., Schneiderman, A.M., & Booker, R. (2001). The role of Ultrabithorax in the patterning of adult thoracic muscles in Drosophila melanogaster. Dev Genes Evol 211, 55-66. DOI 10.1007/s004270000126* *9. Gordon, S., & Dickinson, M.H. (2006). Role of calcium in the regulation of mechanical power in insect flight. PNAS 103(11), 4311-4315. DOI 10.1073/pnas.0510109103* *10. Tanouye, M.A., & Wyman, R.J. (1980). Motor outputs of the giant fiber in Drosophila. Journal of Neurophysiology, 44(2), 405-421.* *11. Thomas, J.B., & Wyman, R.J. (1983). Normal and mutant connectivity between identified neurons in Drosophila. TINS 214-219.* *12. Euk Oh, C., McMahon, R., Benzer, S., & Tanouye, M.A. (1994). bendless, a Drosophila gene affecting neuronal connectivity, encodes a Ubiquitin-conjugating enzyme homolog. The Journal of Neuroscience, 14(5), 3166-3179.* *13. Kroll, J.R., Wong, K.G., Siddiqui, F.M., & Tanouye, M.A. (2015). Disruption of endocytosis with the dynamin mutant shibirets1 suppresses seizures in Drosophila. Genetics, 201(3), 1087-1102. doi: 10.1093/genetics/201.3.NP* *14. Augustin, H., McGourty, K., Allen, M.J., Adcott, J., Wong, C.T., Boucrot, E., & Partridge, L. (2018). Impact of insulin signaling and proteasomal activity on physiological output of a neuronal circuit in aging Drosophila melanogaster. Neurobiology of Aging, 66, 149-157. doi: 10.1016/j.neurobiolaging.2018.02.027* *15. Baird, D.H., Schalet, A.P., & Wyman, R.J. (1990). The Passover locus in Drosophila melanogaster: Complex complementation and different effects on the giant fiber neural pathway. Genetics, 126, 1045-1059.* *16. Allen, M.J., Shan, X., Caruccio, P., Froggett, S.J., Moffat, K.G., & Murphey, R.K. (1999). Targeted expression of truncated Glued disrupts giant fiber synapse formation in Drosophila. The Journal of Neuroscience, 19(21), 9374-9384.* *17. Allen, M.J., Shan, X., & Murphey, R.K. (2000). A role for Drosophila Drac1 in neurite outgrowth and synaptogenesis in the giant fiber system. Molecular and Cellular Neuroscience, 16, 754-765. doi: 10.1006/mcne.2000.0903* *18. Godenschwege, T.A., Hu, H., Shan-Crofts, X., Goodman, C.S., & Murphey, R.K. (2002). Bi-directional signaling by Semaphorin 1a during central synapse formation in Drosophila. Nature Neuroscience, 5, 1294-1301.* *19. Murphey, R.K., Froggett, S.J., Caruccio, P., Shan-Crofts, X., Kitamoto, T., & Godenschwege, T.A. (2003). Targeted expression of shibire^ts^ and semaphorin 1a reveals critical periods for synapse formation in the giant fiber of Drosophila. Development, 130(16), 3671-3682. doi: 10.1242/dev.005682* *20. Allen, M.J., & Murphey, R.K. (2007). The chemical component of the mixed GF-TTMn synapse in Drosophila melanogaster uses acetylcholine as its neurotransmitter. European Journal of Neuroscience, 26, 439-445.* *21. Uthaman, S.B., Godenschwege, T.A., & Murphey, R.K. (2008). A mechanism distinct from Highwire for the Drosophila ubiquitin conjugase Bendless in synaptic growth and maturation. The Journal of Neuroscience, 28(34), 8615-8623.* *22. Godenschwege, T.A., & Murphey, R.K. (2009). Genetic interaction of Neuroglian and Semaphorin1a during guidance and synapse formation. Journal of Neurogenetics, 23(1-2), 147-155. DOI: 10.1080/01677060802441380* *23. Zhao, X.-L., Wang, W.-A., Tan, J.-X., Huang, J.-K., Zhang, X., Zhang, B.-Z., Wang, Y.-H., YangCheng, H.-Y., Zhu, H.-L., Sun, X.-J., & Huang, F.-D. (2010). Expression of ß-Amyloid induced age-dependent presynaptic and axonal changes in Drosophila. The Journal of Neuroscience, 30(4):1512-1522.* *24. Meija, M., Heghinian, M.D., Busch, A., Armishaw, C.J., Mari, F., & Godenschwege, T.A. (2010). A novel approach for in vivo screening of toxins using the Drosophila Giant Fiber circuit. Toxicon, 56(8), 1398-1407. doi: 10.1016/toxicon.2010.08.005* *25. Mejia, M., Heghinian, M.D., Busch, A., Mari, F., & Godenschwege, T.A. (2012). Paired nanoinjection and electrophysiology assay to screen for bioactivity of compounds using the Drosophila melanogaster Giant Fiber System. Journal of Visualized Experiments, (62), 3597. doi: 10.3791/3597* *26. Lin, J.-Y., Wang, W.-A., Zhang, X., Liu, H.-Y., Zhao, X.-L., & Huang, F.-D. (2014). Intraneuronal accumulation of Aß42 induces age-dependent slowing of neuronal transmission in Drosophila. Neurosci Bull 30(2), 185-190. DOI: 10.1007/s12264-013-1409-9* *27. Pezier, A.P., Jezzini, S.H., Bacon, J.P., & Blagburn, J.M. (2016). Shaking B mediates synaptic coupling between auditory sensory neurons and the giant fiber of Drosophila melanogaster. PLoS ONE, 11(4), e0152211. doi: 10.1371/journal.pone.0152211* *28. Borgen, M., Rowland, K., Boerner, J., Lloyd, B., Khan, A., & Murphey, R. (2017). Axon termination, pruning, and synaptogenesis in the giant fiber system of Drosophila melanogaster is promoted by Highwire. Genetics, 205(3), 1229-1245. doi: 10.1534/genetics.116.197343* *29. Lee, J., Iyengar, A., & Wu, C.-F. (2019). Distinctions among electroconvulsion- and proconvulsant-induced seizure discharges and native motor patterns during flight and grooming: quantitative spike pattern analysis in Drosophila flight muscles. Journal of Neurogenetics, 33(2). doi: 10.1080/01677063.2019.1581188* *30. Huang, J.-K., Ma, P.-L., Ji, S.-Y., Zhao, X.-L., Tan, J.-X., Sun, X.-J., & Huang, F.-D. (2013). Age-dependent alterations in the presynaptic active zone in a Drosophila model of Alzheimer’s disease. Neurobiology of Disease, 51, 161-167. doi: 10.1016/j.nbd.2021.11.06* *31. Hummon, M.R., & Costello, W.J. (1988). Induced neuroma formation and target muscle perturbation in the giant fiber pathway of the Drosophila temperature-sensitive mutant shibire. Roux’s Arch Dev Biol, 197, 383-393.* *32. De la Pompa, J.L., Garcia, J.R., & Ferrus, A. (1989). Genetic analysis of muscle development in Drosophila melanogaster. Developmental Biology 131, 439-454.* *33. Hummon, M.R., & Costello, W.J. (1993). Flight muscle formation in Drosophila mosaics: Requirement for normal shibire function of endocytosis. Roux’s Arch Dev Biol 202, 95-102.* *34. Coggshall, J.C. (1978). Neurons associated with the dorsal longitudinal flight muscles of Drosophila melanogaster. J. Comp. Neur. 177, 707-720.* *35. Sun, Y.-A., & Wyman, R.J. (1997). Neurons of the Drosophila Giant Fiber System: I. Dorsal Longitudinal Motor Neurons. The Journal of Comparative Neurology, 387, 157-166.* *36. Engel, J.E., & Wu, C.-F. (1992). Interactions of membrane excitability mutations affecting potassium and sodium currents in the flight and giant fiber escape systems of Drosophila. J Comp Physiol A, 171, 93-104.* *37. Lee, J., & Wu, C.-F. (2002). Electroconvulsive seizure behavior in Drosophila: Analysis of the physiological repertoire underlying a stereotyped action pattern in bang-sensitive mutants. The Journal of Neuroscience, 22(24), 11065-11079.* *38. Tanouye, M.A., & Wyman, R.J. (1981). Inhibition between flight motoneuron in Drosophila. J Comp Physiol, 144, 345-355.* *39. Hummon, M.R., & Costello, W.J. (1992). Cell lineage of flight muscle fibers in Drosophila: A fate map of induced shibire phenotype in mosaics. Roux’s Arch Dev Biol 201, 88-94.* *40. Costello, W.J., & Wyman, R.J. (1986). Development of an indirect flight muscle in a muscle-specific mutant of Drosophila melanogaster. Developmental Biology, 118(1), 247-258. DOI: 10.1016/0012-1606(86)90092-8* *41. Atreya, K.B., & Fernandes, J.J. (2008). Founder cells regulate fiber number but not fiber formation during adult myogenesis in Drosophila. Developmental Biology, 321(1), 123-140. doi: 10.1016/j.ydbio.2008.06.023* *42. Farrell, E.R., Fernandes, J., & Keshishian, H. (1996). Muscle organizers in Drosophila: The role of persistent larval fibers in adult flight muscle development. Developmental Biology 176, 220-229.* *43. Hebbar, S., & Fernandes, J.J. (2004). Pruning of motor neuron branches establishes the DLM innervation pattern in Drosophila. Journal of Neurobiology, 60(4), 499-516.* *44. Banerjee, S., Lee, J., Wu, C.-F., & Hasan, G. (2004). Loss of flight and associated neuronal rhythmicity in inositol 1,4,5-triphosphate receptor mutants of Drosophila. The Journal of Neuroscience, 24(36), 7869-7878. DOI: 10.1523/JNEUROSCI.0656-04.2004* *45. Chaturvedi, D., Prabhakar, S., Aggarwal, A., Atreya, K.B., & VijayRaghavan, K. (2019). Adult Drosophila muscle morphometry through microCT reveals dynamics during ageing. Open Biol. 9: 190087 http://dx.doi.org/10.1098/rsob.190087* *46. Fernandes, J., Bate, M., & VijayRaghavan, K. (1991). Development of the indirect flight muscles of Drosophila. Development 113, 67-77.* *47. Lee, J.C., VijayRaghavan, K., Celniker, S.E., & Tanouye, M.A. (1995). Identification of a Drosophila muscle development gene with structural homology to mammalian early growth response transcription factors. Proc. Natl. Acad. Sci. USA 92, 10344-10348.* *48. Fernandes, J.J., & Keshishian, H. (1996). Patterning the dorsal longitudinal flight muscles (DLM) of Drosophila: Insights from the ablation of larval scaffolds. Development 122, 3755-3763.* *49. Fernandes, J.J., & Keshishian, H. (1998). Nerve-muscle interactions during flight muscle development in Drosophila. Development 125, 1769-1779.* *50. Schmid, A., Chiba, A., & Doe, C.Q. (1999). Clonal analysis of Drosophila embryonic neuroblasts: neural cell types, axon projections and muscle targets. Development 126, 4653-4689* *51. Hebbar, S., & Fernandes, J.J. (2005). A role for Fas II in the stabilization of motor neuron branches during pruning in Drosophila. Developmental Biolog, 285(1), 185-199. doi: 10.1016/j.ydbio.2005.06.015* *52. Hebbar, S., & Fernandes, J.J. (2010). Glial remodeling during metamorphosis influences the stabilization of motor neuron branches in Drosophila. Developmental Biology, 340(2), 344-354. doi: 10.1016/j.ydbio.2010.01.005* *53. Lee, J., & Wu, C.-F. (2006). Genetic modifications of seizure susceptibility and expression by altered excitability in Drosophila Na+ and K+ channel mutants. J Neurophysiol 96, 2465-2478. doi:10.1152/jn.00499.2006.* *54. Zhang, T., Wang, Z., Wang, L., Luo, N., Jiang, L., Liu, Z., Wu, C.-F., & Dong, K. (2013). Role of the DSC1 channel in regulating neuronal excitability in Drosophila melanogaster: Extending nervous system stability under stress. PLOS Genetics. doi: 10.1371/journal.pgen.1003327* *55. Engel, J.E., & Wu, C.-F. (1996). Altered habituation of an identified escape circuit in Drosophila memory mutants. The Journal of Neuroscience, 16(10), 3486-3499.* *56. Lehmann, F.-O., Skandalis, D.A., & Berthe. (2013). Calcium signalling indicates bilateral power balancing in the Drosophila flight muscle during manoeuvring flight. J R Soc Interface 10, 20121050. doi:10.1098/rsif.2012.1050* *57. Restifo, L.L., & White, K. (1992) Mutations in a steroid hormone-regulated gene disrupt the metamorphosis of internal tissues in Drosophila: Salivary glands, muscle, and gut. Roux’s Arch Dev Biol., 201, 221-234.* *58. Gorczyca, M., & Hall, J.C. (1984). Identification of a cholinergic synapse in the giant fiber pathway of Drosophila using conditional mutations of acetylcholine synthesis. Journal of Neurogenetics, 1(4), 289-313.* *59. Harcombe, E.S., & Wyman, R.J. (1977). Output pattern generation by Drosophila flight motoneurons. Journal of Neurophysiology, 40(5), 1066-1077.* *60. Harcombe, E.S., & Wyman, R.J. (1978). The cyclically repetitive firing sequences of identified Drosophila flight motoneurons. J. comp. Physiol., 123, 271-279.* *61. Elkins, T., Ganetzky, B., & Wu, C.-F. (1986). A Drosophila mutation that eliminates a calcium-dependent potassium current. Proc. Natl. Acad. Sci. USA, 83, 8415-8419.* *62. De Rose, F., Marotta, R., Talani, G., Catelani, T., Solari, P., Poddighe, S., Borghero, G., Marrosu, F., Sanna, E., Kasture, S., Acquas, E., & Liscia, A. (2017). Differential effects of phytotherapic preparations in the hSOD1 Drosophila melanogaster model of ALS. Scientific Reports, 7, 41059.* *63. Allen, M.A., & Godenschwege, T.A. (2010). Electrophysiological recordings from the Drosophila giant fiber system (GFS). Cold Spring Harb Protoc, 7. doi:10.1101/pdb.prot5453* *64. Kadas, D., Tzortzopoulos, A., Skoulakis, E.M.C., & Consoulas, C. (2012). Constitutive activation of Ca2+/Calmodulin-Dependent Protein Kinase II during development impairs central cholinergic transmission in a circuit underlying escape behavior in Drosophila. The Journal of Neuroscience, 32(1), 170-182. doi: 10.1523/JNEUROSCI.6583-10.2012* *65. Koenig, J.H., & Ikeda, K. (1983). Reciprocal excitation between identified flight motor neurons in Drosophila and its effect on pattern generation. J Comp Physiol, 150, 305-317.* *66. Hutchinson, K.M., Vonhoff, F., & Duch, C. (2014). Dscam1 is required for normal dendrite growth and branching but not for dendritic spacing in Drosophila motoneurons. The Journal of Neuroscience 34(5), 1924-1931.* *67. Koenig, J.H., & Ikeda, K. (1980). Neural interactions controlling timing of flight muscle activity in Drosophila. J. exp. Biol. 87, 121-136.* *68. Koenig, J.H., & Ikeda, K. (1983). Characterization of the intracellularly recorded response of identified flight motor neurons in Drosophila. J Comp Physiol, 150, 295-303.* *69. Levine, J., & Tracey, D. (1973). Structure and function of the giant motorneuron of Drosophila melanogaster. J. comp. Physiol. 87, 213-235.* *70. Levine, J.D., & Hughes, M. (1973). Stereotaxic map of the muscle fibers in the indirect flight muscles of Drosophila melanogaster. J. Morph., 140, 153-158.* *71. Benshalom, G., & Dagan, D. (1981). Electrophysiological analysis of the temperature-sensitive paralytic Drosophila mutant, para ts. J Comp Physiol, 144, 409-417.* *72. Benshalom, G., & Dagan, D. (1985). Drosophila neural pathways: Genetic and electrophysiological analysis. J Comp Physiol, 156, 13-23.* *73. Ikeda, K., Koenig, J.H., & Tsuruhara, T. (1980). Organization of identified flight muscle of Drosophila melanogaster. Journal of Neurocytology, 9, 799-823* *74. Wang, D., Keng, Z.C., Hsu, K., & Tan, C.C. (1989) Drosophila mutants with progressive atrophy in dorsal longitudinal muscles. Journal of Neurogenetics, 6(1), 27-39. DOI: 10.3109/01677068909107098* *75. Koenig, J.H., Goto, J.J., & Ikeda, K. (2015). Novel NMDA receptor-specific desensitization/inactivation produced by ingestion of the neurotoxins, ß-N-methylamine-L-alanine (BMAA) or ß-N-oxalylamino-L-alanine (BOAA/ß-ODAP). Comparative Biochemistry and Physiology Part C: Toxicology & Pharmacology, 167, 43-50. doi: 10.1016/j.cbpc.2014.08.006* *76. Rai, M., Katti, P., & Nongthomba, U. (2016). Spatio-temporal coordination of cell cycle exit, fusion and differentiation of adult muscle precursors by Drosophila Erect wing (Ewg). Mechanisms of Development, 141, 109-118. doi: 10.1016/j.mod.2016.03.004* *77. Trimarchi, J.R., & Schneiderman, A.M. (1995). Flight initiations in Drosophila melanogaster are mediated by several distinct motor patterns. J Comp Physiol A 176, 355-364.* *78. Glasscock, E., & Tanouye, M.A. (2005). Drosophila couch potato mutants exhibit complex neurological abnormalities including epilepsy phenotypes. Genetics, 169(4), 2137-2149. doi: 10.1534/genetics.104.028357* *79. Pavlidis, P., & Tanouye, M.A. (1995). Seizures and failures in the giant fiber pathway of Drosophila bang-sensitive paralytic mutants. The Journal of Neuroscience, 15(8), 5810-5819.* *80. Ikeda, K., & Koenig, J.H. (1988). Morphological identification of the motor neurons innervating the dorsal longitudinal flight muscle of Drosophila melanogaster. The Journal of Comparative Neurology, 273, 436-444.* *81. Consoulas, C., Restifo, L.L., & Levine, R.B. (2002). Dendritic remodeling and growth of motoneurons during metamorphosis of Drosophila melanogaster. The Journal of Neuroscience 22(12), 4906-4917.* *82. Ryglewski, S., Kadas, D., Hutchinson, K., Schuetzler, N., Vonhoff, F., & Duch, C. (2014). Dendrites are dispensible for basic motoneuron function but essential for fine tuning of behavior. PNAS 111(50), 18049-18054. doi/10.1073/pnas.1416247111* *83. Kadas, D., Duch, C., & Consoulas, C. (2019). Postnatal increase in axonal conduction velocity of an identified Drosophila interneuron require fast sodium, L-type calcium and Shaker potassium channels. eNeuro, 6(4), doi: 10.1523/ENEURO.0181-19.2019* *84. Rai, M., & Nongthomba, U. (2013). Effect of myonuclear number and mitochondrial fusion on Drosophila indirect flight muscle organization and size. Experimental Cell Research, 319(17), 2566-2577. doi: 10.1016/j.yexcr.2013.06.021* *85. Koenig, J.H., & Ikeda, K. (2005). Relationship of the reserve vesicle population to synaptic depression in the tergotrochanteral and dorsal longitudinal muscles of Drosophila. Journal of Neurophysiology, 94(3), 2111-2119. doi: 10.1152/jn.00323.2005* *86. Fernandes, J., & VijayRaghavan, K. (1993). The development of indirect flight muscle innervation in Drosophila melanogaster. Development 118, 215-227.* *87. Ryglewski, S., & Duch, C. (2009). Shaker and Shal mediate transient calcium-independent potassium current in a Drosophila flight motoneuron. Journal of Neurophysiology, 102(6), 3673-3688. doi: 10.1152/jn.00693.2009* *88. Nachtigall, W., & Wilson, D.M. (1967). Neuro-muscular control of Dipteran flight. J. Exp. Biol. 47, 77-97.* *89. Ikeda, K. (1977) Flight motor innervation of a flesh fly. In: Hoyle G. editor. Identified Neurons and Behavior of Arthropods. Springer. pp. 357-358* *90. Miyan, J.A., & Ewing, A.W. (1985). How Diptera move their wings: A re-examination of the wing base articulation and muscle systems concerned with flight. Philosophical Transactions of the Royal Society of London. Series B, Biological Sciences, 311(1150), 271-302.* *91. Heide, G. (1979). Proprioceptive feedback dominates the central oscillator in the patterning of the flight motoneuron output in Tipula (Diptera). J. Comp. Physiol., 134, 177-189.* *92. Schlurmann, M., & Hausen, K. (2007). Motoneurons of the flight power muscles of the blowfly Calliphora erythrocephala: Structures and mutual dye coupling. The Journal of Comparative Neurology, 500, 448-464.* *93. Adams, M.E., & Miller, T.A. (1980). Neural and behavioral correlates of pyrethroid and DDT-type poisoning in the house fly, Musca domestica L. Pesticide Biochemistry and Physiology, 13(2), 137-147. doi: 10.1016/0048-3575(80)90065-6* *94. Bayline, R.J., Duch, C. & Levine, R.B. (2001). Nerve-muscle interactions regulation motor terminal growth and myoblast distribution during muscle development. Developmental Biology 231, 348-363. doi:10.1006/dbio.2001.0158* *95. Duch, C., Bayline, R.J., & Levine, R.B. (2000). Postembryonic development of the dorsal longitudinal flight muscle and its innervation in Manduca sexta. The Journal of Comparative Neurology, 422, 1-17* *96. Duch, C., & Levine, R.B. (2000). Remodeling of membrane properties and dendritic architecture accompanies the postembryonic conversion of a slow into a fast motoneuron. The Journal of Neuroscience, 20(18), 6950-6961.* *97. Duch, C., & Mentel, T. (2003). Stage-specific activity patterns affect motoneuron axonal retraction and outgrowth during the metamorphosis of Manduca sexta. European Journal of Neuroscience, 17, 945-962.* *98. Tu, M.S., & Daniel, T.L. (2004). Submaximal power output from the dorsolongitudinal flight muscles of the hawkmoth Manduca sexta. J Exp Biol, 207(26), 4651-4662. doi: 10.1242/jeb.01321* *99. George, N.T., & Daniel, T.L. (2011). Temperature gradients in the flight muscles of Manduca sexta imply a spatial gradient in muscle force and energy output. J Exp Biol, 214(6), 894-900. doi: 10.1242/jeb.047969* *100. George, N.T., Sponberg, S., & Daniel, T.L. (2012). Temperature gradients drive mechanical energy gradients in the flight muscle of Manduca sexta. J Exp Biol, 215(3), 571-579. doi: 10.1242/jeb.062901* *101. Komai, Y. (1998). Augmented respiration in a flying insect. The Journal of Experimental Biology 201, 2359-2366.* *102. Ando, N., & Kanzaki, R. (2004). Changing motor patterns of the 3rd axillary muscle activities associated with longitudinal control in freely flying hawkmoths. Zoological Science, 21, 123-130.* *103. Ando, N., Wang, H., Shirai, K., & Kanzaki, R. (2011). Central projections of the wing afferents in the hawkmoth, Agrius convolvuli. Journal of Insect Physiology, 51(11), 1518-1536. doi: 10.1016/j.jinsphys.2011.08.002* *104. Kondoh, Y., & Obara, Y. (1982). Anatomy of motoneurones innervating mesothoracic indirect flight muscles in the silkmoth Bombyx mori. J. exp. Biol., 98, 23-37* *105. Hanegan, J.L., & Heath, J.E. (1970). Temperature dependence of the neural control of the moth flight system. J. Exp. Biol., 53, 629-639.*

**Table 4:** Matches in the MANC electron microscopy connectome. For each cell type that we defined based on morphology in light microscopy images (LM name), we list the body ids of the best MANC matches and the MANC cell type. The five most uncertain cell types, which are particularly likely to represent developmental variants in morphology rather than distinct cell types, are marked with asterisks. We assigned a confidence rating to each match: 3 = high confidence; 2 = medium confidence; 1 = low confidence. The driver line that is the source of the segmented single-cell image used to represent the cell type in this paper is listed. Other plausible MANC cell type matches are listed.

Similar to *Drosophila*, the DLMns in moths consists of a group of neurons innervating ventral DLM fibers whose somata form a tightly apposed cluster and project ipsilaterally (DLMnc-f in *Drosophila*; Figure 2C-F), along with a motoneuron that innervates two dorsal DLM fibers contralateral to its soma (DLMna/b in *Drosophila*; Figure 2B) (Agee and Orona, 1988, Ando et al., 2011, Duch et al., 2000, Kondoh and Obara, 1982). However, this contralaterally projecting DLMn—DLMna/b in *Drosophila* and DLMn1e in moths—arborizes much more symmetrically in the *Drosophila* VNC than in the moth (Ando et al., 2011, Duch and Levine, 2000, Kondoh and Obara, 1982).

The other set of power motoneurons, the DVMns, exhibit much less similarity between moths and flies, and have only been described in the anterior thorax of silk moths to date (Kondoh and Obara, 1982). In both silk moths and fruit flies, the DVMns can be grouped into three sets of MNs that exit the VNC in distinct nerves. Silk moth DVMns tend to be more lateralized than fruit fly DVMns, and, in contrast to the DLMns, silk moths have a population of DVMns for which there is no clear homologue among the *Drosophila* power motoneurons (Kondoh and Obara, 1982). This seemingly silk-moth-unique MN, however, innervates a DVM that is sometimes called the tergotrochanter (Kondoh and Obara, 1982), and so might be more directly comparable to the *Drosophila* tt MN. The factors that presumably allow this greater evolvability in the DVMns versus the DLMns is a fascinating topic for future research.

### Homologous wing control motoneurons in other insects

The morphology of many *Drosophila* wing control MNs has been reported in previous studies (O’Sullivan et al., 2018). We are the first to describe the anatomy of the hg3 MN in the VNC and to produce sparse driver lines targeting this MN. Moreover, while the morphology of i1 MN has previously been described (Trimarchi and Schneiderman, 1994), but we created the first sparse driver line targeting this MN.

The morphology of steering MNs may vary dramatically between different Dipteran genera. For example, in *Drosophila*, b1 MN has a relatively simple morphology, with two long major fibers: one that projects across the midline to the contralateral side and one that projects posteriorly (Figure 6A, C). In contrast, b1 MN in *Calliphora* has a completely different morphology: the posterior main branch found in *Drosophila* appears to be absent in *Calliphora*, and instead, the *Calliphora* b1 MN has numerous relatively short branches projecting both anteriorly and posteriorly (Fayyazuddin and Dickinson, 1996). Thus, the b1 MN of *Calliphora* more closely resembles i2 MN in *Drosophila* than b1 MN in *Drosophila.* While such differences in b1 MN morphology may relate to differences in its presynaptic partners, a putatively important haltere sensory input to b1 MN has been shown to be similar in *Drosophila* and *Calliphora* (Trimarchi and Murphey, 1997).

In contrast to the striking morphological differences in b1 MN across genera, the overall morphology of tt MN is similar between *Drosophila* and *Calliphora*--tt MN has the same major branches in both. A possible difference between the two genera is that tt MN in *Calliphora* has many very thin, complex sub-branches splitting off from its major branches (Strausfeld et al., 1984). These fine sub-branches are not visible in *Drosophila* (Figure 5).

### Homologous haltere motoneurons in other insects

In Diptera, the hindwings have evolved into club-shaped halteres, which have their own reduced set of muscles. The morphology of the haltere MNs of *Drosophila* have not been described in detail before. We created driver lines targeting haltere MNs and used MCFO to visualize six haltere MNs.

It may be informative to compare the haltere MNs with the MNs of the muscles associated with the hindwings in other insects, such as Lepidoptera. Moths have two pairs of wings, but the hindwings are smaller than the forewings and are mechanically linked to them, similar to the mechanical linkage between wings and halteres in Diptera (Deora et al., 2015), and in contrast to the well-developed hindwings that flap out of phase with the forewings in more distantly related insects like locusts and dragonflies. Further, the projection patterns of sensory neurons from the fore- and hindwings of the moth are similar to those of campaniform sensilla from the wings and halteres of flies (Ando et al., 2011). Given these similarities, moth hindwing MNs and fly haltere MNs may be expected to have some similarities as well.

MNs of the hawkmoth hindwing DLMns have their dendrites in a dorsal layer of the nerve cord, with their cell bodies more ventral, the same as the haltere MNs of *Drosophila* (Ando et al., 2011). Like hDVMn, hi1 MN, and hi2 MN in *Drosophila*, the dendrites of the hindwing MNs in the hawkmoth appear to be close to where the axon exits the nerve cord. On the other hand, the somata of the hawkmoth hindwing MNs also appear to be close to the dendrites and axon on the anterior-posterior axis (Ando et al., 2011), while in *Drosophila*, the somata of haltere MNs are usually far posterior to their dendrites and axon. *Drosophila*’s haltere power muscle MN, hDVMn, thus has some anatomical similarities but also differences with hawkmoth hindwing power muscle MNs.

### Homologous interneurons in other insects

A small subset of dorsal VNC interneurons isolated from our collection could be matched to interneurons from studies in other insects based on their morphologies.

One of the interneurons targeted by our splits is the peripheral synapsing interneuron (PSI). The PSI is important for escape, along with the tt MN that triggers jumping and the DLMns that drive wing depression. The PSI is unique in that its axon terminals are not within the central nervous system, but rather its synaptic output is in the peripheral nerve (King and Wyman, 1980). Signals from the PSI need not travel through the dendritic arbors of the DLMns but can immediately activate the fibers of DLMns within the peripheral nerve, thus producing a much more rapid effect, in an escape behavior in which reactions must be as fast as possible. The PSI has been previously described in *Calliphora* (Strausfeld et al., 1984). However, the anatomy of the PSI is very different between the two genera. The PSI of *Drosophila* has only medial dendrites within the VNC, while the PSI of *Calliphora* also has lateral dendrites. The dendrites of the *Drosophila* PSI project fairly far anteriorly into T1 and posteriorly. In contrast, the dendrites of *Calliphora* PSI are restricted to a narrow area on the anterior-posterior axis. This is surprising, considering that other parts of the escape circuit, the DLMns and giant fiber have extremely similar morphology in *Drosophila* and *Calliphora*, and that tt MN also has more similar anatomy in *Drosophila* and *Calliphora*.

Another morphologically distinct interneuron population are the contralaterally projecting haltere interneurons (cHINs), which receive input in the haltere campaniform sensilla and are coupled to neck motoneurons; eight to ten of these neurons have been identified in *Calliphora* (Strausfeld and Seyan, 1985). We visualized four types of similar cHINs: IN08Ba1_T3-IN08Ba4_T3 in our nomenclature. All of these *Drosophila* cHINs have different morphology than *Calliphora* cHINs. Different haltere interneurons such as cHIN types may receive input from different populations of sensilla and thus respond to different types of haltere motion (Yarger and Fox, 2018). However, the dendrites where cHINs receive input in the haltere neuropil have extremely similar morphology in *Calliphora* and *Drosophila*. On the other hand, in the neck neuropil, *Calliphora* cHINs have many fine, thin branches. In contrast, each *Drosophila* cHIN innervates the neck neuropil with just one or two varicose terminals. This may suggest that the *Drosophila* cHINs have different connections in the neck neuropil than *Calliphora* cHINs.

cHINs can be distinguished as n-cHINs that innervate the neck neuropil (Strausfeld and Seyan, 1985) versus w-cHINs that innervate the wing neuropil (Trimarchi and Murphey, 1997). Both cHIN types are present in *Calliphora* (Hengstenberg, 1991). In our interneuron collection, IN08Ba1_T3–IN08Ba4_T3 are n-cHINs, while IN08Bb1_T3, IN08Bc1_T3, IN08Bb2_T3, IN07Bh1_T3, IN07Bh2_T3, and w-cHINa1_T2–w-cHINa4_T2 are w-cHINs. In *Drosophila,* some w-cHINs—possibly IN08Bb1_T3, IN08Bc1_T3, and IN08Bb2_T3—are coupled to b1 MN (Trimarchi and Murphey, 1997).

### Developmental origins of VNC neurons

As the insect analog of the vertebrate spinal cord, the VNC offers the potential for investigating the principles governing development-function links with wide applicability. The motor control systems of the spinal cord include highly diverse populations of interneurons which are organized into cardinal classes based on their common ontogeny and function (Arber, 2012, Grillner and Jessell, 2009). Similarly, the majority of VNC cells can be assigned to a specific hemilineage, with cells in a given hemilineage sharing similar functional and anatomical characteristics (Harris et al., 2015, Lacin et al., 2019, Shepherd et al., 2019).

In this study, we annotated the hemilineage identities of all ventral unpaired median neurons and interneurons targeted in our driver line collection using single-cell MCFO images (Figures 15-17 and supplements). Moreover, because hemilineage organization is shared across insect species (Shepherd et al., 2019), results obtained from the study of our driver lines will inform studies of all other insect nervous systems. Such cross-taxa comparisons have the potential to elucidate general principles of motor system organization, and our driver line collection offers a unique toolkit to pursue such questions.

In aggregate, our interneuron collection contains cells from 14 VNC hemilineages: 0A, 2A, 3B, 5B, 6A, 6B, 7B, 8B, 11B, 12A, 17A, 18B, 19A, and 19B. Consistent with our goal of producing driver lines targeting wing control circuitry, the hemilineages represented in our interneuron collection primarily arborize either solely in dorsal VNC neuropil regions (2A, 6A, 6B, and 11B) or bridge connections between ventral and dorsal VNC neuropils (0A, 3B, 5B, 7B, 8B, 12A, 17A, 19A, and 19B) as might be involved in take-off behaviors (Shepherd et al., 2019). By analyzing single-cell images, we found that many hemilineages which innervate both dorsal and ventral regions of the VNC include neurons which arborize only in the dorsal VNC. The hemilineages which have been reported to connect the dorsal and ventral VNC do contain interneurons which connect different layers of the VNC, but they also include solely dorsal interneurons, and presumably solely ventral interneurons which are not included in this study. Thus, our sparse driver lines and single-cell analysis revealed diversity within hemilineages which had not previously been reported. In addition, activation experiments from previous studies have linked the hemilineages in our interneuron collection to several behaviors (Harris et al., 2015), including wing buzzing (2A, 7B, 11B, 12A, 18B), wing waving (3B, 12A), takeoff (7B, 11B, 18B), postural changes (3B, 5B, 8B), and uncoordinated leg movements (6A, 6B). The confluence of anatomical and behavioral evidence suggests that the hemilineages represented in our interneuron collection are likely to represent discrete populations involved in the control of wing behaviors; future investigations involving these interneurons can leverage this hemilineage identification to probe their functional significance.

### Compatibility with the *Drosophila* neurobiology toolkit

The resources and analytic approaches presented in the study will be greatly complemented by the burgeoning set of genetic tools, experimental techniques, and novel data sets available for the investigation of the *Drosophila* central nervous system. As alluded to above, the use of driver lines in combination with tools that allow transient or chronic manipulation of genetically targeted neurons will be an invaluable resource for linking neuronal activity in the VNC to behavior (Simpson and Looger, 2018, Venken et al., 2011). In cases where our driver lines target multiple cell types, techniques for systematically reducing driver line expression patterns will help hone the results of such efforts (Isaacman-Beck et al., 2020). Similarly, expressing new, advanced calcium indicators in the cells targeted by our driver lines (Zhang et al., 2023) and utilizing recently developed techniques for monitoring neuronal activity in the VNC (Chen et al., 2018, Shiozaki et al., 2024), will facilitate a greater understanding of the temporal dynamics of VNC circuitry. Genetic techniques for synaptic tracing in combination with our driver lines (Cachero et al., 2020, Talay et al., 2017) or leveraging the matching between our segmented light microscopy images and electron microscopy reconstructions of the fly VNC (Takemura et al., 2023) will allow circuit-level analysis. Finally, the increasing number of other such driver line sets can be used in combination with the one presented here to gain a richer understanding of how neuronal circuits function at the scale of the full CNS.

## Materials and Methods

### Fly Stocks

Split-GAL4 driver lines stocks were generated as described below and maintained as homozygous stocks. For experiments, the driver lines were crossed to one of the following effector lines: 20XUAS-CsChrimson-mVenus trafficked in attP18 for activation experiments and screening VNC expression; w;UAS-Kir2.1-GFP(III) or w+ DL; DL; pJFRC49-10XUAS-IVS-eGFPKir2.1 in attP2 (DL) for courtship song silencing experiments with Kir; w+; UAS-TNTe for courtship song silencing experiments with tetanus; or pJFRC7-20xUAS-IVS-mCD8::GFP in attP2 (Pfeiffer et al., 2010) for imaging innervation of muscles with phalloidin staining. The progeny from these crosses were reared on cornmeal-based food in a 25°C incubator with a 16:8 light cycle.

### Generation of split Gal4 lines targeting VNC neurons

14,284 confocal stacks of the VNCs of female Janelia Generation1 GAL4 flies (Jenett et al., 2012, Meissner et al., 2023) were registered to a common template. For alignment, confocal stacks of the VNC were converted to an 8-bit nrrd file format. The reference channel, which used the nc82 antibody against Bruchpilot protein (Developmental Studies Hybridoma Bank, deposited by Buchner, E.) to label neuropils, was used to normalize contrast across samples. Each stack was rotated so that the anterior-poster axis was vertical, and the reference channel was aligned to a template by nonrigid warping using the computational morphometry toolkit (Rohlfing and Maurer, 2003). VNC images were aligned to the JRC2018_VNC_UNISEX template (Bogovic et al., 2020). The signal channel was then transformed using the warped mesh.

Each of these aligned stacks was then used to generate a maximum intensity projection (MIP), in which the color of the signal encoded its depth in the dorsal-ventral axis, called a Color MIP. Masks were drawn around neurites of targeted wing motoneurons in these Color MIPs and used to search the entire collection for a match using a custom-written FIJI plugin (Schindelin et al., 2012, Namiki et al., 2018, Otsuna et al., 2018). This process generally narrowed down the number of possible matching expression patterns to a few hundred.

Split-GAL4 driver lines were made by crossing split-GAL4 activation domain (AD) and DNA binding domain (DBD) hemidriver fly lines, from the collections described in (Dionne et al., 2018) and (Tirian and Dickson, 2017). Over the course of this project, we selected 1,658 AD/DBD split GAL4 intersections to screen. Split intersections were screened by crossing them with 20XUAS-CsChrimson-mVenus trafficked in attP18, dissecting out the central nervous system of 3–10-day old female offspring, immunolabeling against mVenus and nc82, embedding the preparations in DPX, viewing them with a Zeiss fluorescent microscope and then imaging selected preparations with a confocal microscope with the 20x objective. Most of the split intersections either had broad expression or dim expression. We selected 195 of the splits to be stabilized by making them homozygous for the AD and DBD transgenes. For stabilized splits, we used multi-color flip out to stochastically label individual neurons. Images of CsChrimson-mVenus expression patterns and of multi-color flip out were aligned and used to produce more color MIPs which could be used to search for sparser splits. Images were segmented using Amira or VVDViewer (https://github.com/JaneliaSciComp/VVDViewer/). In cases where MCFO of a split line produced sparse enough images that a cell type could be identified based on its morphology, but no segmentable single-cell image of that cell type was produced in this project, we examined the MCFO data for the AD and DBD hemidriver lines used to build the split-GAL4 driver line, and if the hemidriver MCFO data included a segmentable single-cell image of that cell type, we used that as the example image in this paper (i.e. 14E09 for DLMna/b, VT008474 for ps1 MN, 22E12 for IN06Bd1_T2, 26A08 for IN12Aa1_T2, 50G08 for IN02Ab1_T3, VT014604 for IN06Ag3_T1, 22E12 for IN06Bd2-4_T2, IN18Bc1_T3 for HBI002).

### Polarity labeling to visualize output sites

After split intersections were stabilized, they were crossed with pJFRC51-3xUAS-Syt::smGFP-HA in su(Hw)attP1; pJFRC225-5xUAS-IVS-myr::smGFP-FLAG in VK00005. The central nervous system of 3-10 day old male offspring was dissected and immunolabeled against nc82, HA and FLAG. Labeling protocol and additional details are available at https://www.janelia.org/project-team/flylight/protocols.

### Phalloidin Immunohistochemistry and Confocal Imaging

In order to visualize which muscles were innervated by motoneurons with split GAL4 expression, we labeled the muscles with phalloidin, a mushroom toxin which binds to f-actin. Splits were crossed with pJFRC7-20xUAS-IVS-mCD8::GFP in attP2 on cornmeal-based food and kept at 25°C. When the offspring were 3-4 days old, males were chilled on ice to anesthetize them, dunked in ethanol to remove the wax on the cuticle, and dipped in phosphate-buffered saline (PBS) twice to remove any traces of ethanol. The head, abdomen, wings and legs of each fly were removed using forceps and scissors. Then the fly thoraxes were fixed in 1% paraformaldehyde (Fisher Scientific) in PBS overnight at 4°C while nutating. The thoraxes were washed in 0.5% PBS with Triton-X (PBT) four times for 10-15 minutes each while nutating. Then the thoraxes were embedded in 7% SeaKem LE agarose (Fisher Scientific) which was hardened at 4°C for 60 minutes. The thoraxes were each cut in half along the midline by hand with a stiff single-edged razor blade (Personna). The preparations were blocked in PBS with 1% Triton-X, 0.5% DMSO, 3% normal goat serum, with a pinch of NaN_3_, and a pinch of escin. 1:200 bovine hyaluronidase type IV-S (Sigma Aldrich) was also added to break down connective tissue and thus allow better penetration of the antibody. The preparations were blocked for 60 minutes before being incubated in 1:100 phalloidin conjugated with Alexa Fluor 633 (Life Tech) along with 1:50 anti-nc82 in mouse (Univ of Iowa) and 1:1000 anti-GFP in rabbit diluted in the blocking solution. The preparations were left in the primary antibody and phalloidin solution for 5 days at 4°C.

After incubation in the primary antibody, preparations were washed four times for 10-15 minutes in 0.1% PBT. The preparations were blocked again in the same blocking solution but without hyaluronidase for 30 minutes before being incubated with 1:500 goat anti-mouse Cy3 and 1:500 goat anti-rabbit Alexa 488 for 2-3 days at 4°C. Preparations were washed overnight in 0.1% PBT, then fixed in 2% paraformaldehyde in PBS at 4°C, then washed four times 10-15 minutes. The preparations were cleared in a graded series of glycerol for 30 minutes each: 5% glycerol, 10% glycerol, 20% glycerol, 30% glycerol, 50% glycerol, 65% glycerol. Then the preparations were left in 80% glycerol overnight at 4°C. Preparations were dehydrated in a graded series of ethanol for 30 minutes each: 30% ethanol, 60% ethanol, 90% ethanol, then 100% ethanol three times for 15 minutes. The preparations were put into 50% methyl salicylate (Sigma Aldrich) for 30 minutes, then 100% methyl salicylate. To remove any remaining glycerol, the preparations were rinsed in ethanol and then returned to 100% methyl salicylate. The preparations were left in 100% methyl salicylate overnight at 4°C. The preparations were mounted on cover slips in methyl salicylate. Preparations were imaged with a 10x air objective using the Zeiss 880 NLO upright confocal microscope or the Zeiss 710 confocal microscope. Images were processed using Fiji and VVDViewer.

### Courtship song silencing experiments

Courtship behavior assays and analysis were performed as described in Arthur et al., 2013. Motoneuron splits were crossed with w+; UAS-TNTe or w;UAS-Kir2.1-GFP(III) or w+ DL; DL; pJFRC49-10XUAS-IVS-eGFPKir2.1 in attP2 (DL) (ID# 1117481) on semi-defined food (food described in (Backhaus et al., 1984)). As a control, “blank” split-GAL4 lines created from the same Gen1 GAL4 collection but with no central nervous system expression (SS01055 or SS01062) were also crossed with the same reporter lines on power food. Virgin male offspring were collected and socially isolated in small glass vials of power food. To entrain the circadian rhythm of the flies, for at least 48 hours before each experiment the males were kept in an incubator in which the lights turned on daily within two hours before the time of the experiment (i.e. if the incubator’s lights turned on at 9 am every day, the experiment would be done before 11 am). Each male was placed in his own courtship arena, along with a w1118 virgin female who had been collected the day before. An aspirator was used to insert the flies into the arena; they were not anesthetized. The arenas were within an acrylic box on air table to reduce outside noise and vibration. An acrylic platform held each chamber over a pressure gradient microphone. The temperature and humidity were recorded simultaneously using a SHT75 humidity sensor (Sensirion). Song was recorded for 30 minutes. Control and experimental flies were always recorded simultaneously.

### Flight activation experiments

Flight activation experiments were performed as in (Suver et al., 2016). Motoneuron splits, as well as the SS01062 blank split negative control, were crossed with 20XUAS-CsChrimson-mVenus trafficked in attP18 on food with 1:250 retinal and kept in the dark. Female offspring were tested when 3-8 days old. Flies were cooled and tethered by gluing each fly to a wire with Loctite^®^ 3972^TM^ UV-activated cement (Henkel). The tether was positioned on the anterior dorsal thorax on each fly. An arena of 470 nm blue LEDs was used to display closed-loop visual stimuli in the form of vertical stripe patterns, as described in in Kim et al., 2017. Fly behavior during these experiments was measured using three IR-sensitive cameras recording at 100 frames per second, as well as a Wingbeat Tachometer (IO Rodeo) for measuring wingbeat frequency.

A 617 nm red fiber-coupled LED (Thorlabs) suitable for optogenetic activation of the red-shifted channelrhodopsin, CsChrimson, was positioned below the fly, aimed at the ventral thorax and turned on in 0.1 s pulses with 6 s between each pulse. Each closed loop experiment comprised either 20 or 50 consecutive trials. In open loop experiments, the LED arena displayed closed loop visual stimuli for 20 trials, followed by open loop stimuli in random order, repeated 5 times, followed by closed loop stimuli for 10 trials. The open loop stimuli were horizontal expansion, clockwise rotation, counterclockwise rotation, and contraction of the stripes. The intensity of the red light was set to 1.9 milliW/mm^2^ for each experiment.

Kinefly, a machine vision system for quantifying flight behavior in real time (Suver et al., 2016), was used to extract the following wing kinematic parameters from the camera views and Wingbeat Tachometer: stroke amplitude, forward deviation angle, backward deviation angle, and wingbeat frequency.

### Behavioral Data Analysis

Analysis of behavioral data from tethered flight and courtship experiments was performed in MATLAB. For tethered flight experiments, noise in the single-trial wing kinematics was removed using a median-filter with a 15 ms window. To detect changes resulting from the optogenetic perturbation, the mean values of each kinematic variable in the 1 second window prior to an LED pulse were subtracted from individual traces. The filtered, mean-subtracted data was then averaged across trials for each fly/condition, as well as across the left and right wing, since the genetic lines used target the motor neurons bilaterally. The maximum change in the fly-averaged kinematics during the optogenetic stimulus period was then compared to control using the Wilcoxon rank sum test with Bonferroni method multiple comparison correction, as in Figure 10D-G. The 95% confidence intervals for per-fly mean wing kinematics, as in Figure 10C, were calculated by a 500-sample bootstrap procedure.

For courtship experiments, the FlySongSegmenter program (Arthur et al., 2013) was used to automatically segment raw sound recordings into periods of pulse song, sine song, or noise (non-singing). . Additionally, the identified song pulses were grouped into ‘fast’ or ‘slow’ mode pulses using the classifier described in (Clemens et al., 2018). The Wilcoxon rank sum test with Bonferroni method multiple comparison correction was used to identify statistical significance between experimental and control group flies. The per-genotype traces for pulse modes, as in Figure 11E, were calculated as the grand mean across flies, with 95% confidence intervals determined using a 500-sample bootstrap.

### Input/Output Identification

Inputs and outputs in previously defined neuropil regions (Court et al., 2020, Namiki et al., 2018) were identified for each of our interneurons. Post-synaptic dendrites (input sites) and pre-synaptic axonal terminals (output sites) were identified by their respective morphological differences using 63X magnification microscopy images. Post-synaptic dendrites have smooth endings, whereas axon terminals have a varicose shape (Namiki et al., 2018). For further confirmation of neurite type, split driver lines were crossed with pJFRC51-3xUAS-Syt::smGFP-HA in su(Hw)attPa in order to visualize the localization of synaptotagmin. Using this identification strategy, each individual neuron from the driver line collection was scored for the presence or absence of inputs and outputs in each of the 28 identified VNC neuropils (14 neuropil regions, each with an ipsilateral and contralateral half; see Figure 1C). For a given cell and neuropil region combination, the possible annotation values were: “D” (“Dendrite,” signifying input), “A” (“Axon,” signifying output), “M” (“Mixed,” signifying mixed inputs and outputs), and “P” (“Partitioned,” signifying spatially segregated inputs and outputs in the same neuropil region). Moreover, we distinguished between major and minor innervation of a neuropil region using upper and lower case letters, respectively, i.e. “A” would represent major output, while “a” would represent minor output.

### Identifying matching cells in the connectome

To identify matching volumes for our segmented cell images in the male adult VNC (MANC; (Takemura et al., 2023)) connectome database, we used the NeuronBridge web tool (Clements et al., 2024) to perform color depth MIP searches (Otsuna et al., 2018) of our segmented cell images against MANC images. These searches generated a list of possible matches for each cell, from which we manually selected the best overall match. We used hemilineage identity as an additional constraint on the search. Matches for the wing motoneurons in the MANC database were first reported in (Cheong et al., 2023); we include them here for completeness.

## Supporting information

STARMethods

## Data and Code Availability

All code needed to reproduce the analyses in this paper can be found at: https://github.com/samcwhitehead/dorsal_vnc_analysis

## Acknowledgments

This work was part of the Descending Interneuron Project Team and the Ventral Nerve Cord Consortium at Janelia Research Campus, part of the Howard Hughes Medical Institute. We received additional support from the Janelia Visiting Scientist Program. We thank the members of Janelia’s Invertebrate Shared Resource and Project Technical Resources, especially Gudrun Ihrke. We thank Anne von Philipsborn and Troy Shirangi for input on motoneuron identifications and Haluk Lacin for valuable discussions on hemilineages and IHC protocols. Discussions with Han Cheong and Fabienne Reh helped identify T2VUMs in images. We thank Elizabeth Kim for technical assistance and Ben J Arthur for analysis advice on the courtship song recordings. Patrick Breads and members of the Card Lab at Janelia provided technical assistance.

During this effort, the FlyLight Project Team included Gina DePasquale, Zack Dorman, Kaitlyn Forster, Theresa Gibney, Joanna Hausenfluck, Yisheng He, Kristin Hendersen, Jennifer Jeter, Lauren Johnson, Rachel Lazarus, Kelley Lee, Oz Malkesman, Geoffrey Meissner, Brian Melton, Scott Miller, Alexandra Novak, Alyson Petruncio, Jacquelyn Price, Sophia Protopapas, Susana Tae, Allison Vannan, Rebecca Vorimo, and Brianna Yarborough, with Steering Committee of Yoshinori Aso, Gwyneth Card, Barry Dickson, Reed George, Wyatt Korff, Gerald Rubin, and James Truman.

IC was supported by the National Institute of Health (NINDS, 1R01NS116595 and U01NS131438). MHD was supported by the National Institute of Health (NINDs, U19NS104655 and R01NS136988). KI was supported by the AXA Research Fund – AXA Chair: From genome to structure and by the German Science Foundation – NeuroNex Communication, Coordination, and Control in Neuromechanical Systems. KI and EE were supported by the the German Science Foundation Collaborative Research Center (CRC) 1451 – Key mechanisms of motor control in health and disease.

This article is subject to HHMI’s Open Access to Publications policy. HHMI scientists have previously granted a nonexclusive CC BY 4.0 license to the public and a sublicensable license to HHMI in their research articles.

---

**Figure 1—figure supplements 1-5: VNC expression patterns of split-GAL4 driver lines stabilized in this project.** Each split line was crossed with pJFRC51-3xUAS-Syt::smGFP-HA in su(Hw)attP1; pJFRC225-5xUAS-IVS-myr::smGFP-FLAG in VK00005. This figure shows the male expression pattern, except in lines which had no male expression, such as SS51528. Each VNC stack was aligned to the JRC 2018 VNC Unisex template. These maximum intensity projections are color coded to indicate the depth of each pixel. Blue represents the ventral layers of the VNC while red represents the dorsal layers. Panels are arranged by split number, except that splits targeting leg neurons which were serendipitously produced in this project are arranged in the last 18 panels.

**Figure 3—figure supplement 1: Power MN groups based on morphology in the VNC, illustrated using segmented MCFO images aligned to the unisex JRC2018 template.** (**A**) DLMna/b. (**B**) DLMnc-f. (**C**) The MNs innervating DVM 1 and 2. (**D**) The MNs innervating DVM 3. (**E**) Schematic depicting the muscle fibers innervated by the MNs in panels A-D in the medial (left) and lateral (right) thorax.

**Figure 9—figure supplement 1: Wing control MN groups based on morphology in the VNC, illustrated using segmented MCFO images aligned to the unisex JRC2018 template.** (**A**) Unilateral steering MNs with arborization mainly in the wing neuropil: b1 MN, iii3 MN, iii4 MN. (**B**) Unilateral steering MNs with arborization mainly in the wing neuropil: tpN MN, tp1 MN, tp2 MN, ps1 MN, hg2 MN, hg3 MN. (**C**) Steering MNs with approximately equal arborization in the wing neuropil and intermediate tectulum: b3 MN, i1 MN, i2 MN, hg1 MN. (**D**) Steering MNs with little or no arborization in the wing neuropil: tt MN, b2 MN, iii1 MN.

**Figure 11—figure supplement 1: Additional analyses of the effects of wing motoneuron silencing on courtship song.** (**A**) Courtship song statistics for motoneuron driver lines crossed to UAS-Kir2.1. Rows show inter pulse interval (top) and sine song carrier frequency (bottom) for each motoneuron driver line tested (columns). Open circles show per-fly measurements; bars and horizontal lines show interquartile range and population median, respectively. The number of flies per genotype is given in each figure axis. (**B**) Transition matrices giving median probabilities across flies for transitioning between the three song modes: pulse, sine, and null. Rows correspond to data from genetic control flies (top) and motoneuron-silenced flies (bottom). Orange asterisks indicate significant differences between genetic control and silenced groups. Significance assigned using the Wilcoxon rank sum test (***, p<0.001; **, p<0.01; *, p<0.05).

**Figure 13—figure supplement 1: Haltere MN groups based on morphology in the VNC, illustrated using segmented MCFO images aligned to the unisex JRC2018 template.** (**A**) Haltere MNs with unilateral, intersegmental arborization: hb1/2 MN. (**B**) Haltere MNs with bilateral arborization mainly in the haltere neuropil: hDVMn, hi1 MN, hi2 MN.

**Figure 16—figure supplement 1: Morphology of identified intrasegmental interneurons.** MCFO was used to separate interneurons. Each neuron is shown from three views—horizontal (top left), sagittal (bottom left), and transverse (bottom right)—along with neuron name and hemilineage identity (top right). Images taken from preparations of male flies, segmented to isolate individual neurons, and aligned to the JRC 2018 Unisex VNC template. The driver lines used are: SS42439, SS42442, SS42442, SS42442, SS42438, SS42439, SS48215, SS48215, SS48268, SS47192.

**Figure 16—figure supplement 2: Morphology of identified intrasegmental interneurons.** MCFO was used to separate interneurons. Each neuron is shown from three views—horizontal (top left), sagittal (bottom left), and transverse (bottom right)—along with neuron name and hemilineage identity (top right). Images taken from preparations of male flies, segmented to isolate individual neurons, and aligned to the JRC 2018 Unisex VNC template. The driver lines used are: 22E12 (no single-cell image appeared in the MCFO of the split GAL4 line targeting this cell, SS53421, so a generation 1 GAL4 line image was used), SS45830, SS45843, SS45843, SS45830, SS45843, SS25482, SS31309, SS46725 26A08 (no single-cell image appeared in the MCFO of the split GAL4 line targeting this cell, SS48272, so a generation 1 GAL4 line image was used).

**Figure 16—figure supplement 3: Morphology of identified intrasegmental interneurons.** MCFO was used to separate interneurons. Each neuron is shown from three views—horizontal (top left), sagittal (bottom left), and transverse (bottom right)—along with neuron name and hemilineage identity (top right). Images taken from preparations of male flies, segmented to isolate individual neurons, and aligned to the JRC 2018 Unisex VNC template. The driver lines used are: SS48272, SS48272, SS49799, SS49766, SS42438, SS40783, SS42442, SS46675, SS46675, SS40783.

**Figure 16—figure supplement 4: Morphology of identified intrasegmental interneurons.** MCFO was used to separate interneurons. Each neuron is shown from three views—horizontal (top left), sagittal (bottom left), and transverse (bottom right)—along with neuron name and hemilineage identity (top right). Images taken from preparations of male flies, segmented to isolate individual neurons, and aligned to the JRC 2018 Unisex VNC template. The driver lines used are: SS42438, SS42447, SS49779, SS51830, SS37306, SS52389, SS52389, SS52389, SS52389, SS52389.

**Figure 16—figure supplement 5: Morphology of identified intrasegmental interneurons.** MCFO was used to separate interneurons. Each neuron is shown from three views—horizontal (top left), sagittal (bottom left), and transverse (bottom right)—along with neuron name and hemilineage identity (top right). Images taken from preparations of male flies, segmented to isolate individual neurons, and aligned to the JRC 2018 Unisex VNC template. The driver lines used are: SS44002, SS44003, SS48268, SS47192, SS37233, SS20796, SS20796, SS49041, SS40778, SS54506.

**Figure 16—figure supplement 6: Morphology of identified intrasegmental interneurons.** MCFO was used to separate interneurons. Each neuron is shown from three views—horizontal (top left), sagittal (bottom left), and transverse (bottom right)—along with neuron name and hemilineage identity (top right). Images taken from preparations of male flies, segmented to isolate individual neurons, and aligned to the JRC 2018 Unisex VNC template. The driver lines used are: SS44314, SS54506, SS54506, SS54506, SS25502, SS54506, SS54506, 50G08 (no single-cell images were produced from the split GAL4 line targeting this cell, SS54506, so a generation 1 GAL4 image was used), SS49125, SS31274.

**Figure 16—figure supplement 7: Morphology of identified intrasegmental interneurons.** MCFO was used to separate interneurons. Each neuron is shown from three views—horizontal (top left), sagittal (bottom left), and transverse (bottom right)—along with neuron name and hemilineage identity (top right). Images taken from preparations of male flies, segmented to isolate individual neurons, and aligned to the JRC 2018 Unisex VNC template. The driver lines used are: SS25478, SS45611.

**Figure 16—figure supplement 8: Local-segmental VNC interneuron categories, illustrated using segmented MCFO images aligned to the unisex JRC2018 template.** (**A**) Local-segmental interneurons that mainly innervate the neck, AMN or intermediate tectulum. (**B**) Local-segmental interneurons with unilateral arborization mainly in the wing neuropil. (**C**) Local-segmental interneurons with bilateral arborization mainly in the wing neuropil. (**D**) Local-segmental interneurons with arborization mainly in the haltere neuropil.

**Figure 17—figure supplement 1: Morphology of identified intersegmental interneurons.** MCFO was used to separate interneurons. Each neuron is shown from three views—horizontal (top left), sagittal (bottom left), and transverse (bottom right)—along with neuron name and hemilineage identity (top right). Images taken from preparations of male flies, segmented to isolate individual neurons, and aligned to the JRC 2018 Unisex VNC template. The driver lines used are: SS25553, SS42079, SS42079, SS42079, SS49809, SS33489, SS42475, SS49784, SS49784, SS49784.

**Figure 17—figure supplement 2: Morphology of identified intersegmental interneurons.** MCFO was used to separate interneurons. Each neuron is shown from three views—horizontal (top left), sagittal (bottom left), and transverse (bottom right)—along with neuron name and hemilineage identity (top right). Images taken from preparations of male flies, segmented to isolate individual neurons, and aligned to the JRC 2018 Unisex VNC template. The driver lines used are: SS28361, SS46735, SS42464, SS42464, SS31289, SS31290, SS31290, SS49800, SS49778.

**Figure 17—figure supplement 3: Morphology of identified intersegmental interneurons.** MCFO was used to separate interneurons. Each neuron is shown from three views—horizontal (top left), sagittal (bottom left), and transverse (bottom right)—along with neuron name and hemilineage identity (top right). Images taken from preparations of male flies, segmented to isolate individual neurons, and aligned to the JRC 2018 Unisex VNC template. The driver lines used are: SS49807, SS29871, SS49779, 22E12 (no single cell images were produced from the split GAL4 driver lines targeting this cell, SS40456, SS42447 or SS53421, so a generation 1 GAL4 image was used), 22E12 (no single cell images were produced from the split GAL4 driver line targeting this cell, SS53421, so a generation 1 GAL4 image was used,) 22E12 (no single cell images were produced from the split GAL4 driver lines targeting this cell, SS53421, SS42447 or SS40456, so a generation 1 GAL4 image was used), SS40456, SS49799, SS49800, SS49800, SS40764.

**Figure 17—figure supplement 4: Morphology of identified intersegmental interneurons.** MCFO was used to separate interneurons. Each neuron is shown from three views—horizontal (top left), sagittal (bottom left), and transverse (bottom right)—along with neuron name and hemilineage identity (top right). Images taken from preparations of male flies, segmented to isolate individual neurons, and aligned to the JRC 2018 Unisex VNC template. The driver lines used are: SS40764, SS45385, SS45385, SS42446, SS42446, SS42446, SS48204, SS42446, SS48204, SS25553.

**Figure 17—figure supplement 5: Morphology of identified intersegmental interneurons.** MCFO was used to separate interneurons. Each neuron is shown from three views—horizontal (top left), sagittal (bottom left), and transverse (bottom right)—along with neuron name and hemilineage identity (top right). Images taken from preparations of male flies, segmented to isolate individual neurons, and aligned to the JRC 2018 Unisex VNC template. The driver lines used are: SS42499, SS42438, SS46295, SS49779, SS51830, SS49802, SS44005, SS40969, SS40969, SS29600.

**Figure 17—figure supplement 6: Morphology of identified intersegmental interneurons.** MCFO was used to separate interneurons. Each neuron is shown from three views—horizontal (top left), sagittal (bottom left), and transverse (bottom right)—along with neuron name and hemilineage identity (top right). Images taken from preparations of male flies, segmented to isolate individual neurons, and aligned to the JRC 2018 Unisex VNC template. The driver lines used are: SS51531, SS48709, SS51531, SS46260, SS46735, SS49777, 59F02 (no single-cell image appeared in the split GAL4 line targeting this cell, SS49777, so a generation 1 GAL4 image was used), SS42499, SS47214, SS47215.

**Figure 17—figure supplement 7: Morphology of identified intersegmental interneurons.** MCFO was used to separate interneurons. Each neuron is shown from three views—horizontal (top left), sagittal (bottom left), and transverse (bottom right)—along with neuron name and hemilineage identity (top right). Images taken from preparations of male flies, segmented to isolate individual neurons, and aligned to the JRC 2018 Unisex VNC template. The driver lines used are: SS44002, SS47215, SS49853, SS47215, SS49853, SS29535, SS29535. SS44276, SS31246, SS30816.

**Figure 17—figure supplement 8: Morphology of identified intersegmental interneurons.** MCFO was used to separate interneurons. Each neuron is shown from three views—horizontal (top left), sagittal (bottom left), and transverse (bottom right)—along with neuron name and hemilineage identity (top right). Images taken from preparations of male flies, segmented to isolate individual neurons, and aligned to the JRC 2018 Unisex VNC template. The driver lines used are: SS42050, SS49806, SS49806, SS47215, SS29602.

**Figure 17—figure supplement 9: Intersegmental VNC interneuron categories, illustrated using segmented MCFO images aligned to the unisex JRC2018 template.** (**A**) Intersegmental interneurons with their main arborization in the neck neuropil. (**B**) Intersegmental interneurons with mainly arborization in the AMN. (**C**) Intersegmental interneurons with arborization mainly in the tectulum. (**D**) Intersegmental interneurons with unilateral arborization mainly in the wing neuropil. (**E**) Intersegmental interneurons with bilateral arborization mainly in the wing neuropil. (**F**) Intersegmental interneurons with unilateral arborization mainly in the haltere neuropil. (**G**) Intersegmental interneurons with bilateral arborization mainly in the haltere neuropil.

**Figure 18—figure supplement 1-16: Comparison of interneuron single-cell light microscopy images and MANC electron microscopy matches.**

**Figure 18—figure supplement 17-20: Comparison of motoneuron single-cell light microscopy images and MANC electron microscopy matches.**

**Figure 18—figure supplement 21: Comparison of VUM single-cell light microscopy images and MANC electron microscopy matches.**

