## Supplementary material for "Single-cell type analysis of wing premotor circuits in the ventral nerve cord of *Drosophila melanogaster*": Supplimental figures

Supplemental figure 1.1

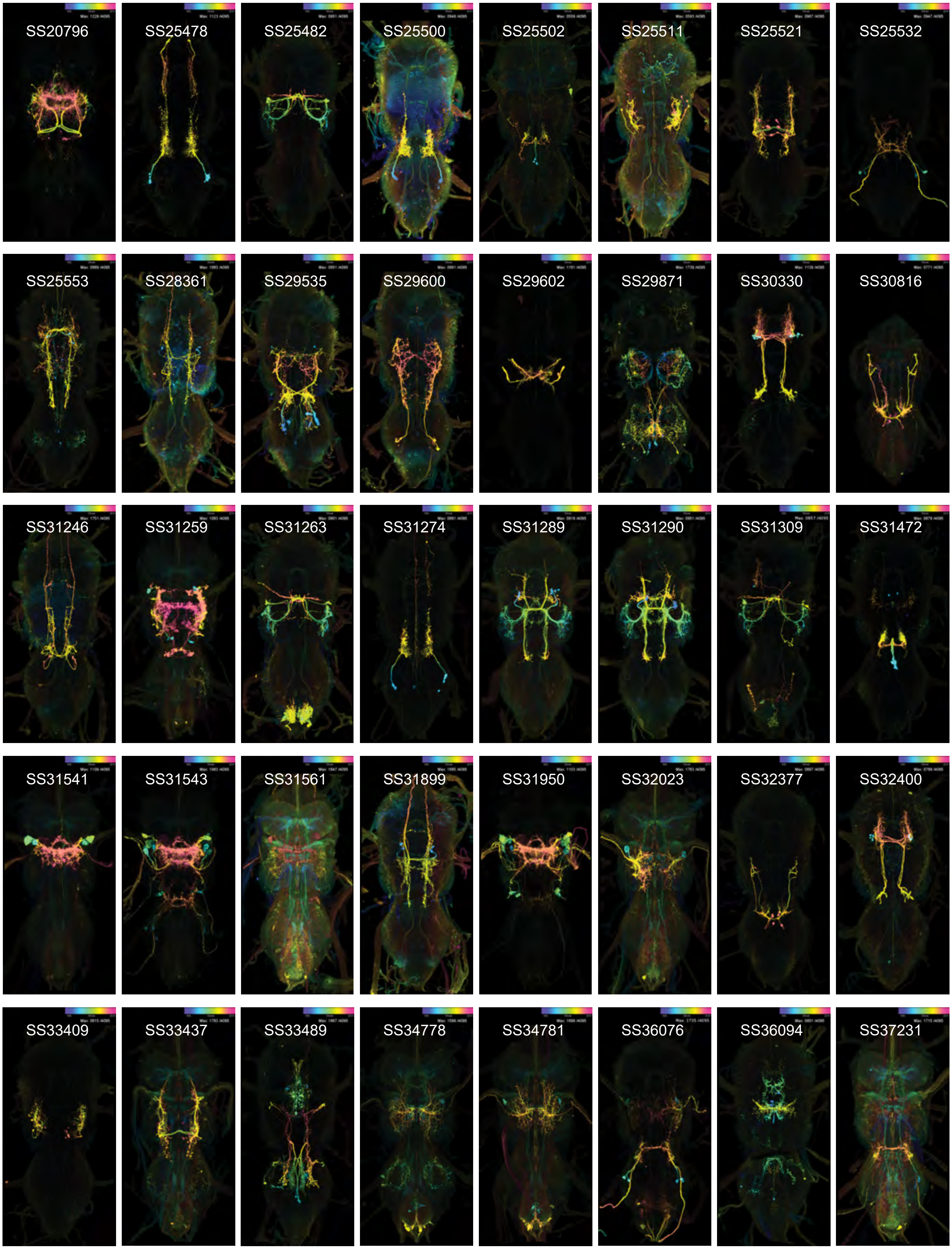

Supplemental figure 1.2

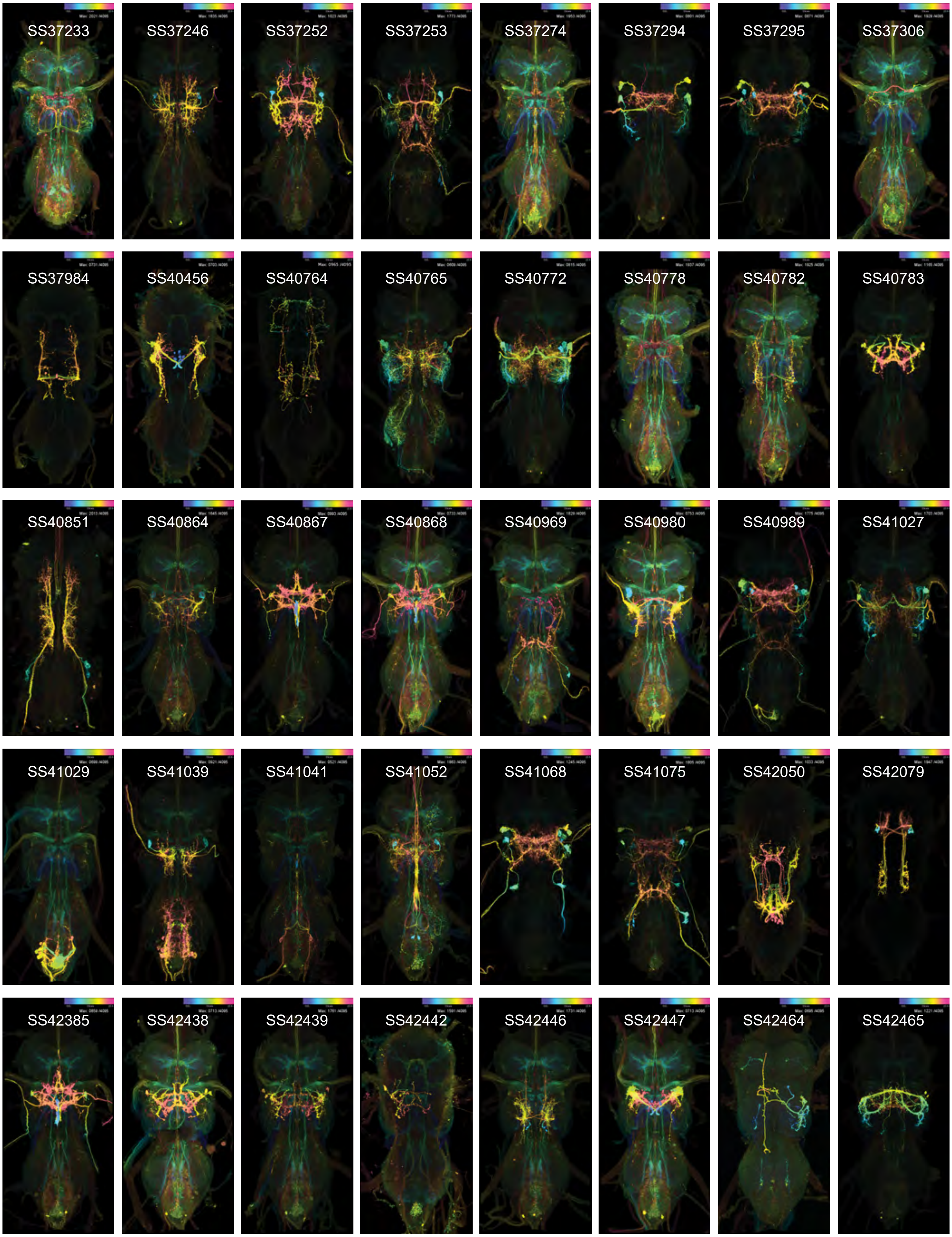

Supplemental Figure 1.3

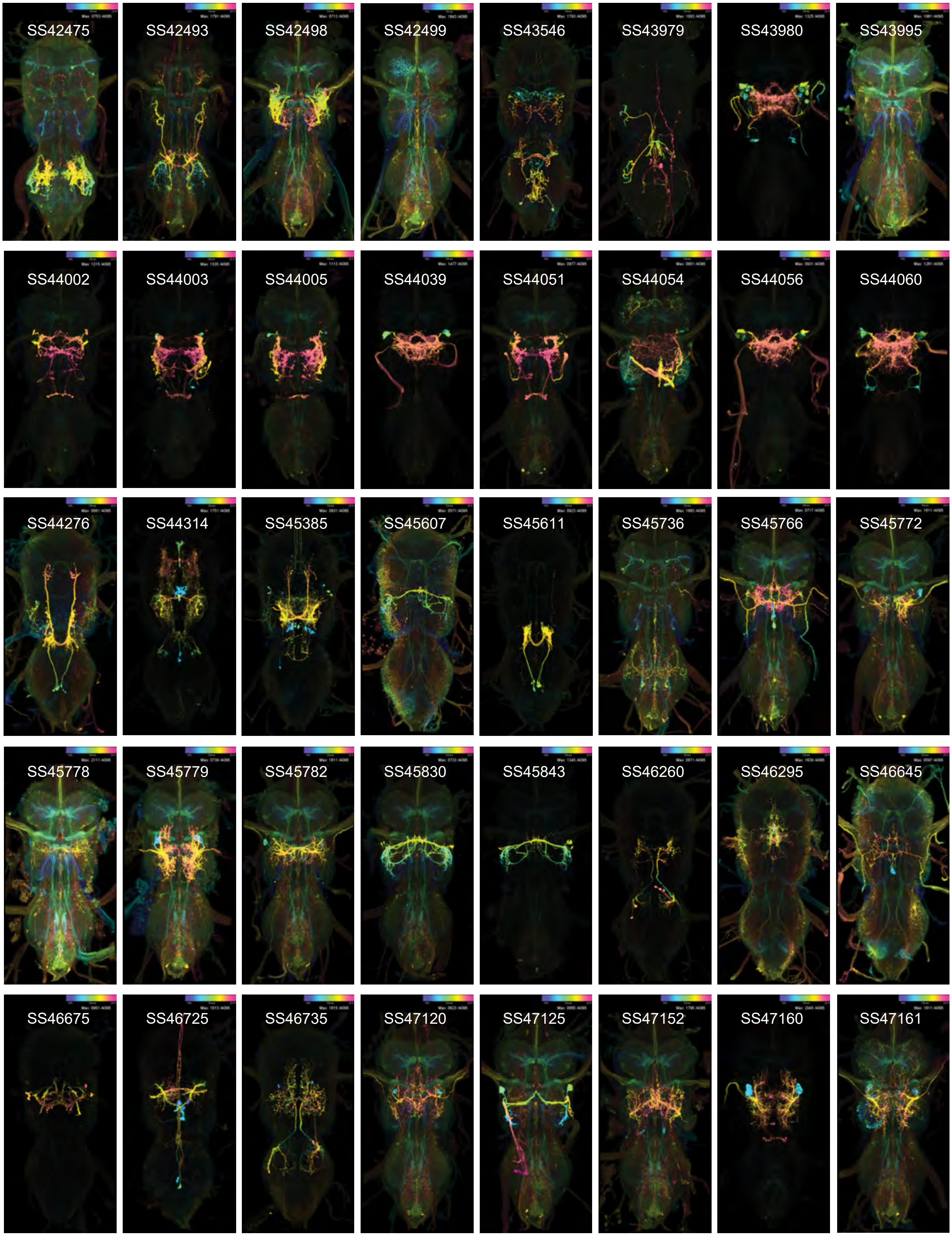

Supplemental Figure 1.4

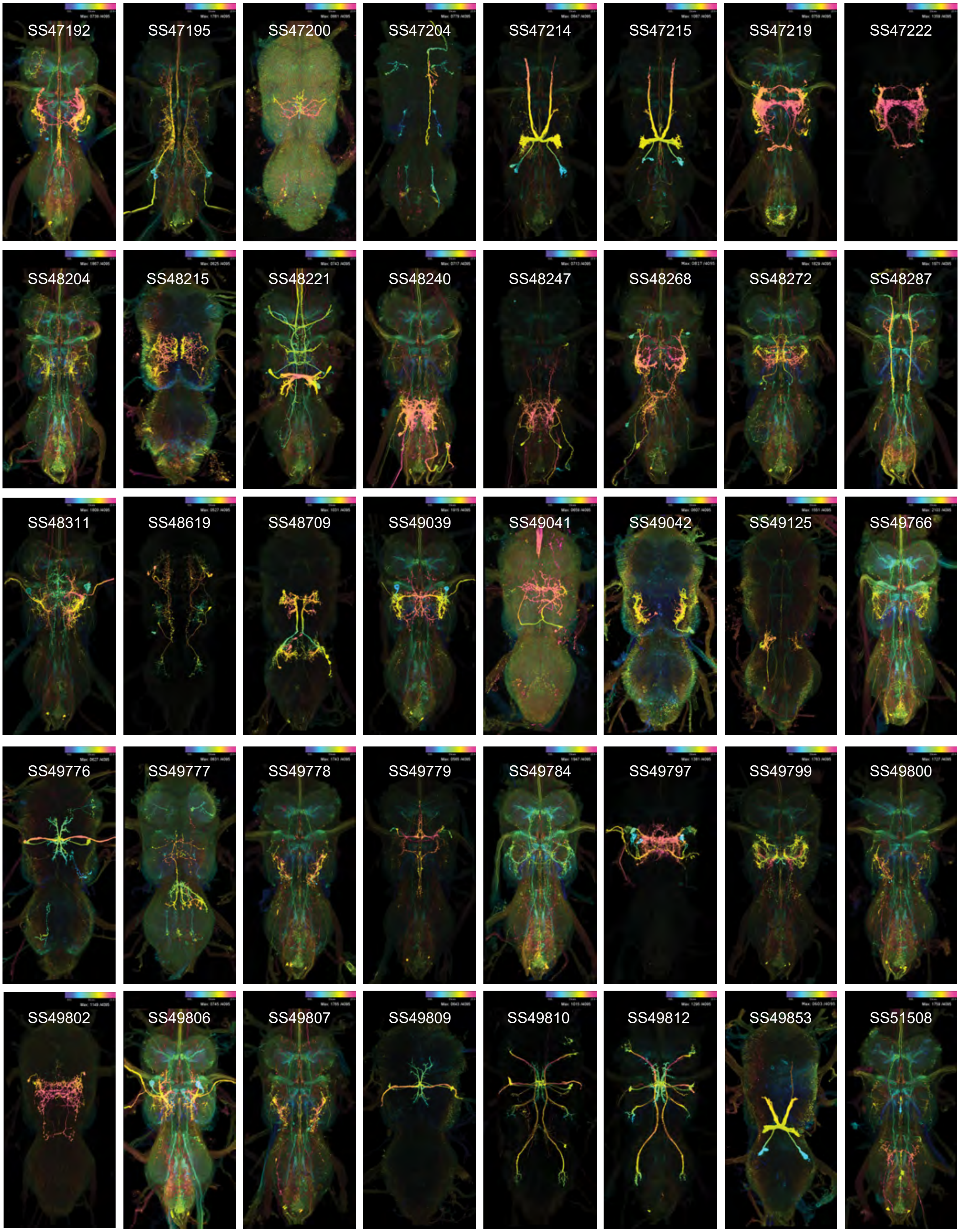

Supplemental Figure 1.5

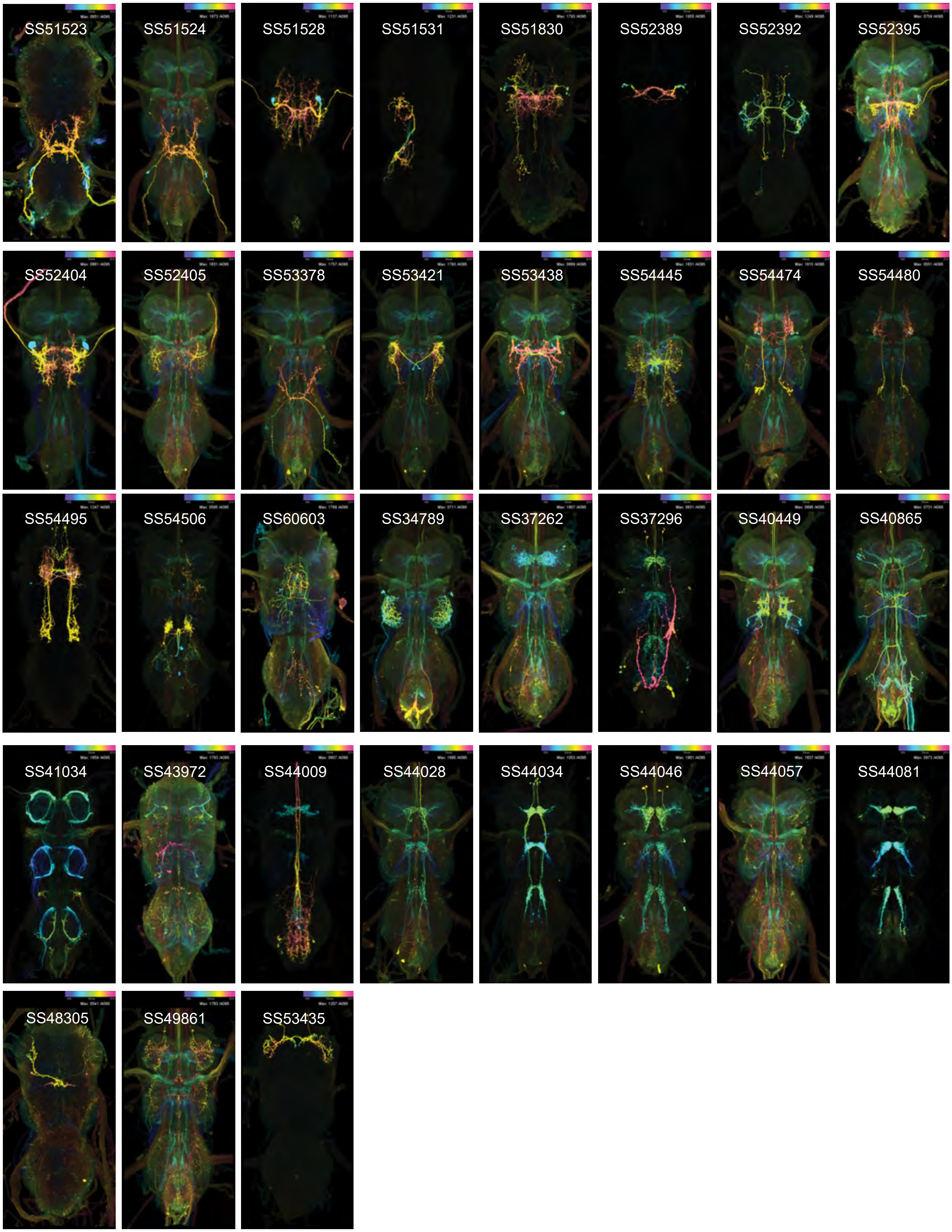

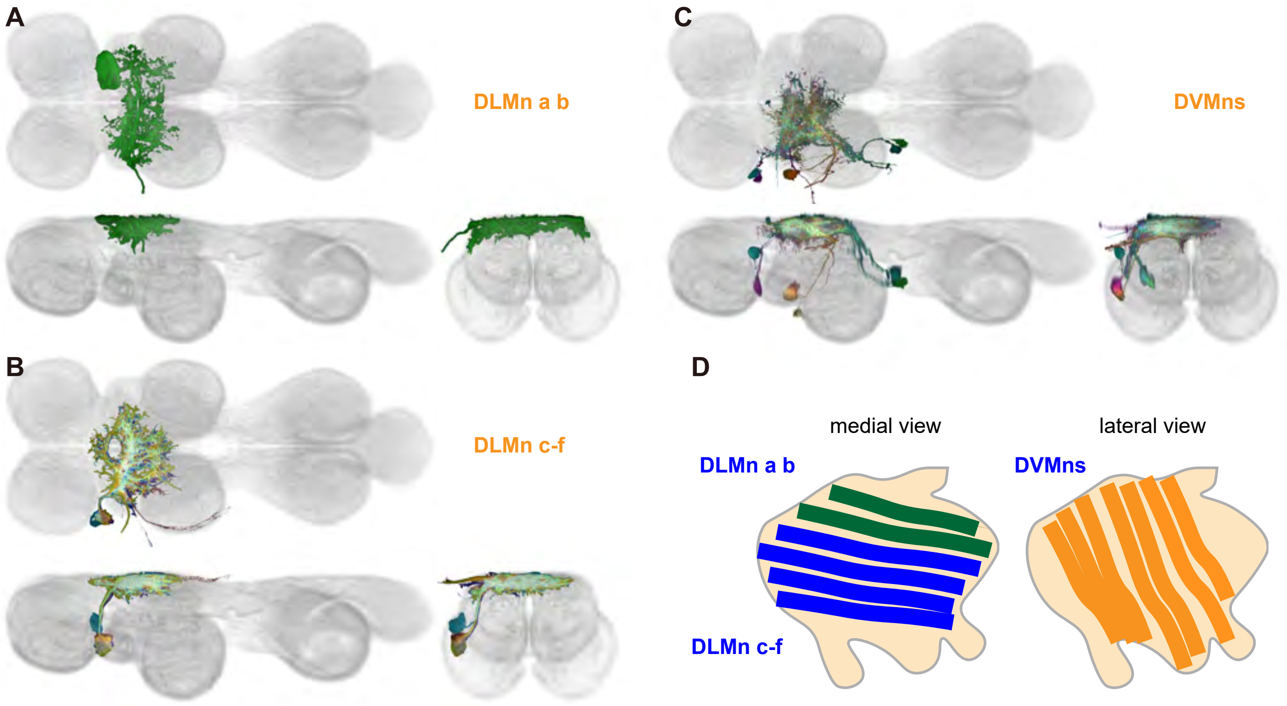
A
DLMn a b
B
DLMn c-f
C
DVMns
D
medial view
lateral view
DLMn a b
DVMns
DLMn c-f

A

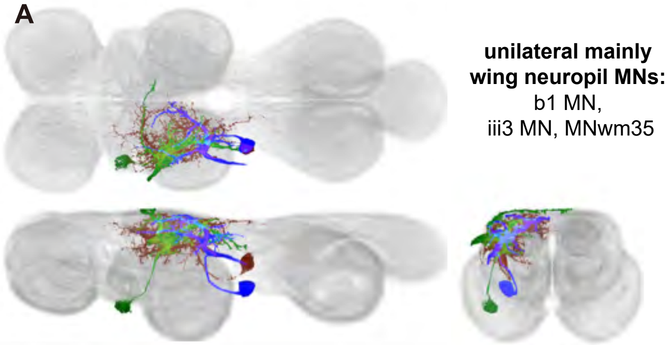
**unilateral mainly wing neuropil MNs:**
b1 MN,
iii3 MN, MNwm35

B

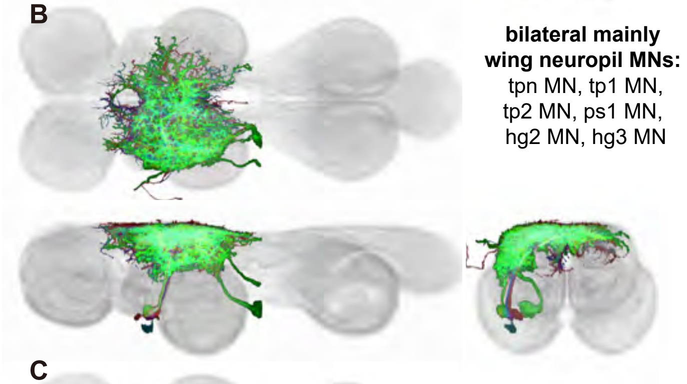
**bilateral mainly wing neuropil MNs:**
tpn MN, tp1 MN,
tp2 MN, ps1 MN,
hg2 MN, hg3 MN

C

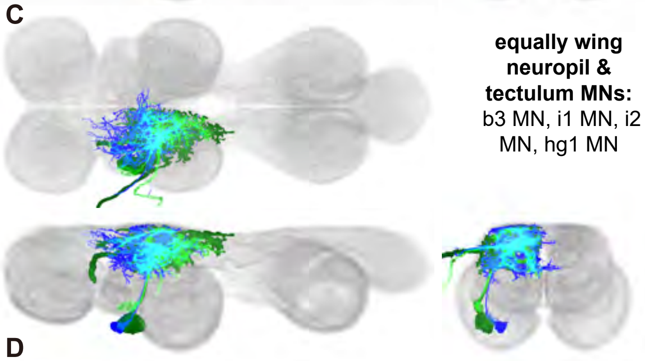
**equally wing neuropil & tectulum MNs:**
b3 MN, i1 MN, i2 MN, hg1 MN

D

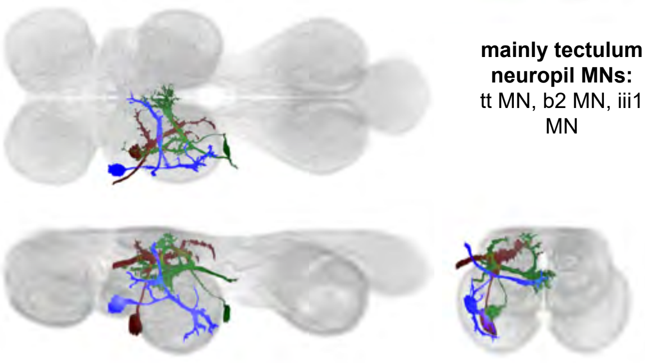
**mainly tectulum neuropil MNs:**
tt MN, b2 MN, iii1 MN

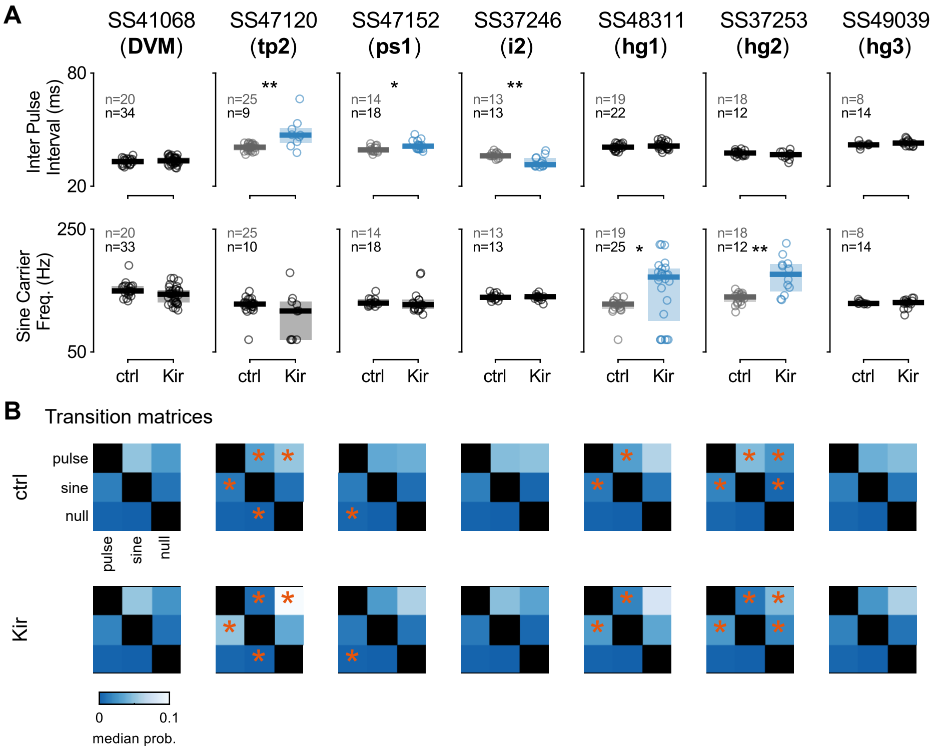
A
SS41068 (DVM)
SS47120 (tp2)
SS47152 (ps1)
SS37246 (i2)
SS48311 (hg1)
SS37253 (hg2)
SS49039 (hg3)
Inter Pulse Interval (ms)
80
20
n=20
n=34
n=25
n=9
**
n=14
n=18
*
n=13
n=13
**
n=19
n=22
n=18
n=12
n=8
n=14
Sine Carrier Freq. (Hz)
250
50
n=20
n=33
n=25
n=10
n=14
n=18
n=13
n=13
n=19
n=25
*
n=18
n=12
**
n=8
n=14
ctrl
Kir
B
Transition matrices
ctrl
Kir
pulse
sine
null
0
0.1
median prob.

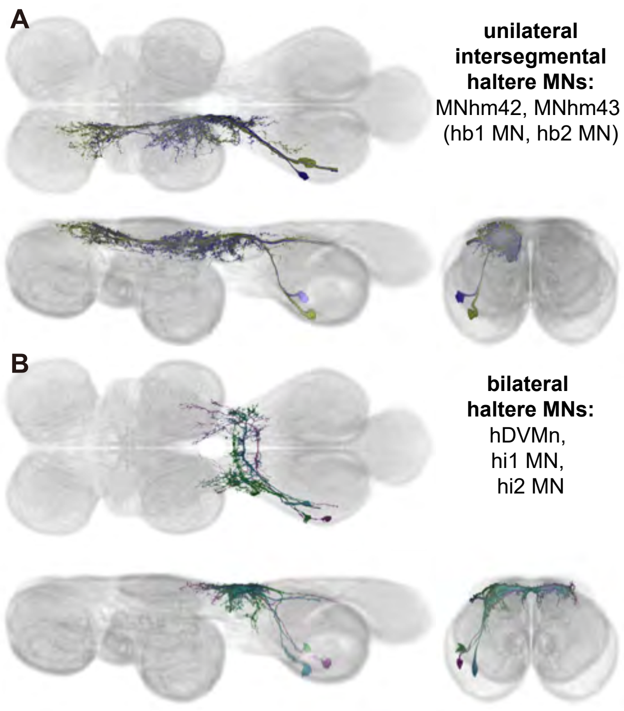
A
unilateral
intersegmental
haltere MNs:
MNhm42, MNhm43
(hb1 MN, hb2 MN)
B
bilateral
haltere MNs:
hDVMn,
hi1 MN,
hi2 MN

**A**

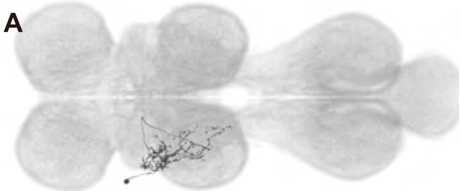

**IN17Ab1_T2**
MANC match:
IN17A055_T2
14661

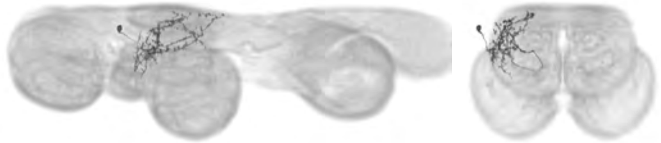

**B**

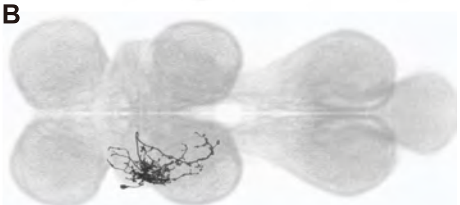

**IN17Aa1_T2**
MANC match:
IN17A049_T2
16009, 15908

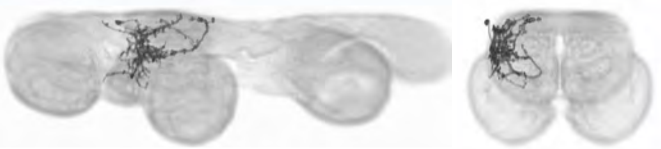

**C**

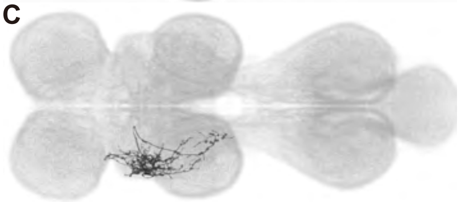

**IN17Aa2_T2**
MANC match:
IN17A049_T2
15754, 13989

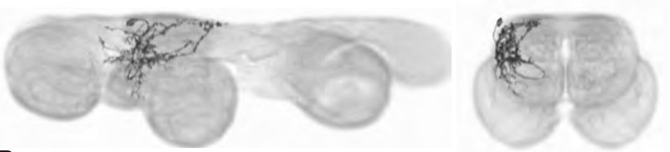

**D**

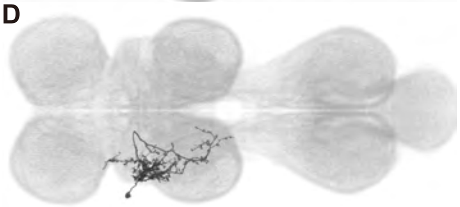

**IN17Ab2_T2 ***
MANC match:
IN17A055_T2
16664

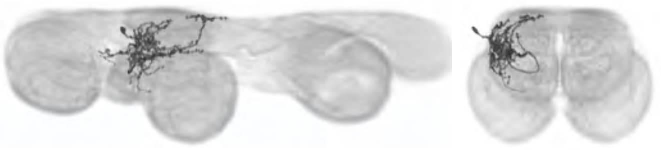

**E**

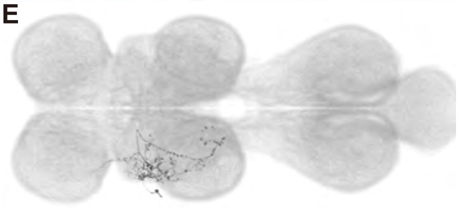

**IN17Ag1_T2**
MANC match:
IN17A107_T2
167001, 32912

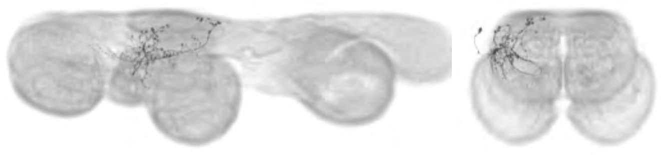

**F**

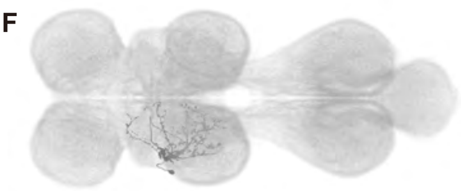

**dMS2a1_T2**
MANC match:
dMS2_T2
14309, 16277,
14425

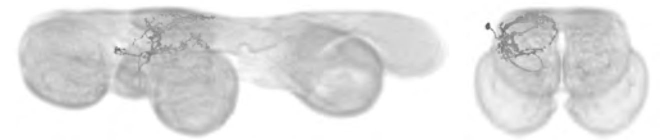

**G**

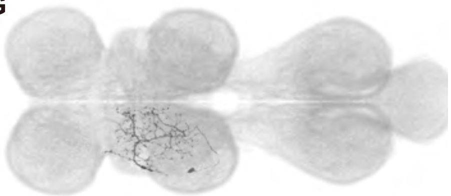

**IN11Bd1_T2**
MANC match:
IN11B013_T2
17679, 17561,
16851, 17159

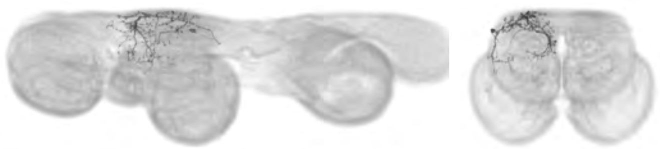

**H**

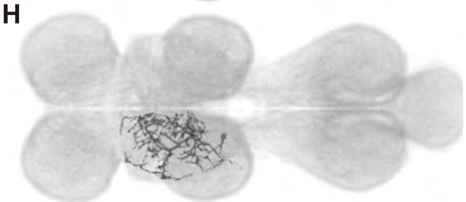

**IN11Bd2_T2**
MANC match:
IN11B013_T2
15175, 16088

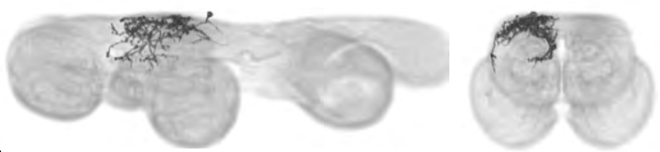

**I**

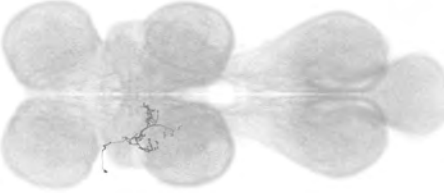

**IN03Bi1_T1**
MANC match:
IN03B077_T1
20763, 24769,
22336, 22643

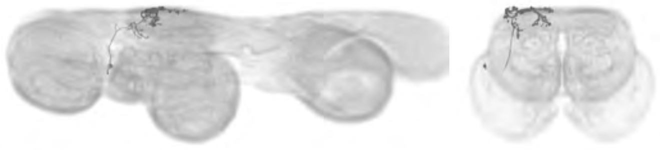

**J**

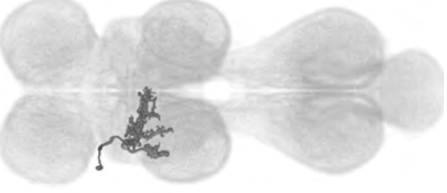

**IN03Bi2_T1**
MANC match:
IN03B077_T1
22846, 101541,
18054, 23168

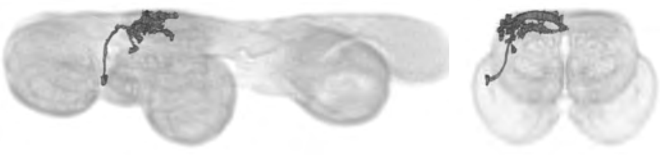

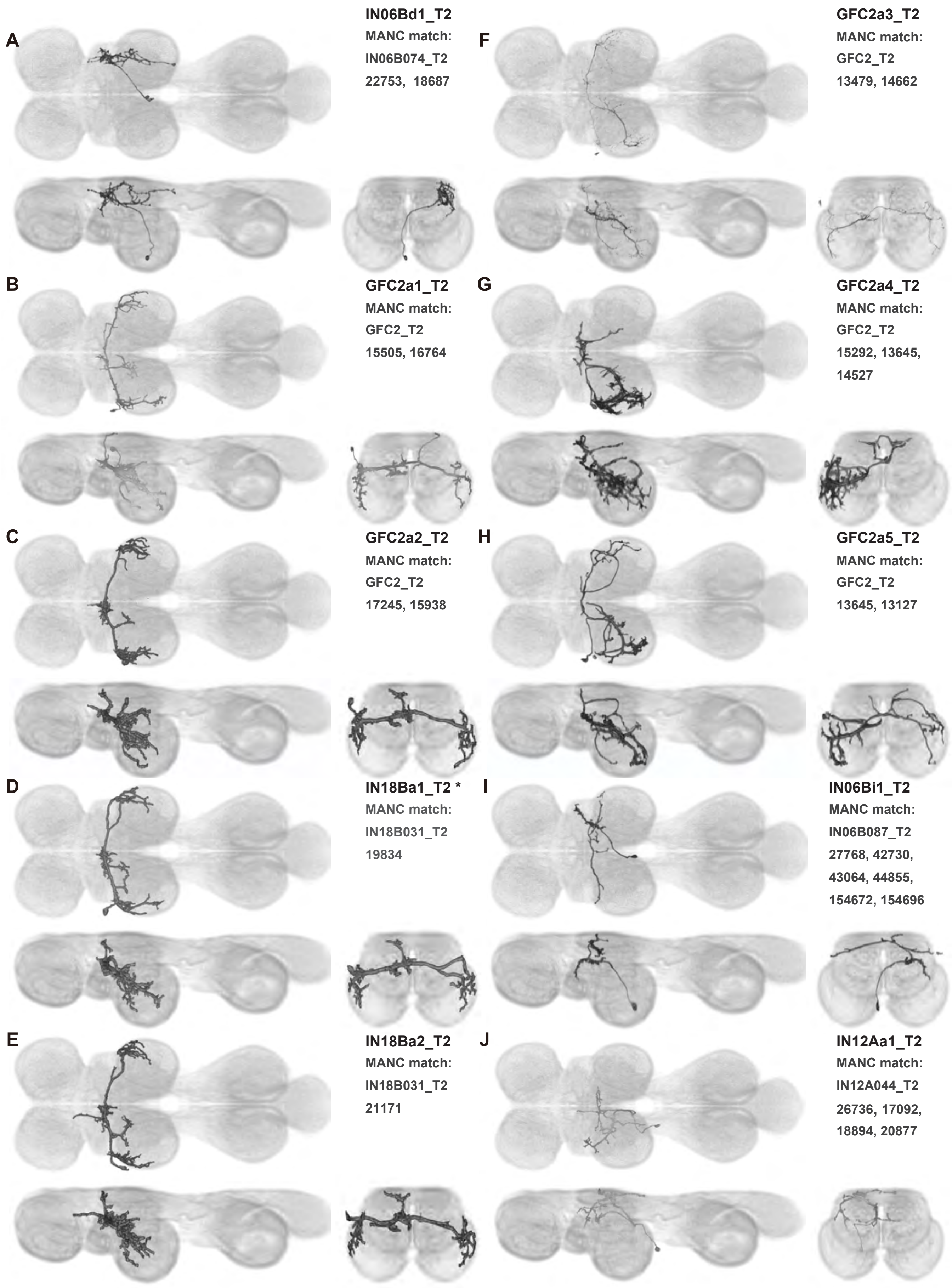
A
IN06Bd1_T2
MANC match:
IN06B074_T2
22753, 18687
F
GFC2a3_T2
MANC match:
GFC2_T2
13479, 14662
B
GFC2a1_T2
MANC match:
GFC2_T2
15505, 16764
G
GFC2a4_T2
MANC match:
GFC2_T2
15292, 13645,
14527
C
GFC2a2_T2
MANC match:
GFC2_T2
17245, 15938
H
GFC2a5_T2
MANC match:
GFC2_T2
13645, 13127
D
IN18Ba1_T2 *
MANC match:
IN18B031_T2
19834
I
IN06Bi1_T2
MANC match:
IN06B087_T2
27768, 42730,
43064, 44855,
154672, 154696
E
IN18Ba2_T2
MANC match:
IN18B031_T2
21171
J
IN12Aa1_T2
MANC match:
IN12A044_T2
26736, 17092,
18894, 20877

A

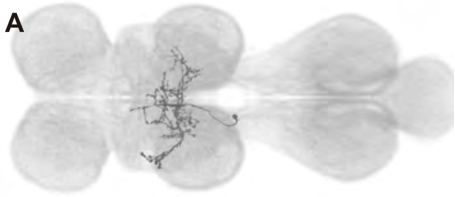

**vMS12a1_T2**
**MANC match:**
**vMS12_T2**
**18452, 12887, 14126, 16889, 19042, 21441, 32714, 101397**

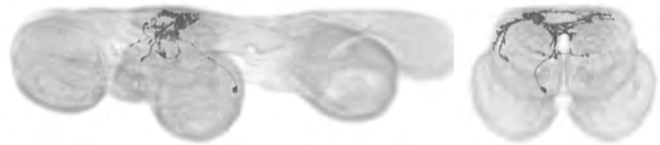

B

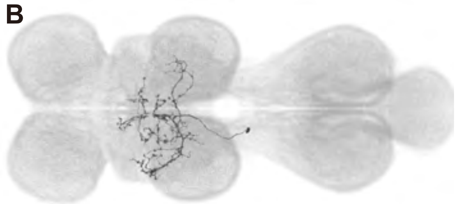

**IN12Aa2_T2**
**MANC match:**
**IN12A044_T2**
**20452, 20890, 21017, 102207**

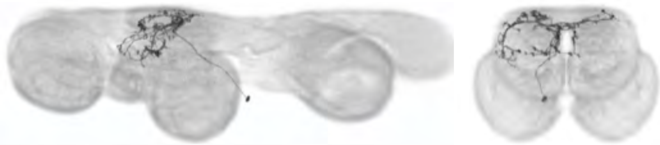

C

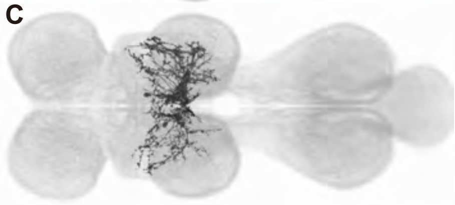

**IN06Ba1_T2**
**MANC match:**
**IN06B043_T2**
**15946, 13408, 13571, 13927, 14152, 14556, 15926, 17422**

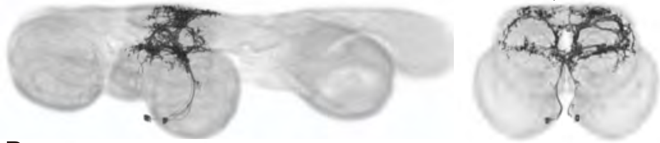

D

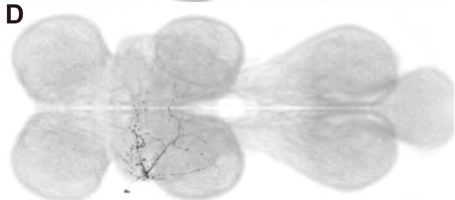

**IN17Ac1_T2**
**MANC match:**
**IN17A071_T2**
**16116, 19687**

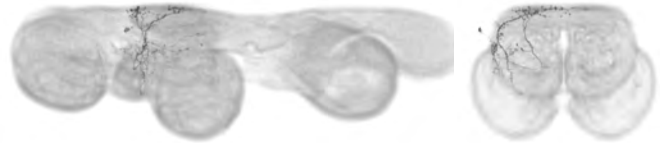

E

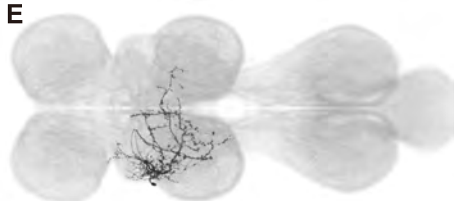

**dMS2a2_T2**
**MANC match:**
**dMS2_T2**
**12742, 14309**

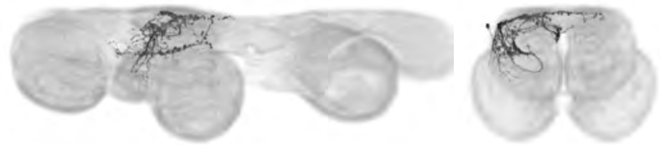

F

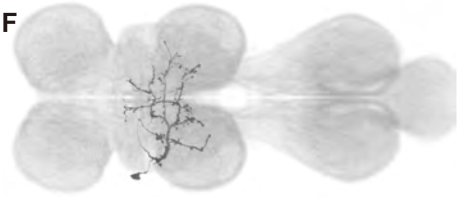

**dMS2a3_T2**
**MANC match:**
**dMS2_T2**
**11302, 10629**

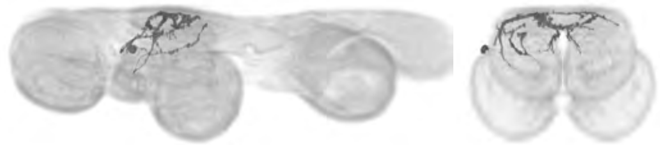

G

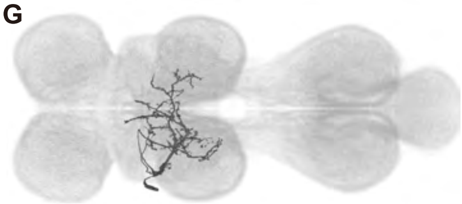

**dMS2a4_T2**
**MANC match:**
**dMS2_T2**
**13229, 14349**

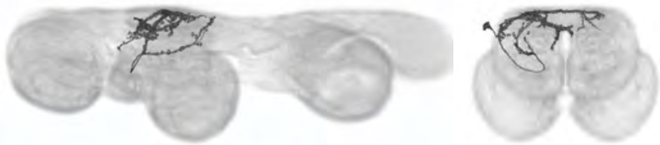

H

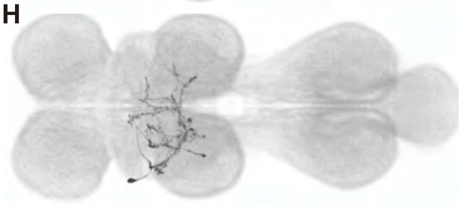

**dMS2a5_T2**
**MANC match:**
**dMS2_T2**
**11032, 11290**

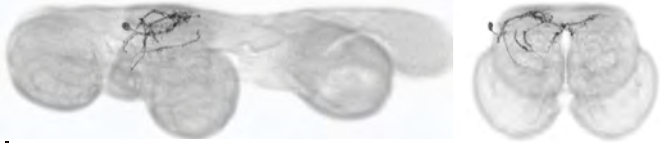

I

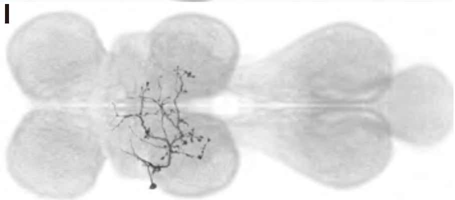

**dMS2a6_T2**
**MANC match:**
**dMS2_T2**
**12520, 14573**

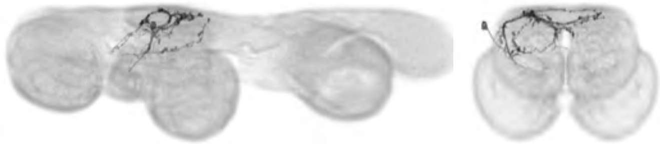

J

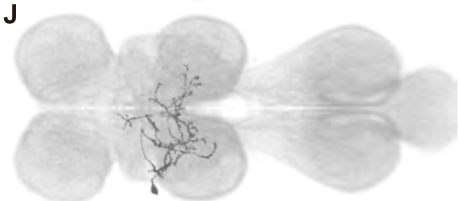

**dMS2a7_T2**
**MANC match:**
**dMS2_T2**
**12207, 16972**

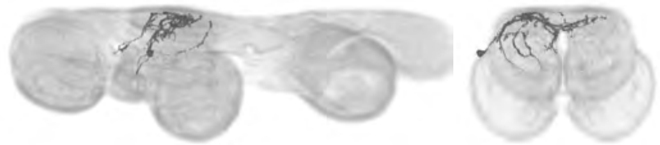

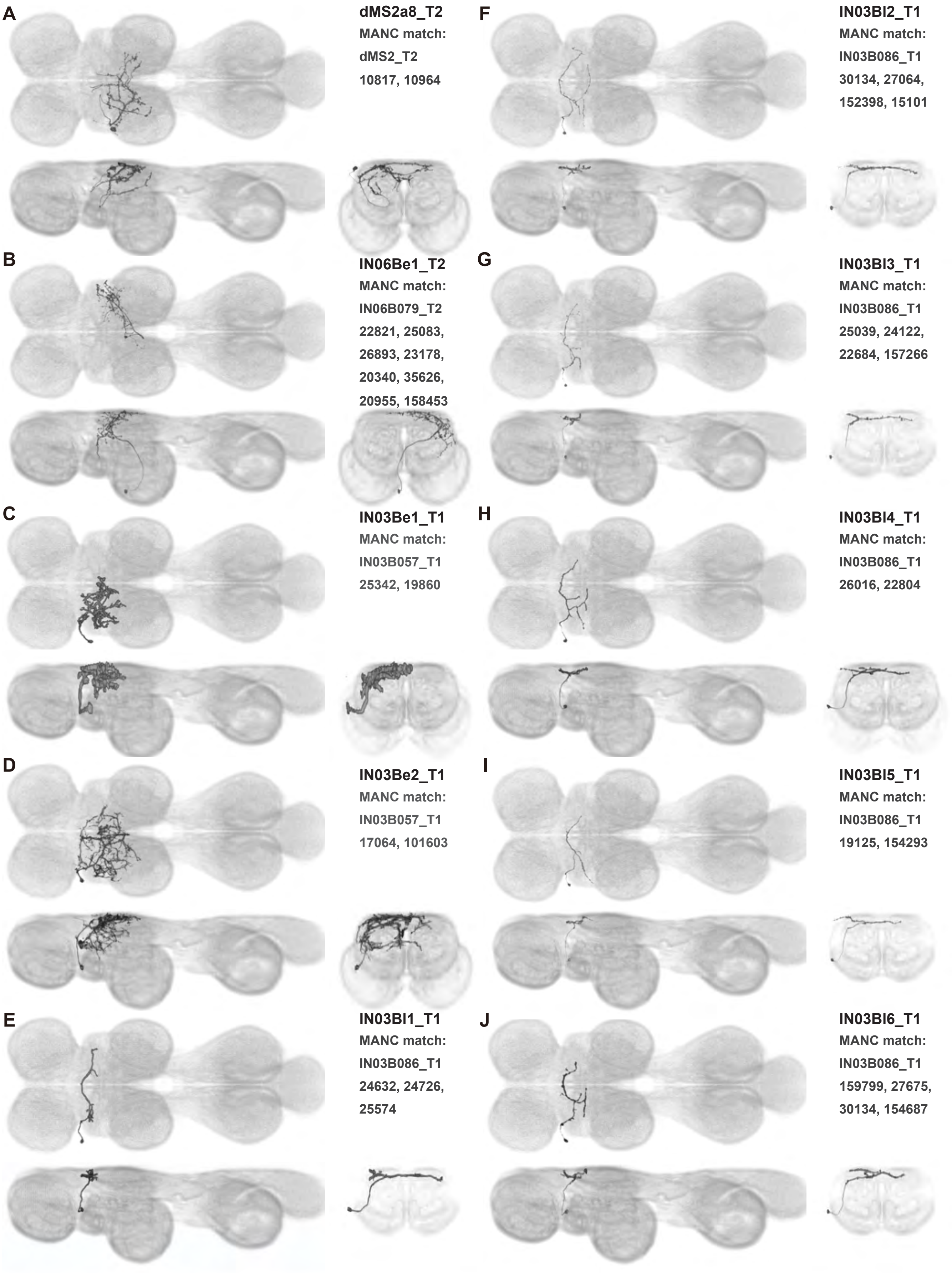
A
dMS2a8_T2
MANC match:
dMS2_T2
10817, 10964
B
IN06Be1_T2
MANC match:
IN06B079_T2
22821, 25083,
26893, 23178,
20340, 35626,
20955, 158453
C
IN03Be1_T1
MANC match:
IN03B057_T1
25342, 19860
D
IN03Be2_T1
MANC match:
IN03B057_T1
17064, 101603
E
IN03Bl1_T1
MANC match:
IN03B086_T1
24632, 24726,
25574
F
IN03Bl2_T1
MANC match:
IN03B086_T1
30134, 27064,
152398, 15101
G
IN03Bl3_T1
MANC match:
IN03B086_T1
25039, 24122,
22684, 157266
H
IN03Bl4_T1
MANC match:
IN03B086_T1
26016, 22804
I
IN03Bl5_T1
MANC match:
IN03B086_T1
19125, 154293
J
IN03Bl6_T1
MANC match:
IN03B086_T1
159799, 27675,
30134, 154687

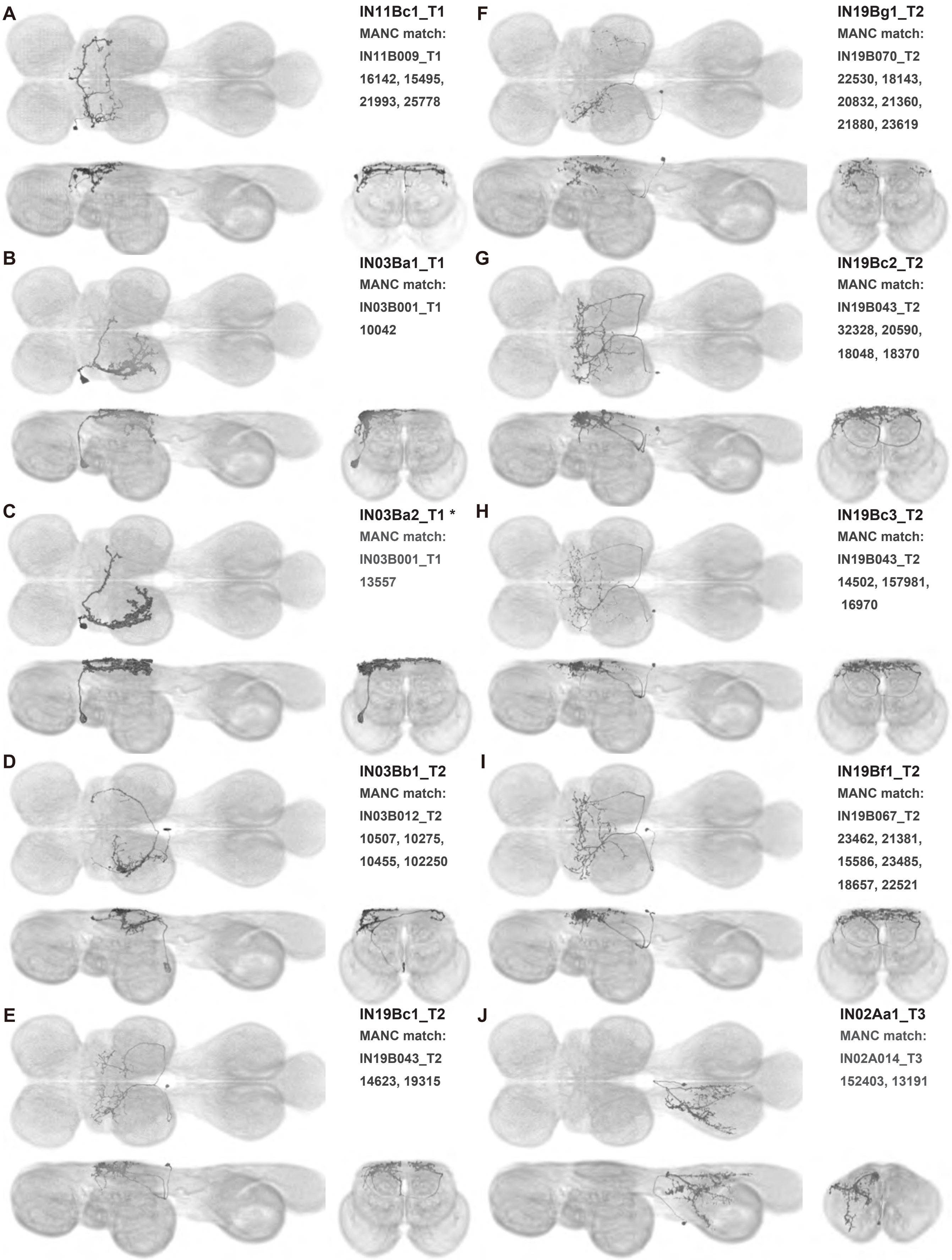
A
IN11Bc1_T1
MANC match:
IN11B009_T1
16142, 15495,
21993, 25778
F
IN19Bg1_T2
MANC match:
IN19B070_T2
22530, 18143,
20832, 21360,
21880, 23619
B
IN03Ba1_T1
MANC match:
IN03B001_T1
10042
G
IN19Bc2_T2
MANC match:
IN19B043_T2
32328, 20590,
18048, 18370
C
IN03Ba2_T1 *
MANC match:
IN03B001_T1
13557
H
IN19Bc3_T2
MANC match:
IN19B043_T2
14502, 157981,
16970
D
IN03Bb1_T2
MANC match:
IN03B012_T2
10507, 10275,
10455, 102250
I
IN19Bf1_T2
MANC match:
IN19B067_T2
23462, 21381,
15586, 23485,
18657, 22521
E
IN19Bc1_T2
MANC match:
IN19B043_T2
14623, 19315
J
IN02Aa1_T3
MANC match:
IN02A014_T3
152403, 13191

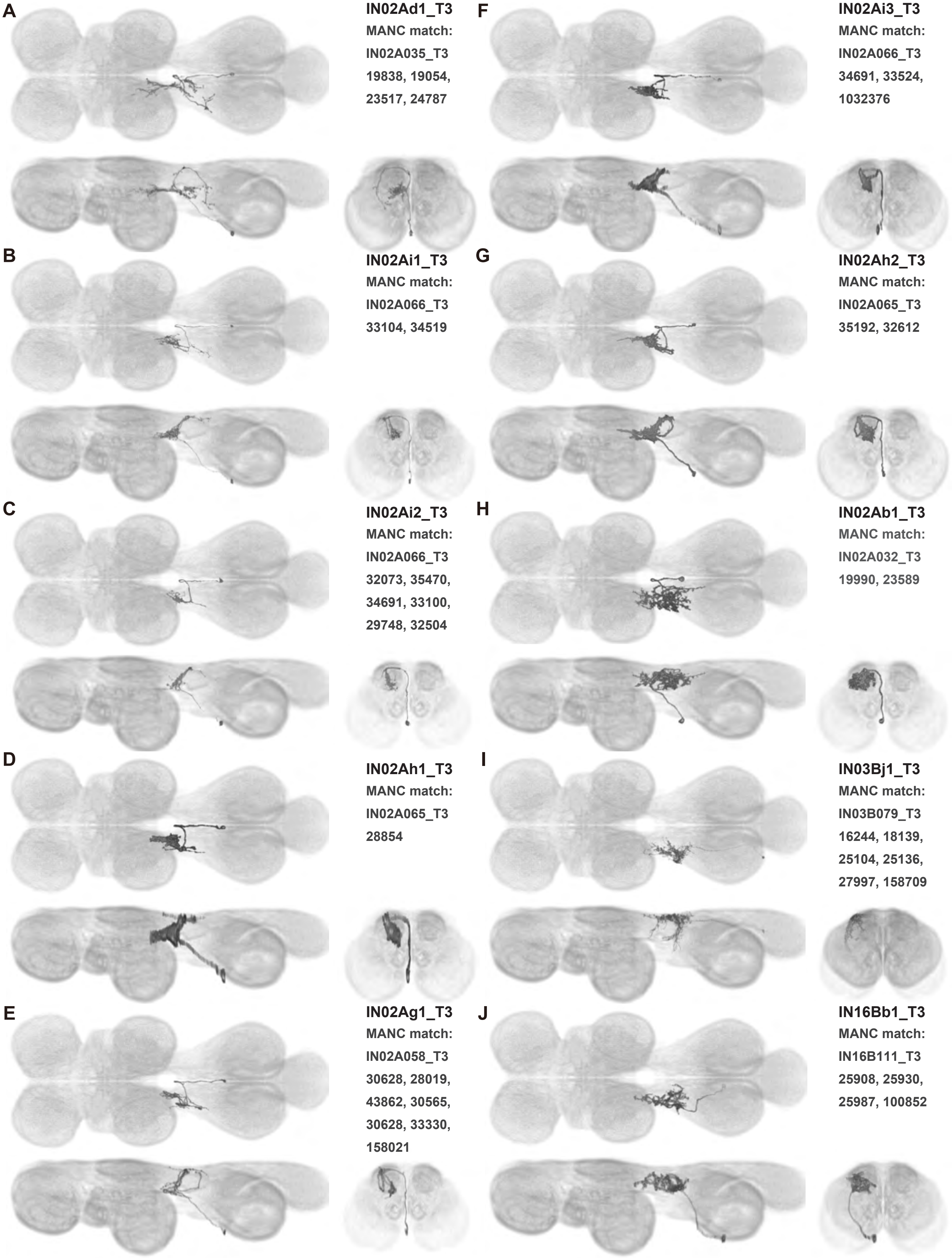
A
IN02Ad1_T3
MANC match:
IN02A035_T3
19838, 19054,
23517, 24787
F
IN02Ai3_T3
MANC match:
IN02A066_T3
34691, 33524,
1032376
B
IN02Ai1_T3
MANC match:
IN02A066_T3
33104, 34519
G
IN02Ah2_T3
MANC match:
IN02A065_T3
35192, 32612
C
IN02Ai2_T3
MANC match:
IN02A066_T3
32073, 35470,
34691, 33100,
29748, 32504
H
IN02Ab1_T3
MANC match:
IN02A032_T3
19990, 23589
D
IN02Ah1_T3
MANC match:
IN02A065_T3
28854
I
IN03Bj1_T3
MANC match:
IN03B079_T3
16244, 18139,
25104, 25136,
27997, 158709
E
IN02Ag1_T3
MANC match:
IN02A058_T3
30628, 28019,
43862, 30565,
30628, 33330,
158021
J
IN16Bb1_T3
MANC match:
IN16B111_T3
25908, 25930,
25987, 100852

**A**

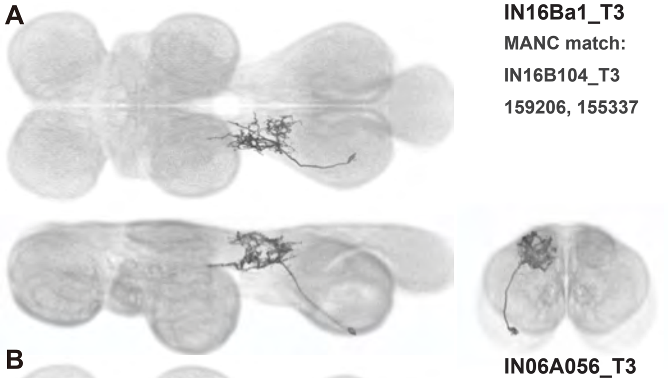

**IN16Ba1_T3**
**MANC match:**
**IN16B104_T3**
**159206, 155337**

**B**

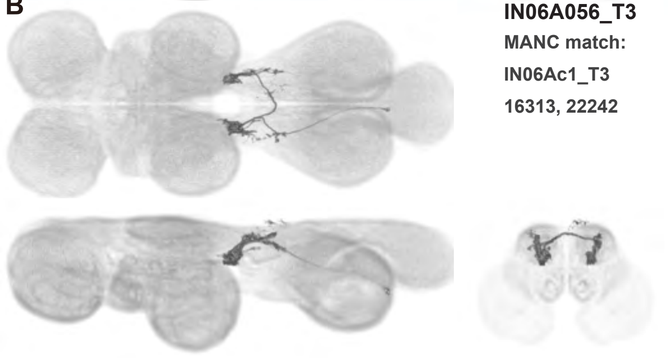

**IN06A056_T3**
**MANC match:**
**IN06Ac1_T3**
**16313, 22242**

Supplemental Figure 16.8

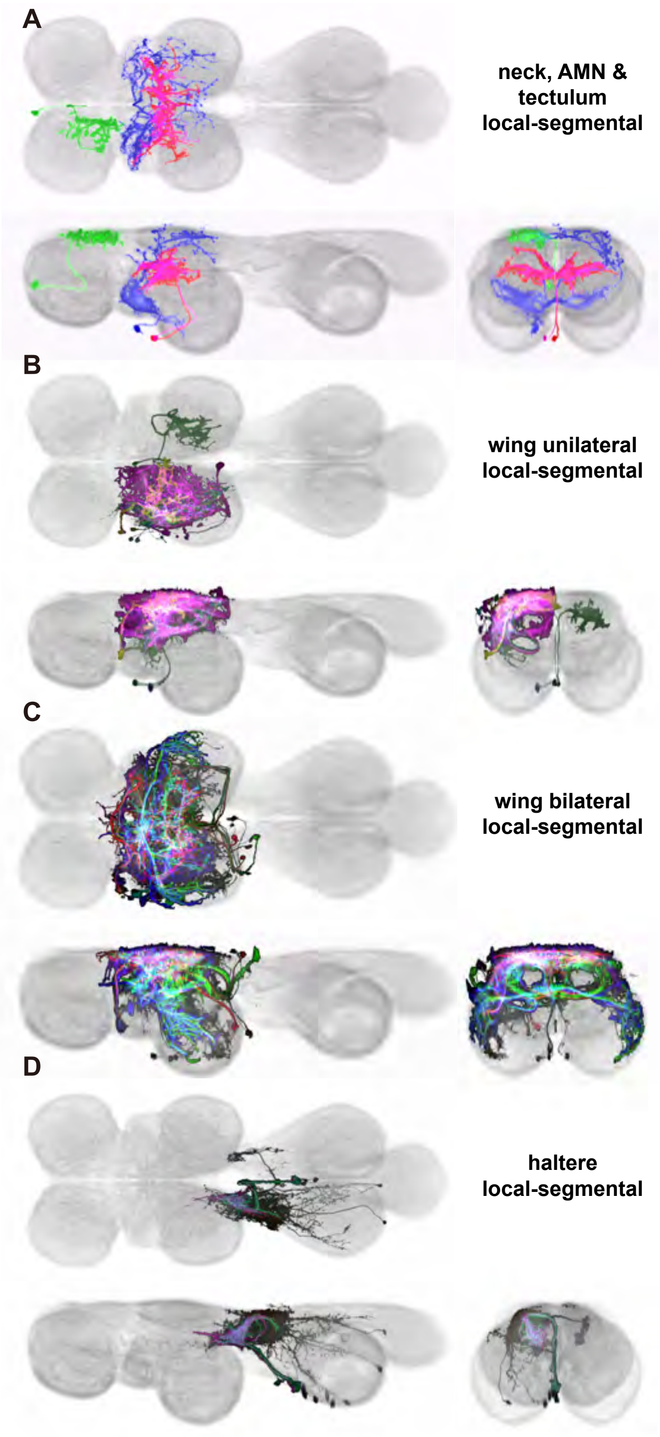

Supplemental Figure 17.1

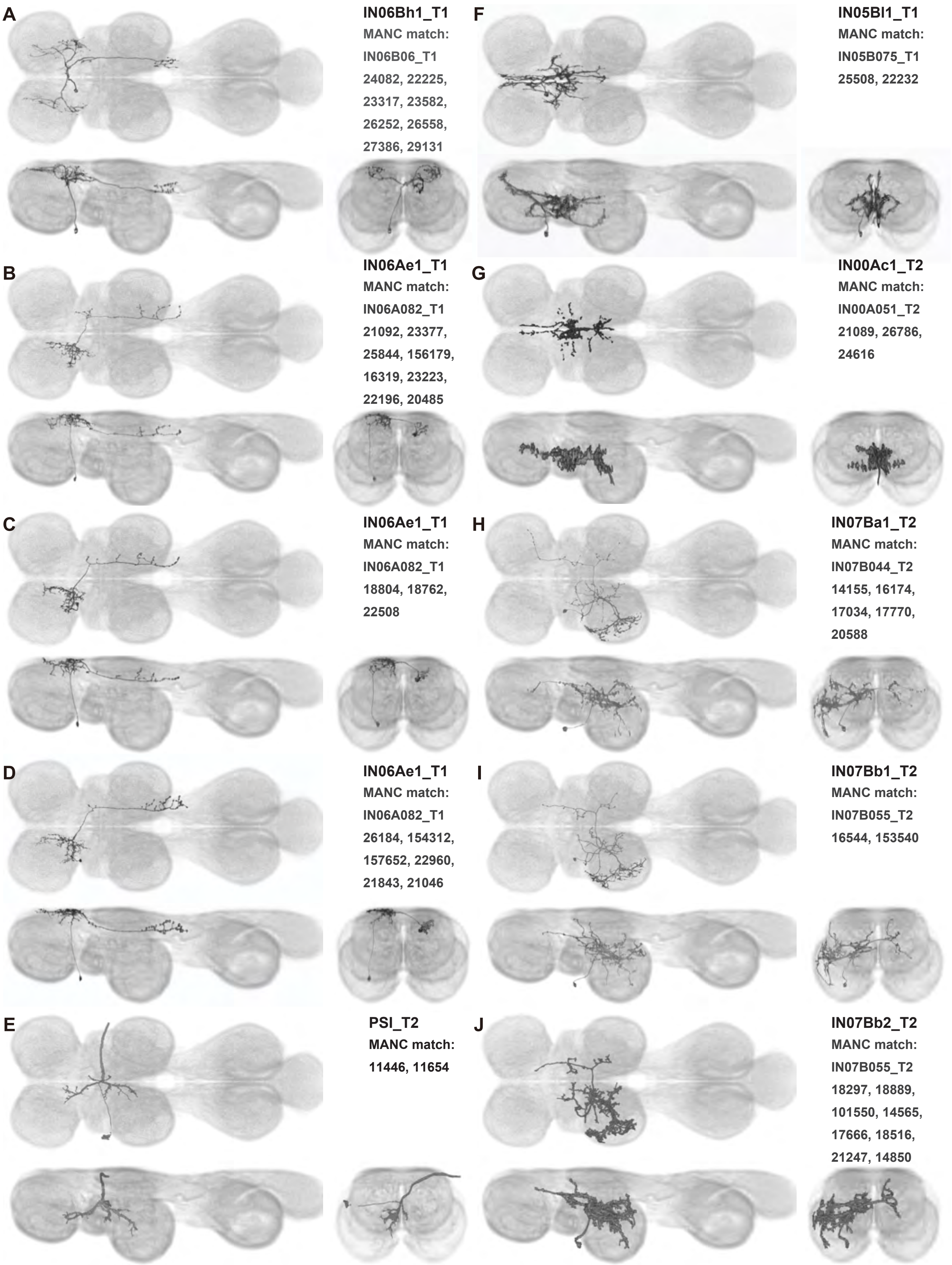

**A**

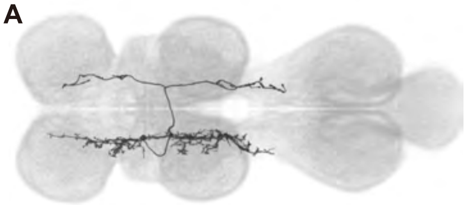

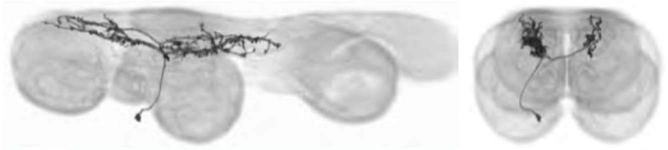

**IN07Bc1_T2**
**MANC match:**
**IN07B063_T2**
**21129, 19664,**
**23839, 24141**

**B**

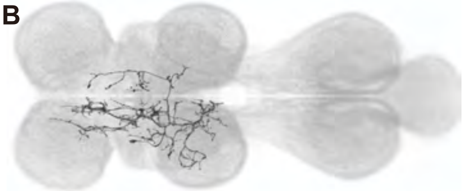

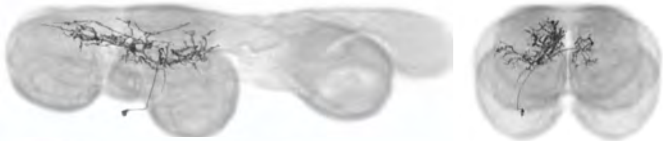

**IN07Bd2_T2**
**MANC match:**
**IN07B066_T2**
**23779, 21437**

**C**

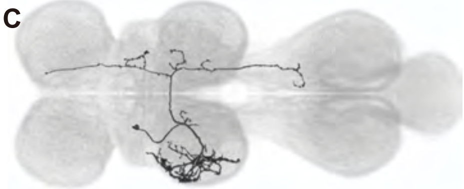

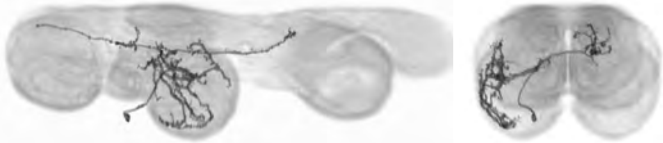

**IN07Be1_T2**
**MANC match:**
**IN07B073_T2**
**23748, 102787,**
**23176, 21760,**
**22311**

**D**

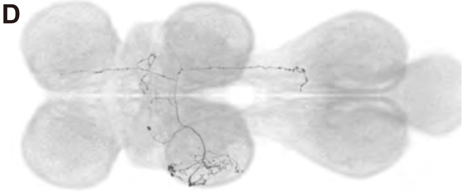

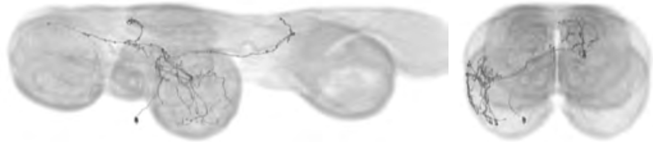

**IN07Be2_T2**
**MANC match:**
**IN07B073_T2**
**24211, 23354**

**E**

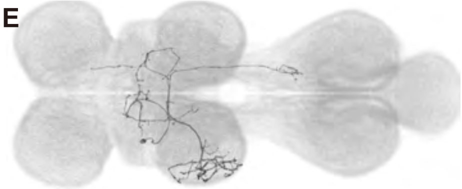

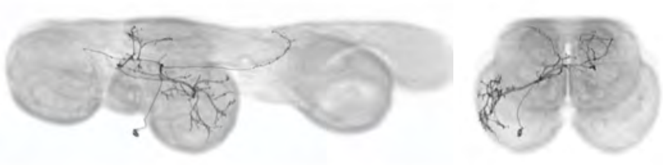

**IN07Be3_T2**
**MANC match:**
**IN07B073_T2**
**21813, 21493,**
**163077, 15584,**
**21153, 30747**

**F**

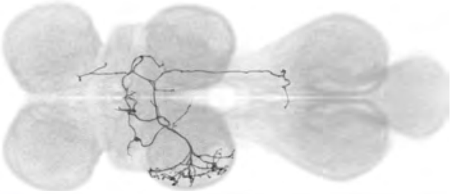

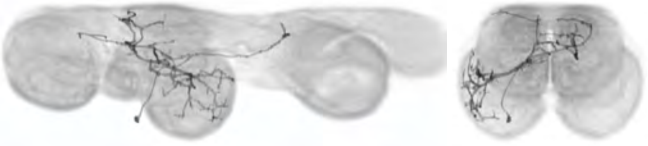

**IN07Be4_T2**
**MANC match:**
**IN07B073_T2**
**21493, 21809,**
**22638, 22266**

**G**

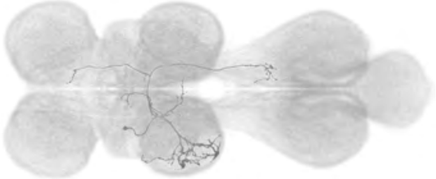

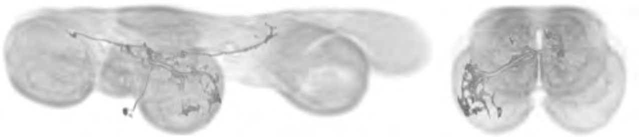

**IN07Be5_T2**
**MANC match:**
**IN07B073_T2**
**22311, 20004,**
**24145, 21839,**
**20223**

**H**

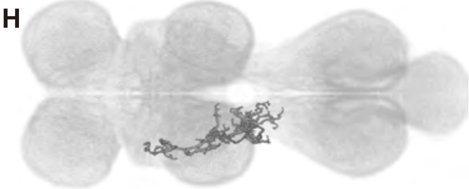

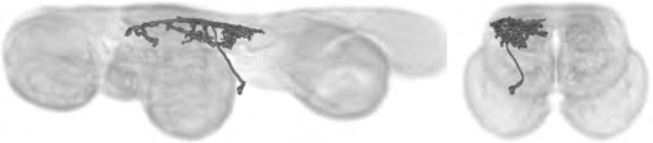

**IN03Bg1_T2**
**MANC match:**
**IN03B068_T2**
**158944, 156198**

**I**

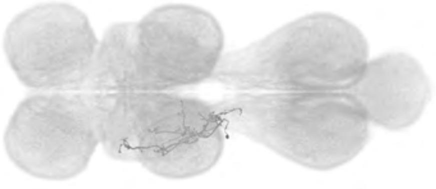

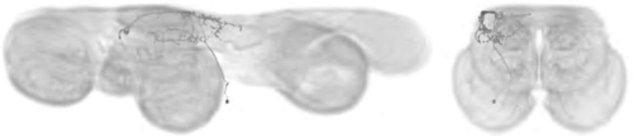

**IN03Bh1_T2**
**MANC match:**
**IN03B070_T2**
**23027, 24249,**
**23612**

**J**

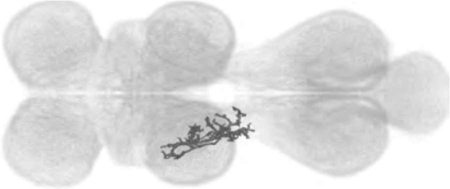

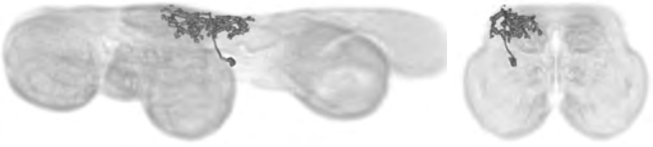

**IN03Bf1_T2**
**MANC match:**
**IN03B060_T2**
**163396, 18385,**
**21623, 159210,**
**152816, 17811,**
**22328, 19033**

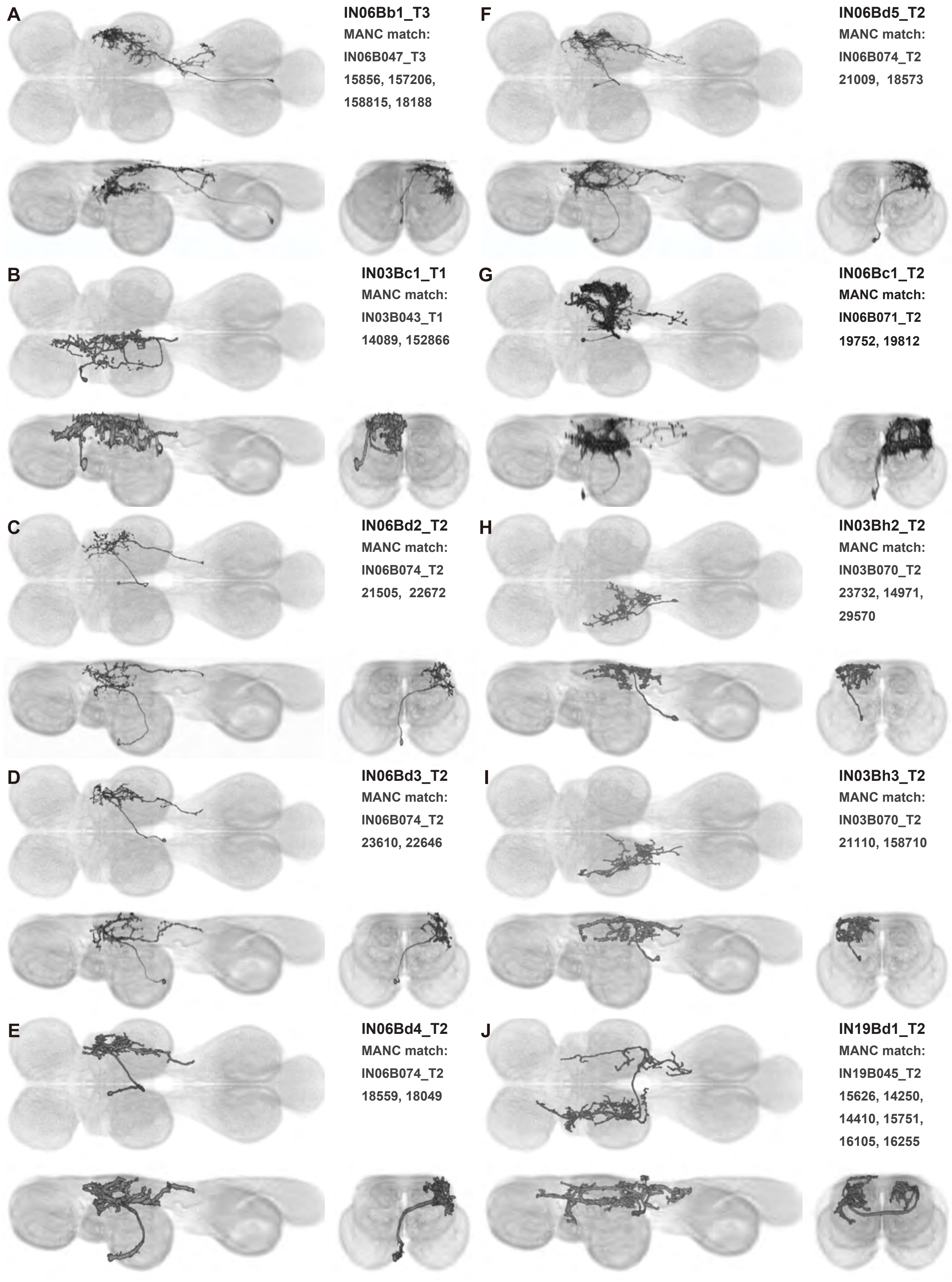
A
IN06Bb1_T3
MANC match:
IN06B047_T3
15856, 157206,
158815, 18188
F
IN06Bd5_T2
MANC match:
IN06B074_T2
21009, 18573
B
IN03Bc1_T1
MANC match:
IN03B043_T1
14089, 152866
G
IN06Bc1_T2
MANC match:
IN06B071_T2
19752, 19812
C
IN06Bd2_T2
MANC match:
IN06B074_T2
21505, 22672
H
IN03Bh2_T2
MANC match:
IN03B070_T2
23732, 14971,
29570
D
IN06Bd3_T2
MANC match:
IN06B074_T2
23610, 22646
I
IN03Bh3_T2
MANC match:
IN03B070_T2
21110, 158710
E
IN06Bd4_T2
MANC match:
IN06B074_T2
18559, 18049
J
IN19Bd1_T2
MANC match:
IN19B045_T2
15626, 14250,
14410, 15751,
16105, 16255

Supplemental Figure 17.4

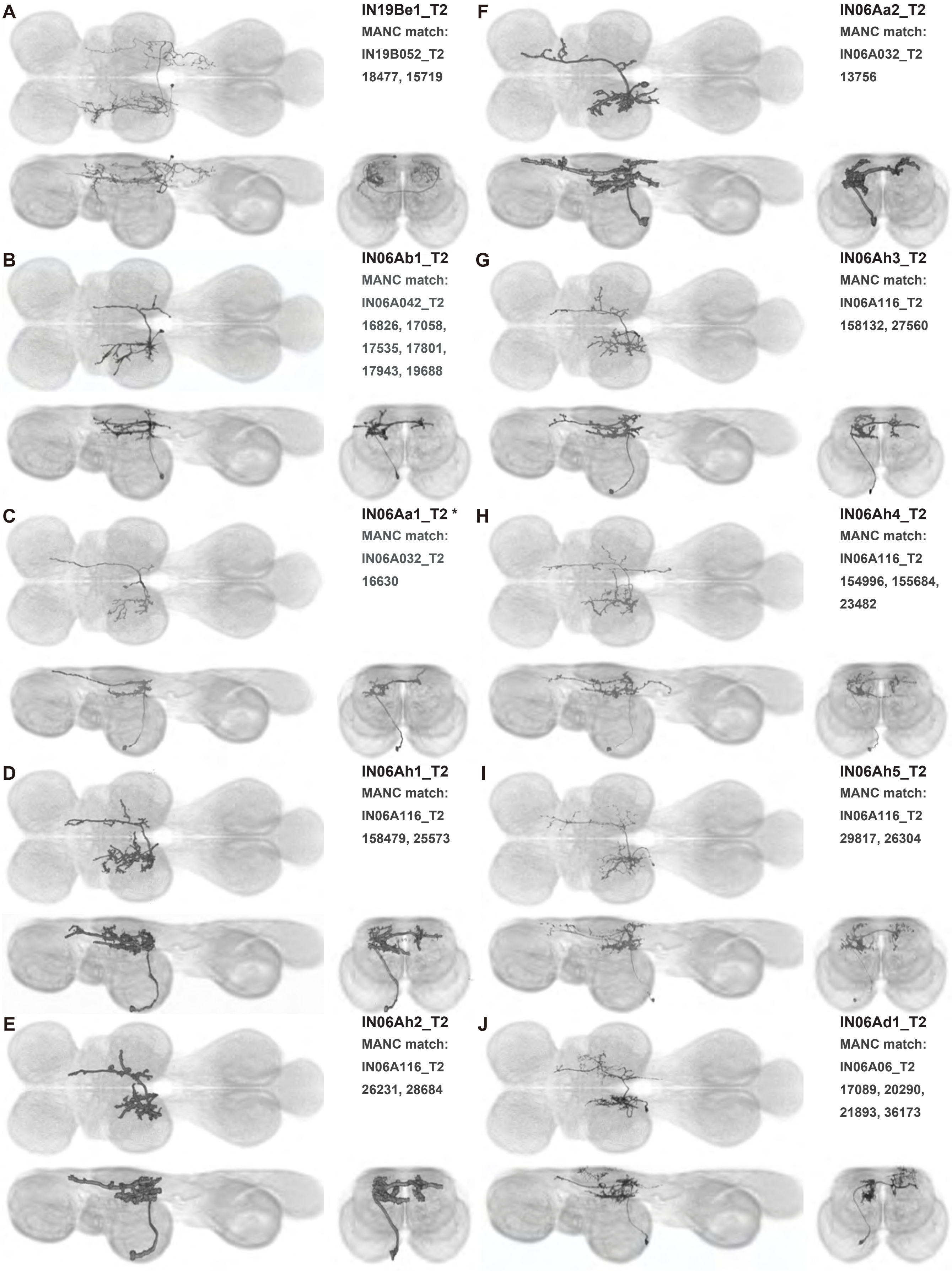

A

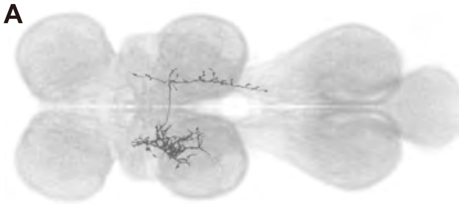

**IN07Bg1_T2**
**MANC match:**
**IN07B084_T2**
**26518, 24270,**
**27702, 159067**

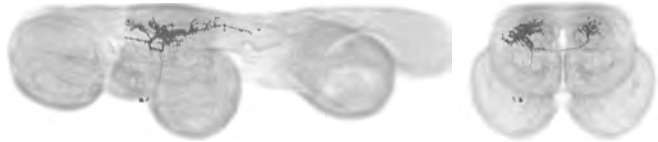

B

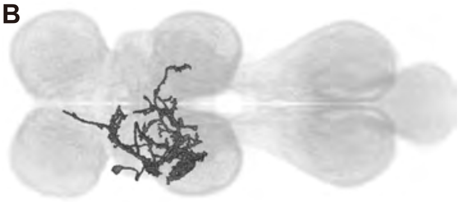

**dMS2a9_T2**
**MANC match:**
**dMS2_T2**
**12056, 11642**

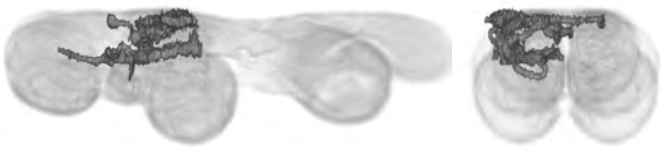

C

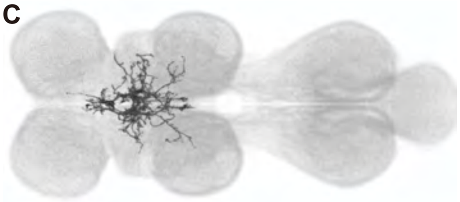

**IN00Ab1_T2**
**MANC match:**
**IN00A043_T1**
**25798, 13859,**
**19605**

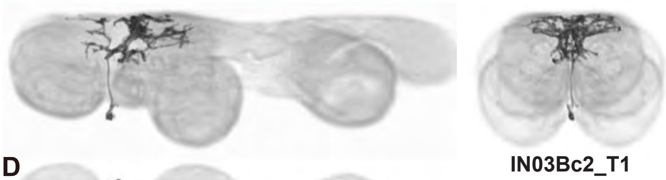

D

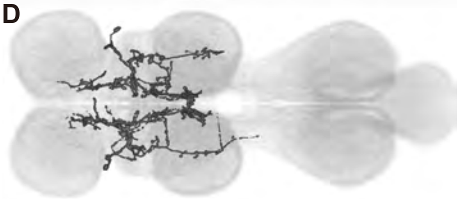

**IN03Bc2_T1**
**MANC match:**
**IN03B043_T1**
**17024, 15233**

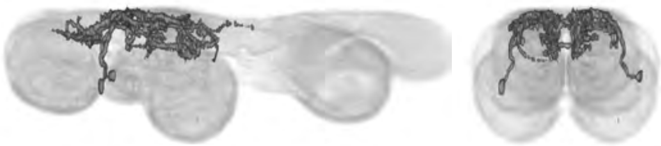

E

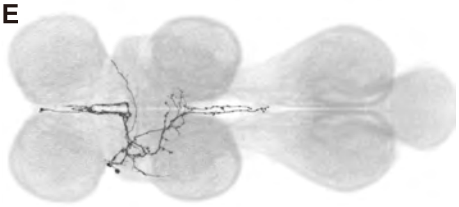

**IN03Bd1_T1**
**MANC match:**
**IN03B054_T1**
**160309, 16231,**
**17758, 20515,**
**21123, 21187**

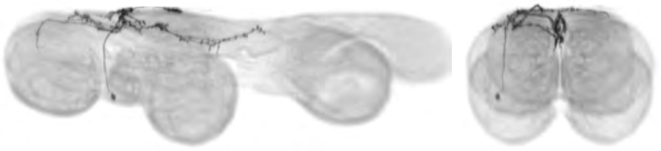

F

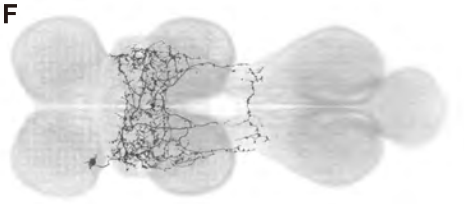

**IN19Bb1_T1**
**MANC match:**
**IN19B034_T1**
**13743, 14036**

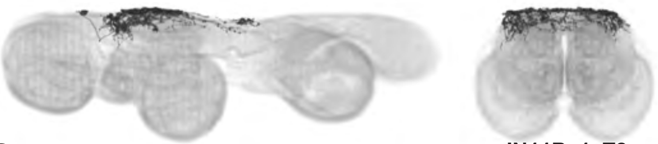

G

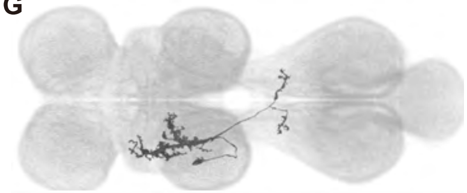

**IN11Ba1_T2**
**MANC match:**
**IN11B001_T2**
**20190, 10326,**
**10729, 11142,**
**11313, 18653**

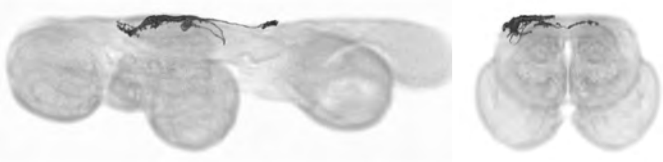

H

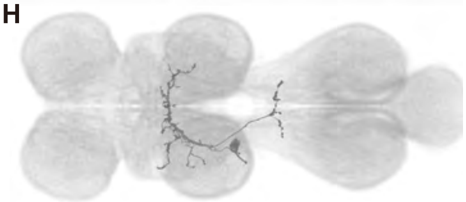

**IN11Bb1_T2**
**MANC match:**
**IN11B003_T2**
**16248, 24971,**
**17120, 13201**

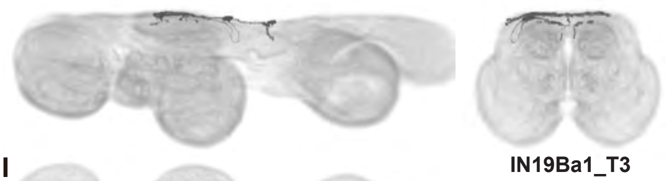

I

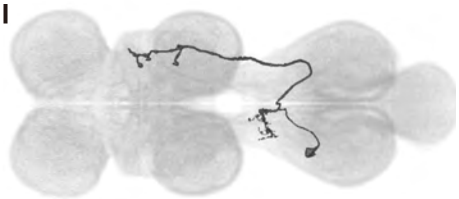

**IN19Ba1_T3**
**MANC match:**
**IN19B013_T3**
**13357, 10177,**
**11181, 11418**

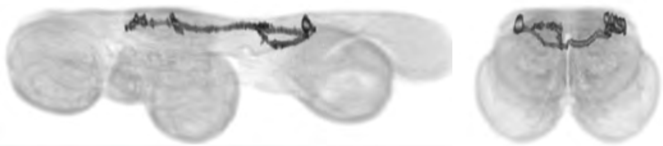

J

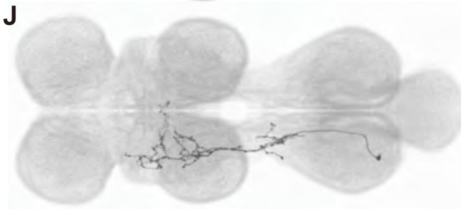

**IN03Bk1_A1**
**MANC match:**
**IN03B082_A1**
**21755, 25441**

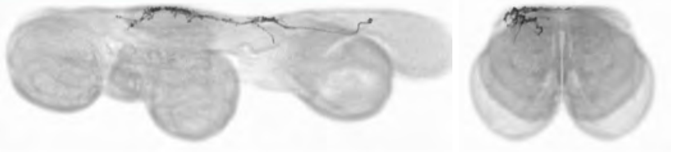

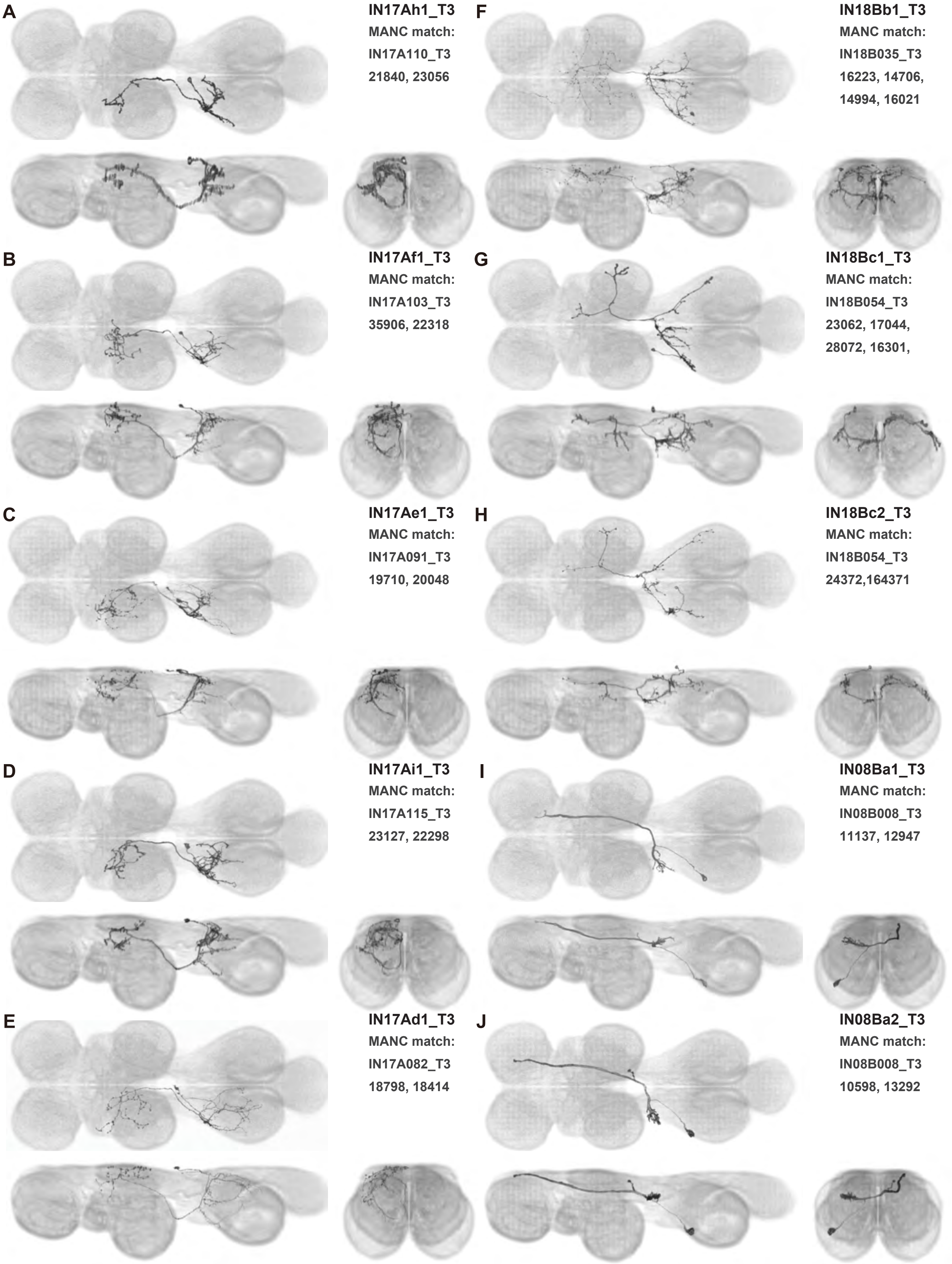
A
IN17Ah1_T3
MANC match:
IN17A110_T3
21840, 23056
F
IN18Bb1_T3
MANC match:
IN18B035_T3
16223, 14706,
14994, 16021
B
IN17Af1_T3
MANC match:
IN17A103_T3
35906, 22318
G
IN18Bc1_T3
MANC match:
IN18B054_T3
23062, 17044,
28072, 16301,
C
IN17Ae1_T3
MANC match:
IN17A091_T3
19710, 20048
H
IN18Bc2_T3
MANC match:
IN18B054_T3
24372,164371
D
IN17Ai1_T3
MANC match:
IN17A115_T3
23127, 22298
I
IN08Ba1_T3
MANC match:
IN08B008_T3
11137, 12947
E
IN17Ad1_T3
MANC match:
IN17A082_T3
18798, 18414
J
IN08Ba2_T3
MANC match:
IN08B008_T3
10598, 13292

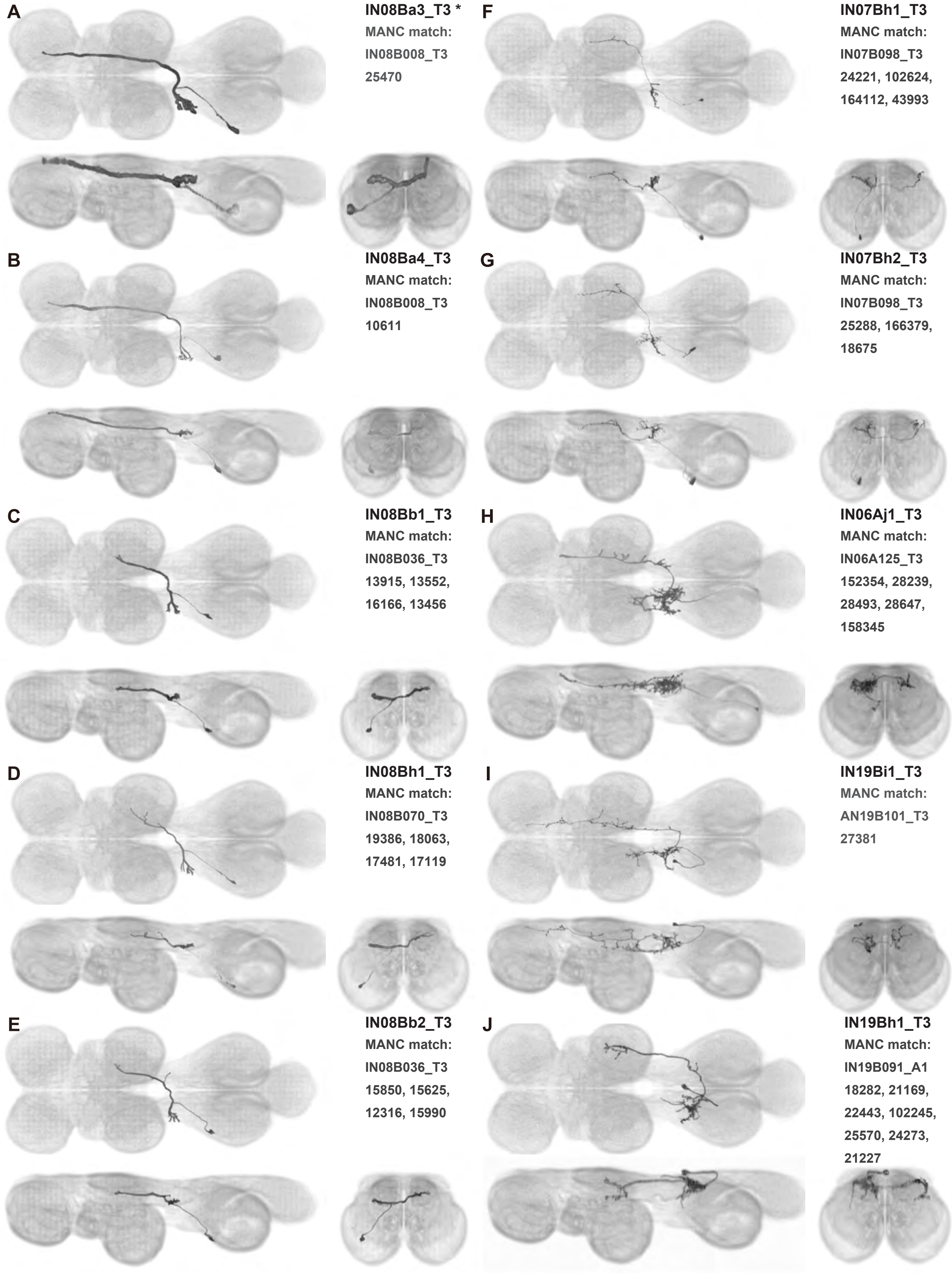

A IN08Ba3_T3 *
MANC match:
IN08B008_T3
25470

B IN08Ba4_T3
MANC match:
IN08B008_T3
10611

C IN08Bb1_T3
MANC match:
IN08B036_T3
13915, 13552,
16166, 13456

D IN08Bh1_T3
MANC match:
IN08B070_T3
19386, 18063,
17481, 17119

E IN08Bb2_T3
MANC match:
IN08B036_T3
15850, 15625,
12316, 15990

F IN07Bh1_T3
MANC match:
IN07B098_T3
24221, 102624,
164112, 43993

G IN07Bh2_T3
MANC match:
IN07B098_T3
25288, 166379,
18675

H IN06Aj1_T3
MANC match:
IN06A125_T3
152354, 28239,
28493, 28647,
158345

I IN19Bi1_T3
MANC match:
AN19B101_T3
27381

J IN19Bh1_T3
MANC match:
IN19B091_A1
18282, 21169,
22443, 102245,
25570, 24273,
21227

A

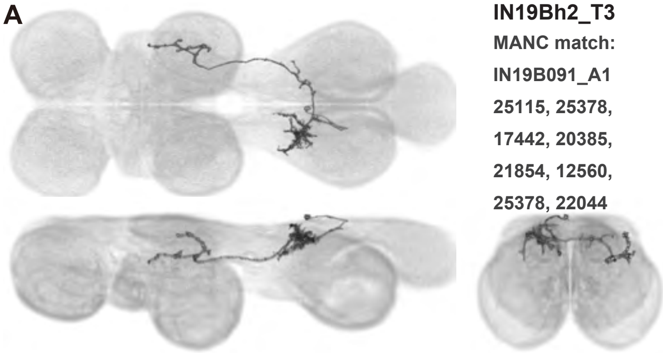

**IN19Bh2_T3**
MANC match:
IN19B091_A1
25115, 25378,
17442, 20385,
21854, 12560,
25378, 22044

B

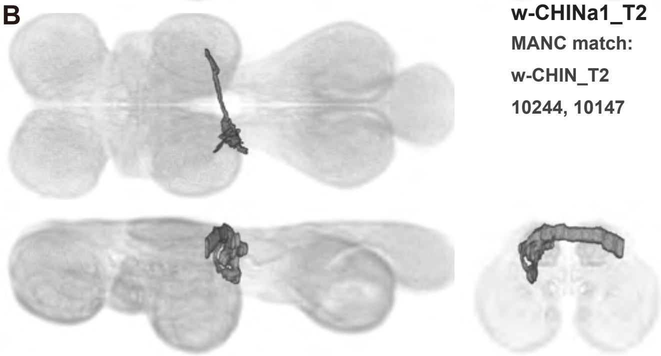

**w-CHINa1_T2**
MANC match:
w-CHIN_T2
10244, 10147

C

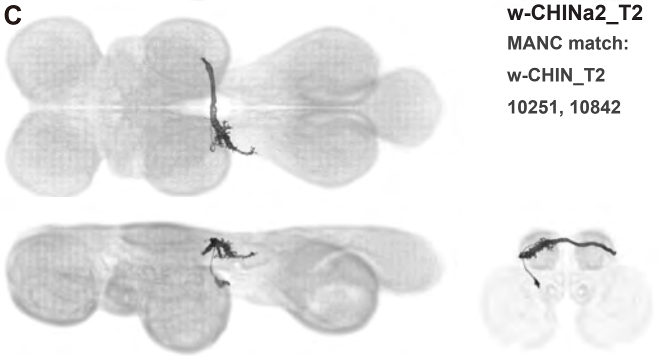

**w-CHINa2_T2**
MANC match:
w-CHIN_T2
10251, 10842

D

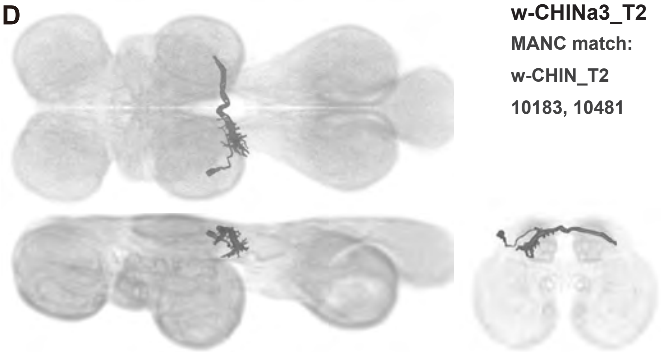

**w-CHINa3_T2**
MANC match:
w-CHIN_T2
10183, 10481

E

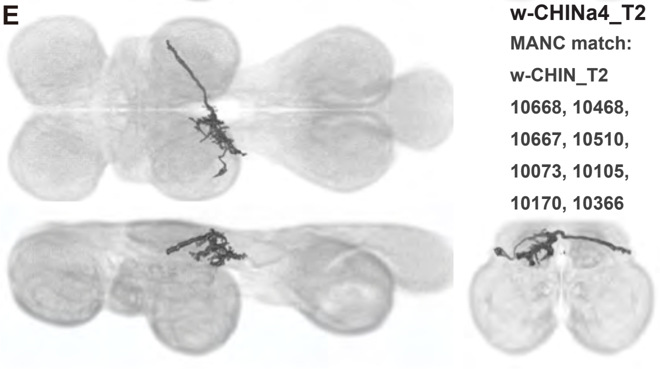

**w-CHINa4_T2**
MANC match:
w-CHIN_T2
10668, 10468,
10667, 10510,
10073, 10105,
10170, 10366

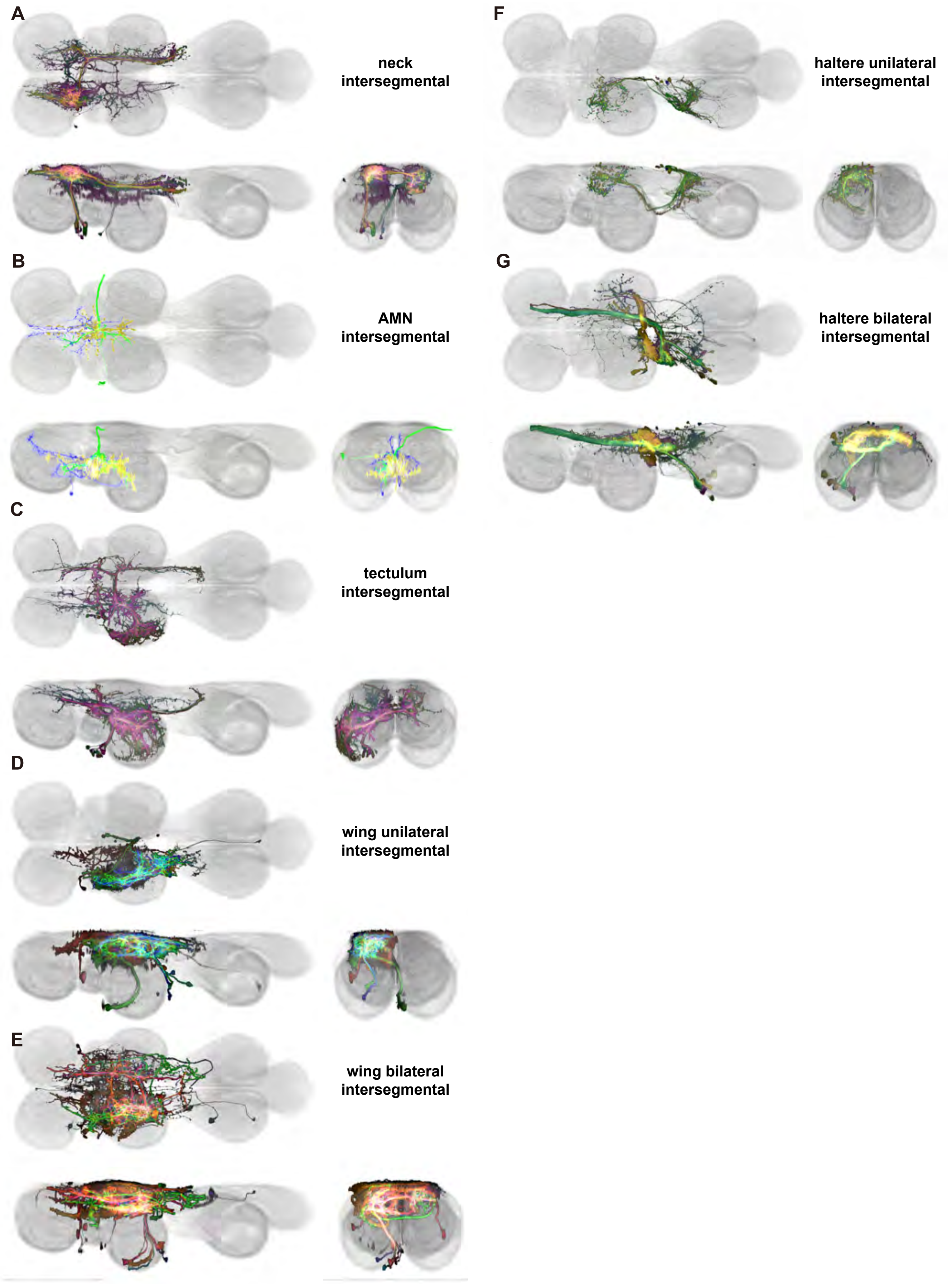
A
neck intersegmental
B
AMN intersegmental
C
tectulum intersegmental
D
wing unilateral intersegmental
E
wing bilateral intersegmental
F
haltere unilateral intersegmental
G
haltere bilateral intersegmental

Supplemental Figure 18.1

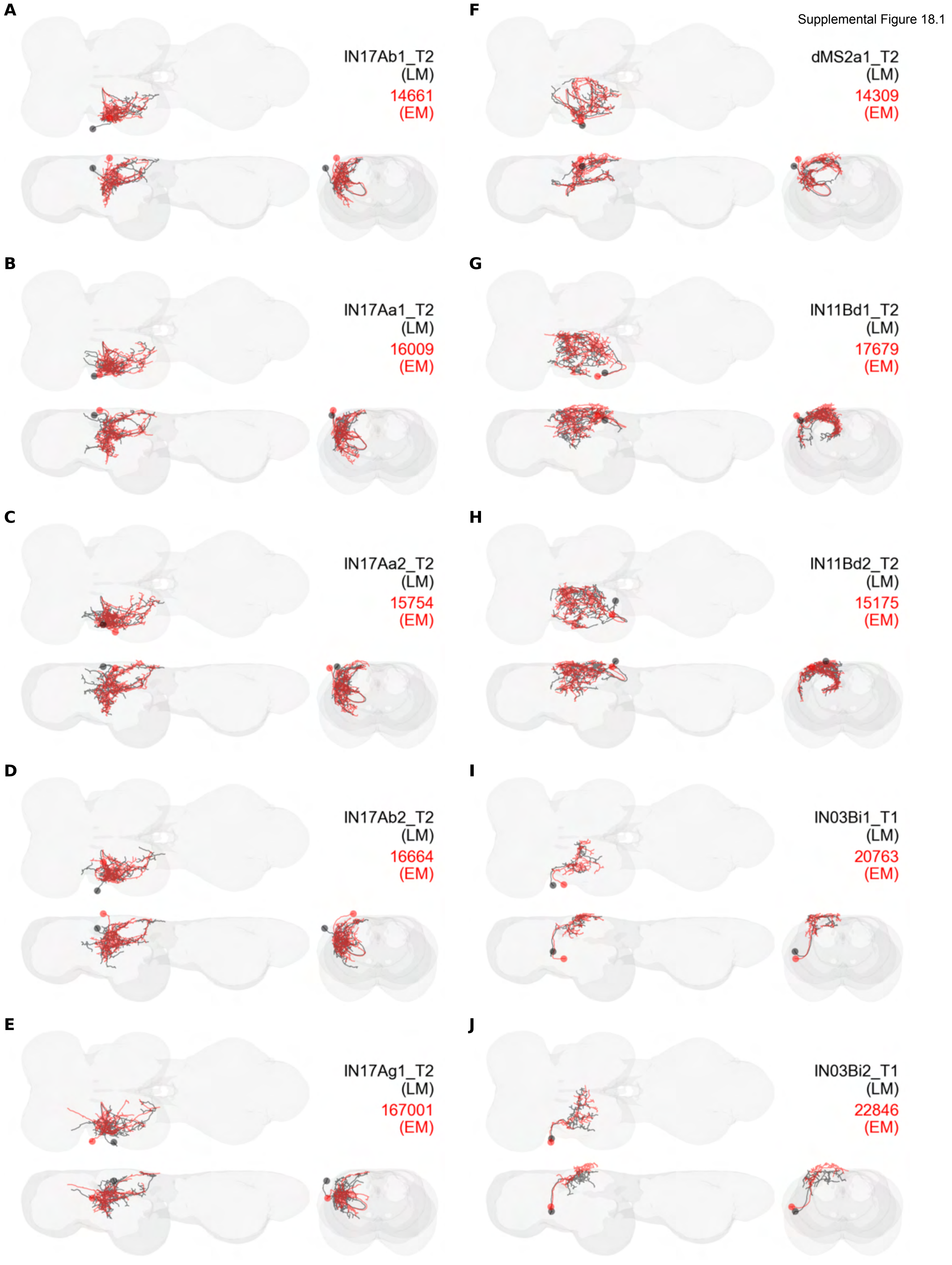

Supplemental Figure 18.2

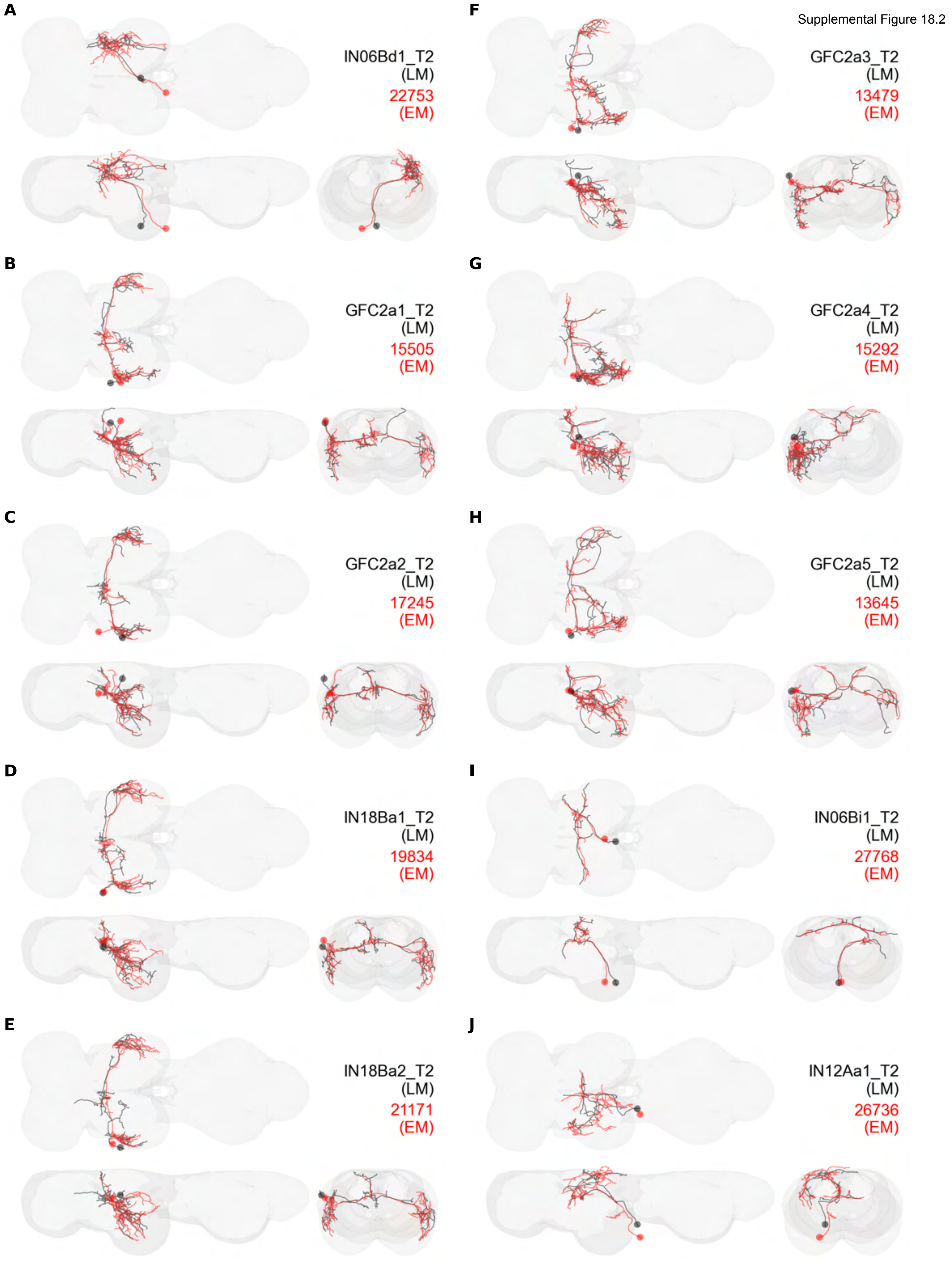

Supplemental Figure 18.3

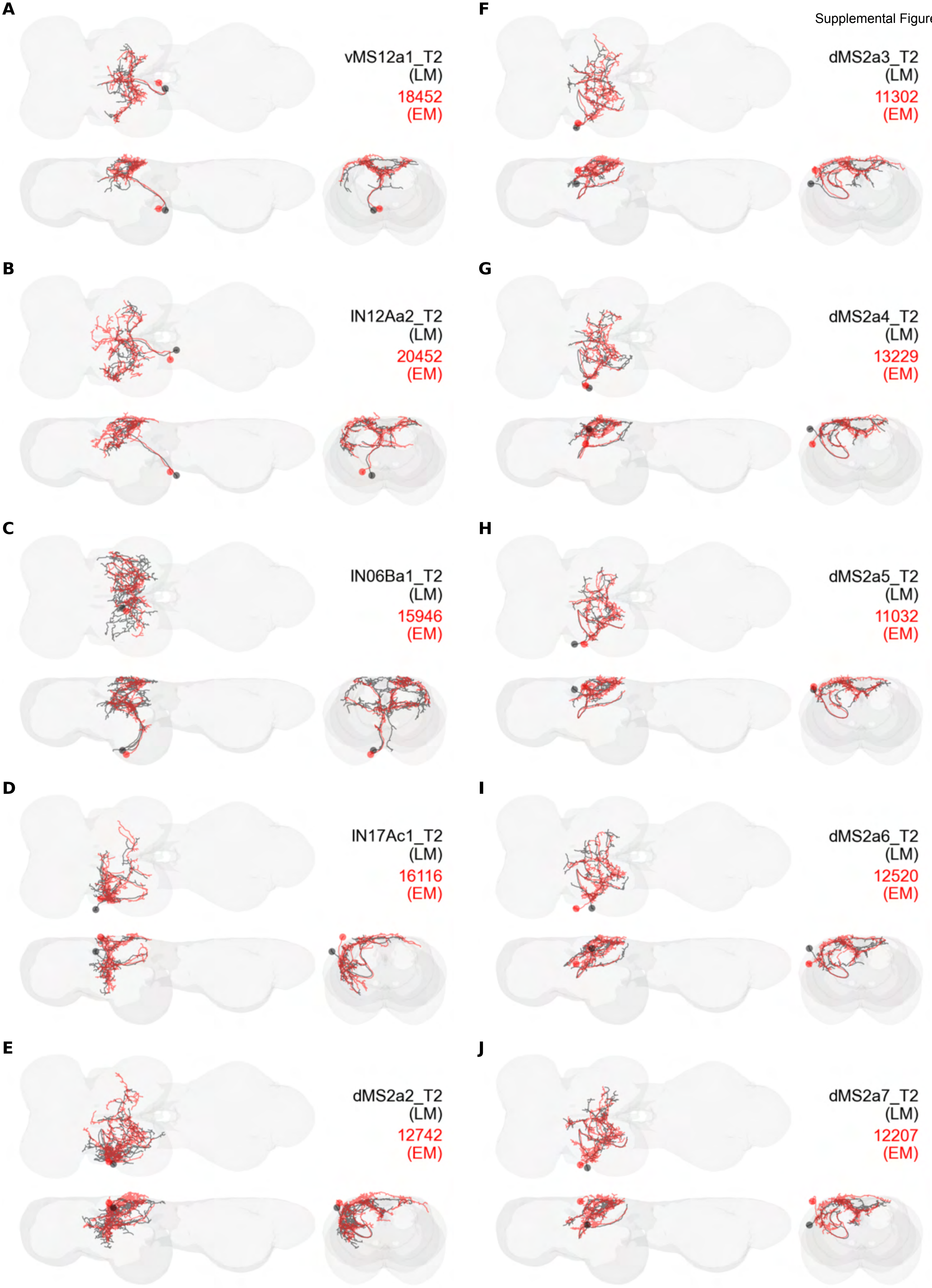

Supplemental Figure 18.4

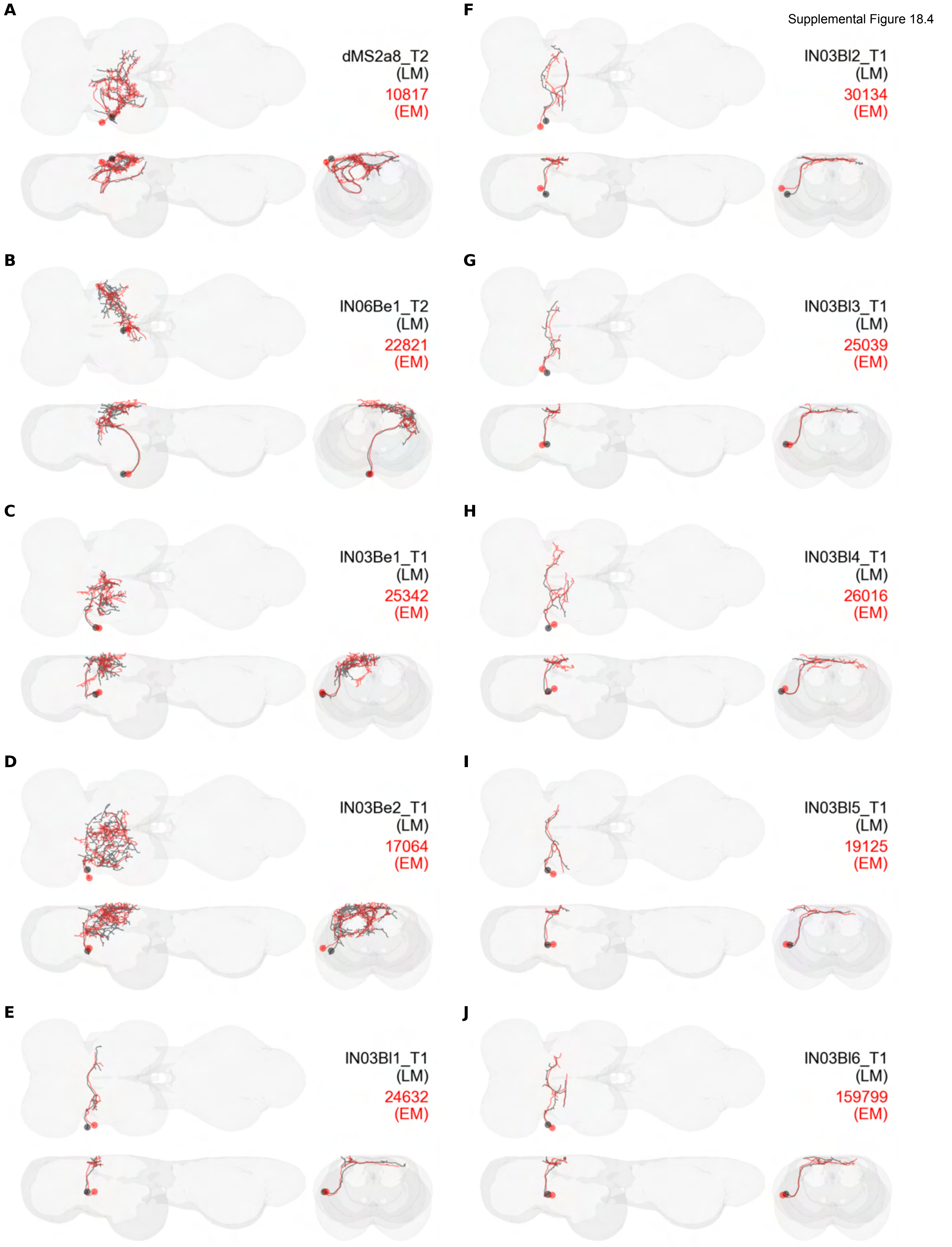

Supplemental Figure 18.5

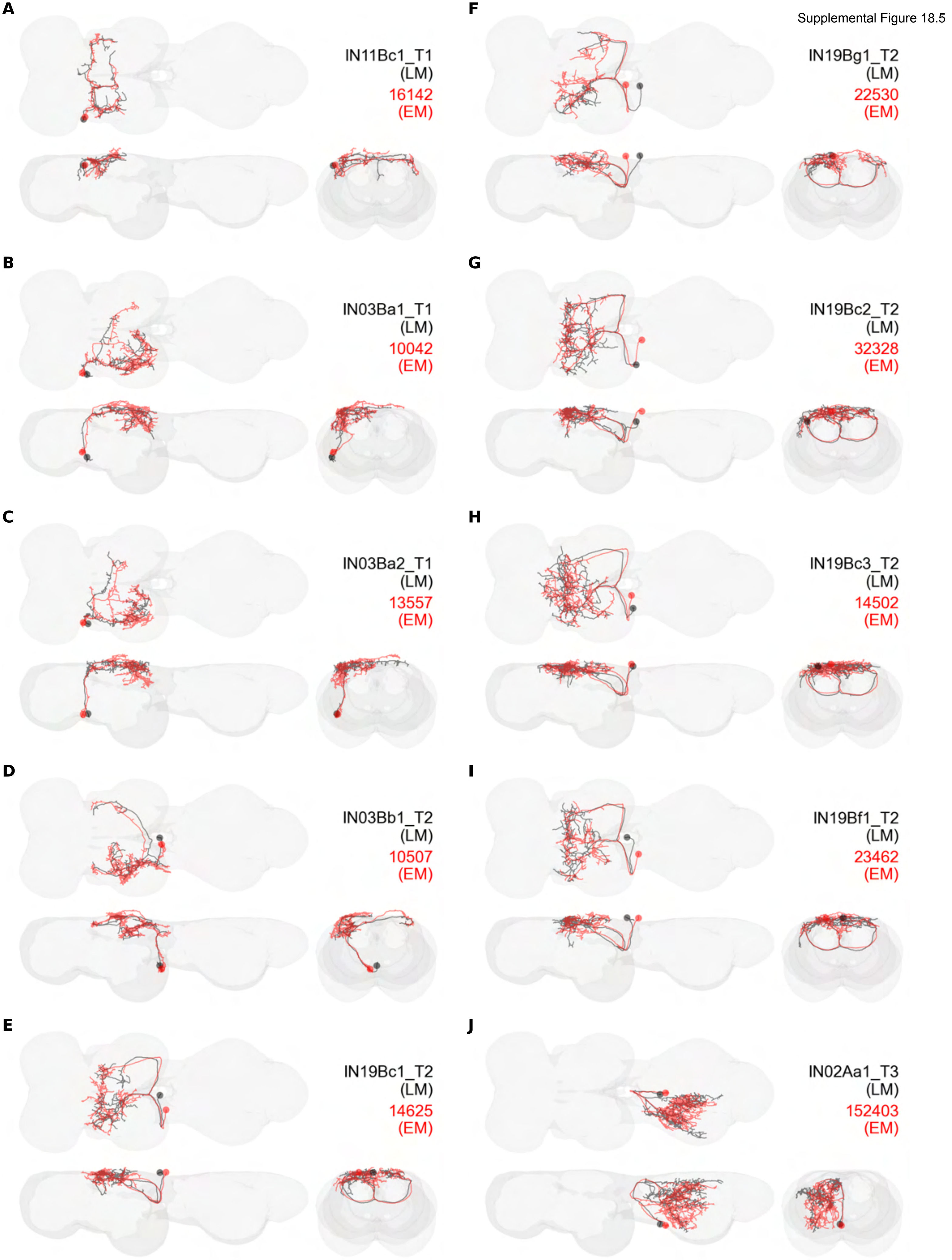

Supplemental Figure 18.6

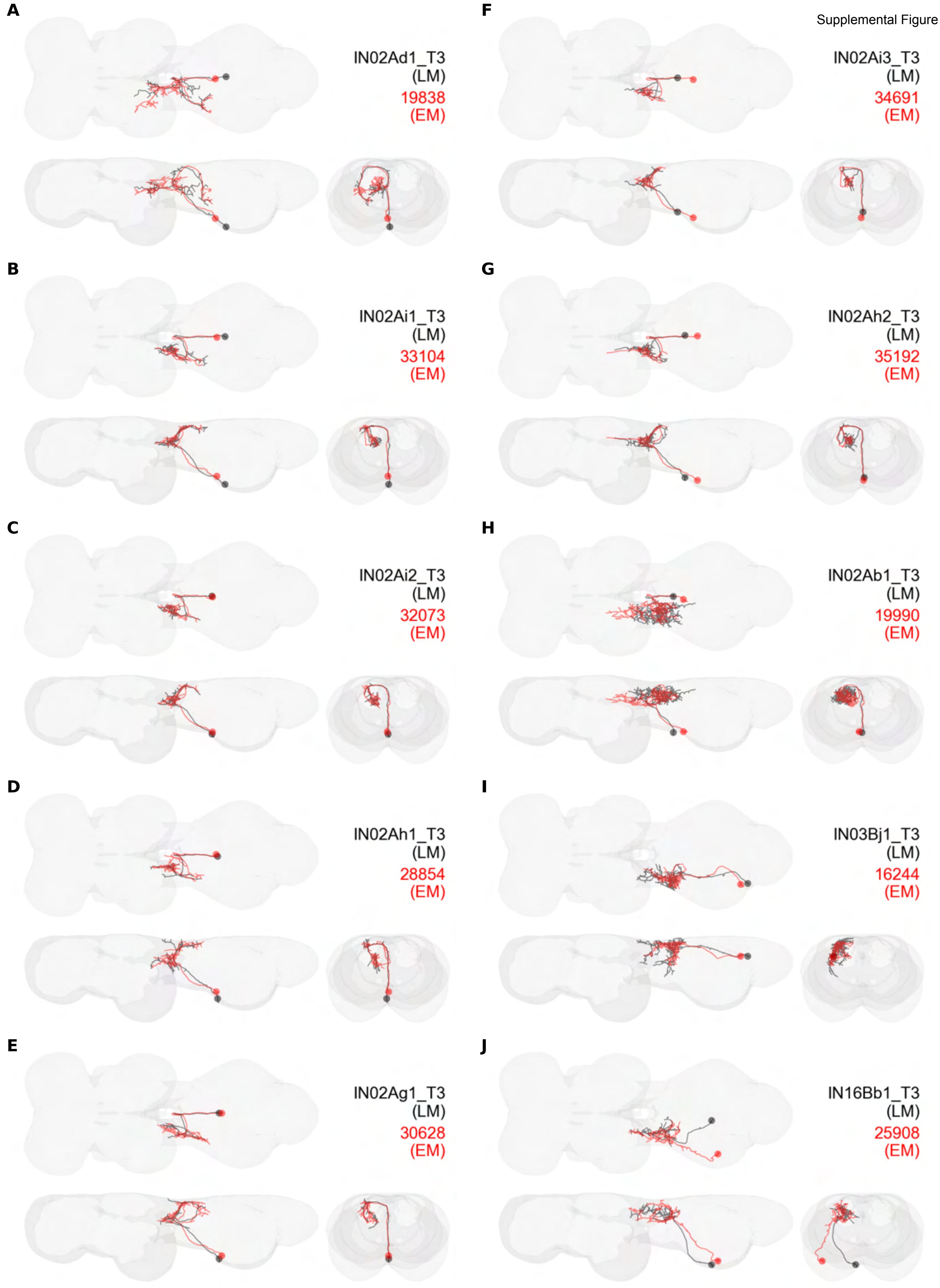

**A**

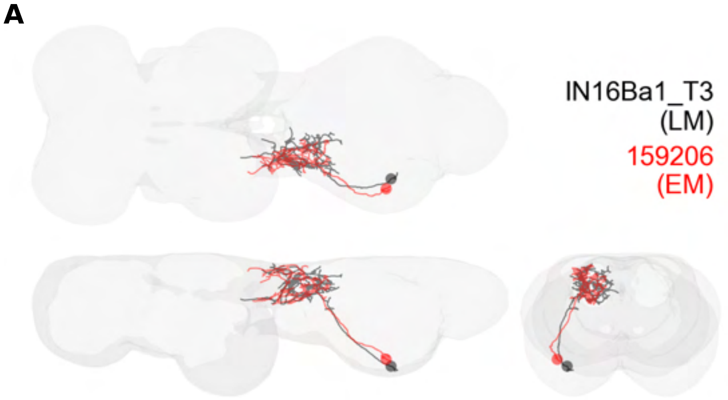


**B**

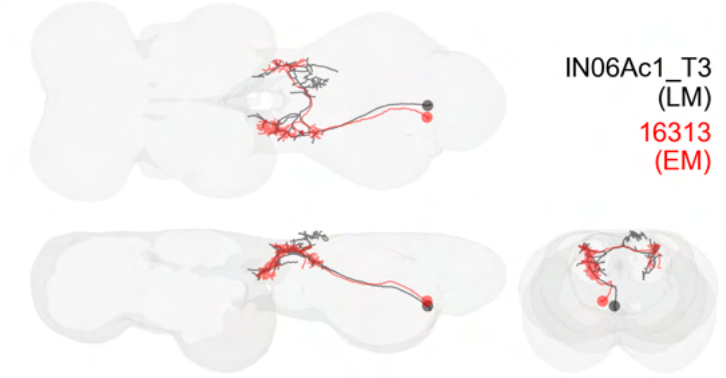

Supplemental Figure 18.8

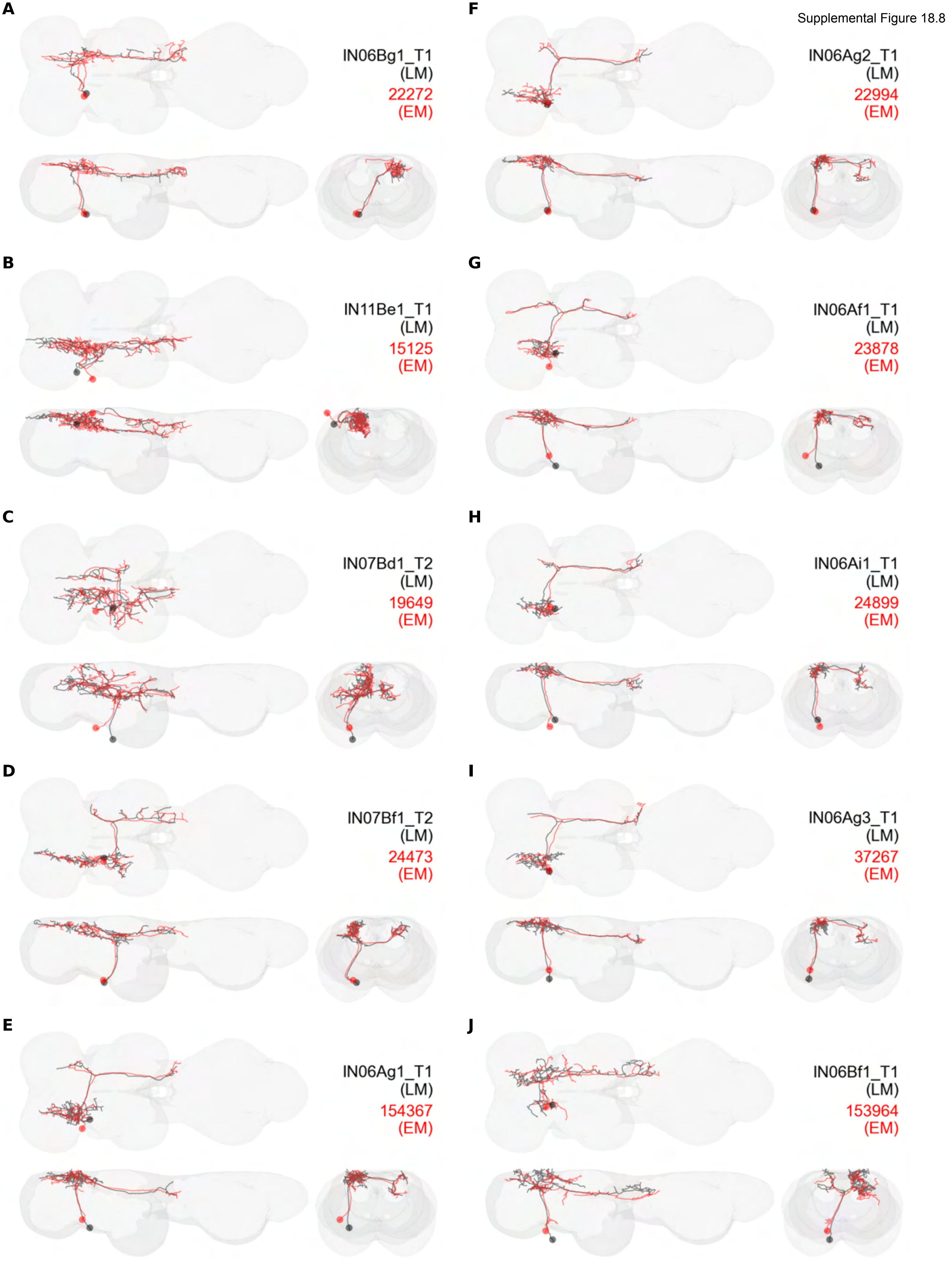

Supplemental Figure 18.9

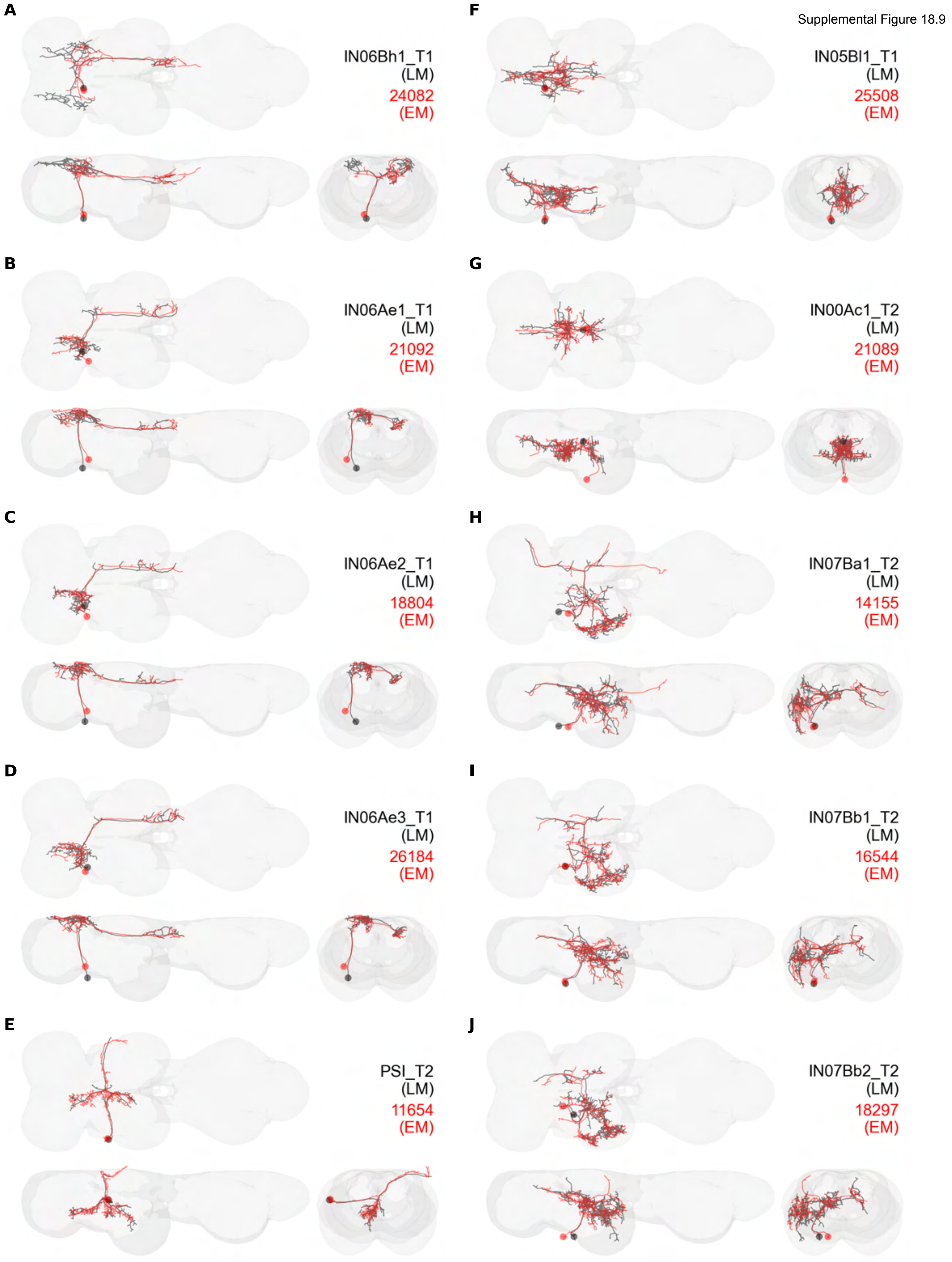

Supplemental Figure 18.10

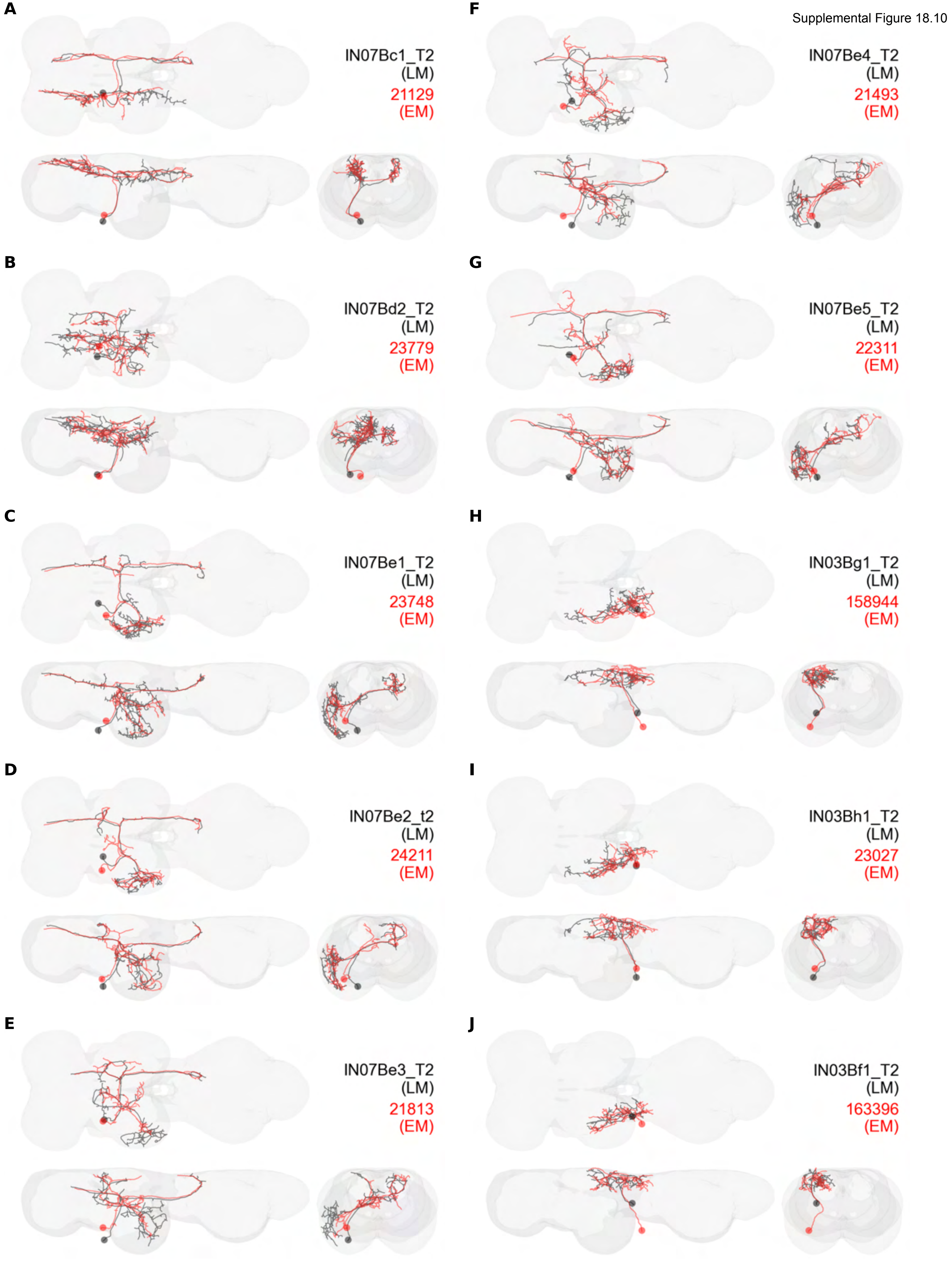

Supplemental Figure 18.11

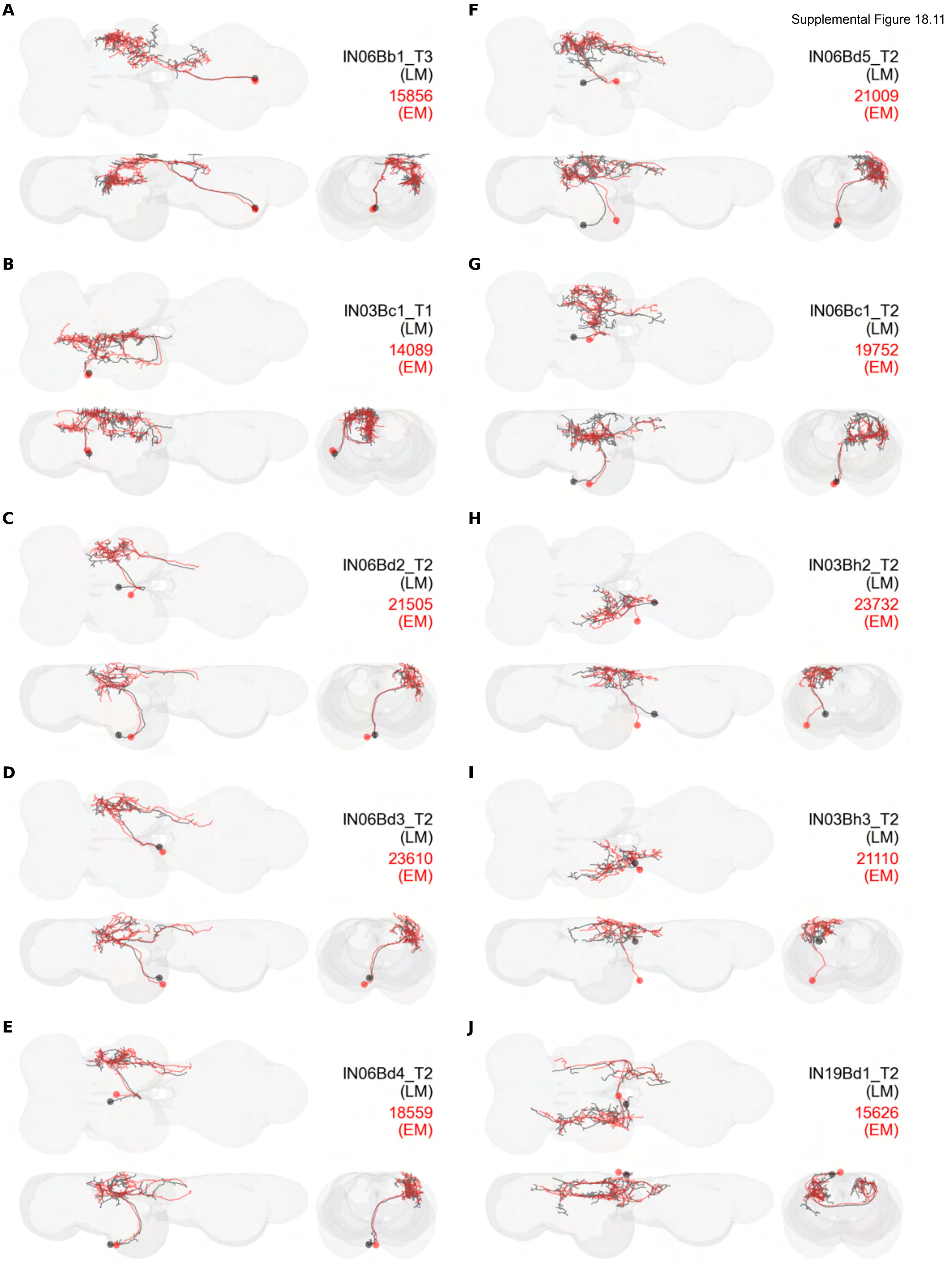

Supplemental Figure 18.12

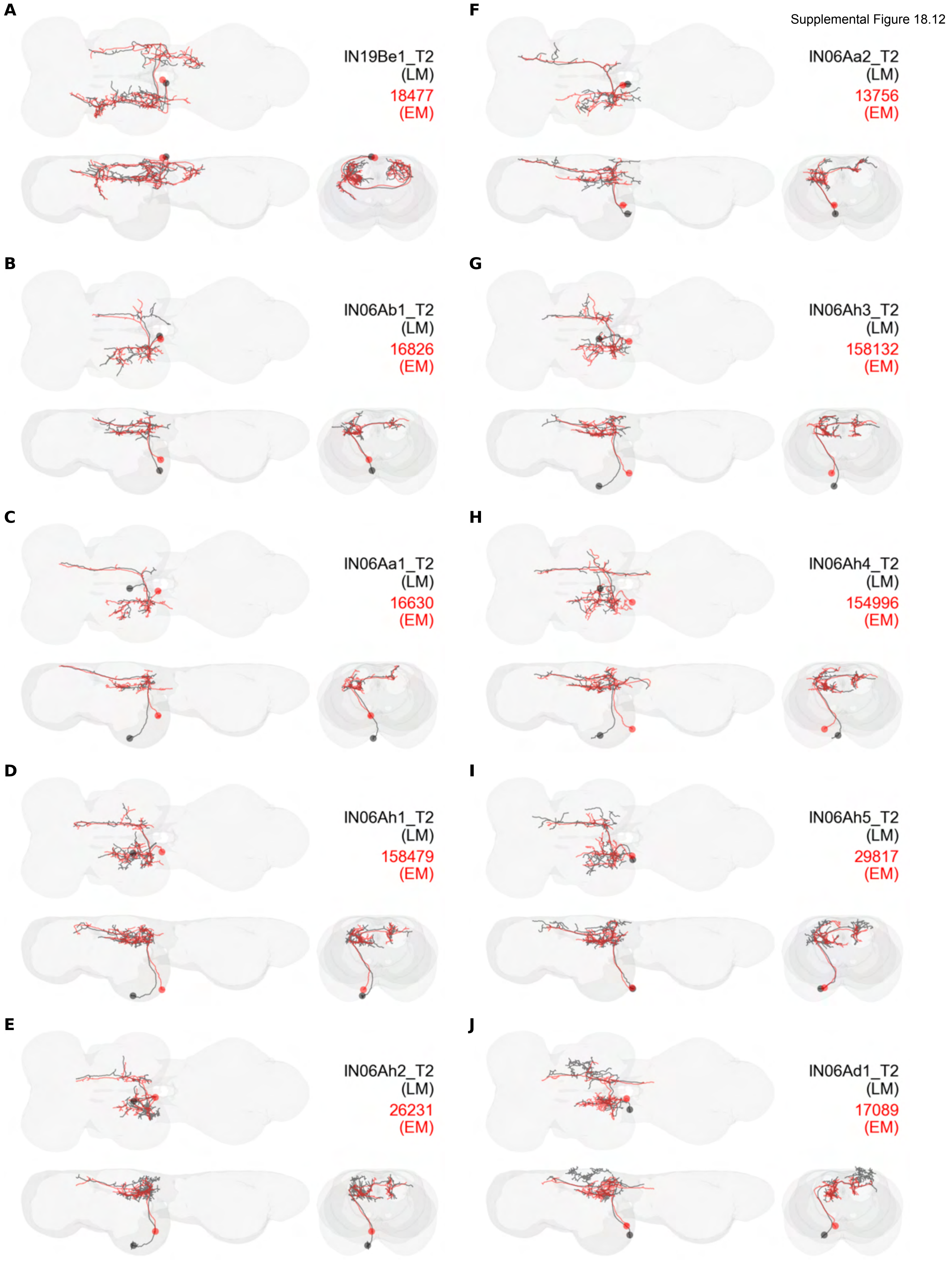

Supplemental Figure 18.13

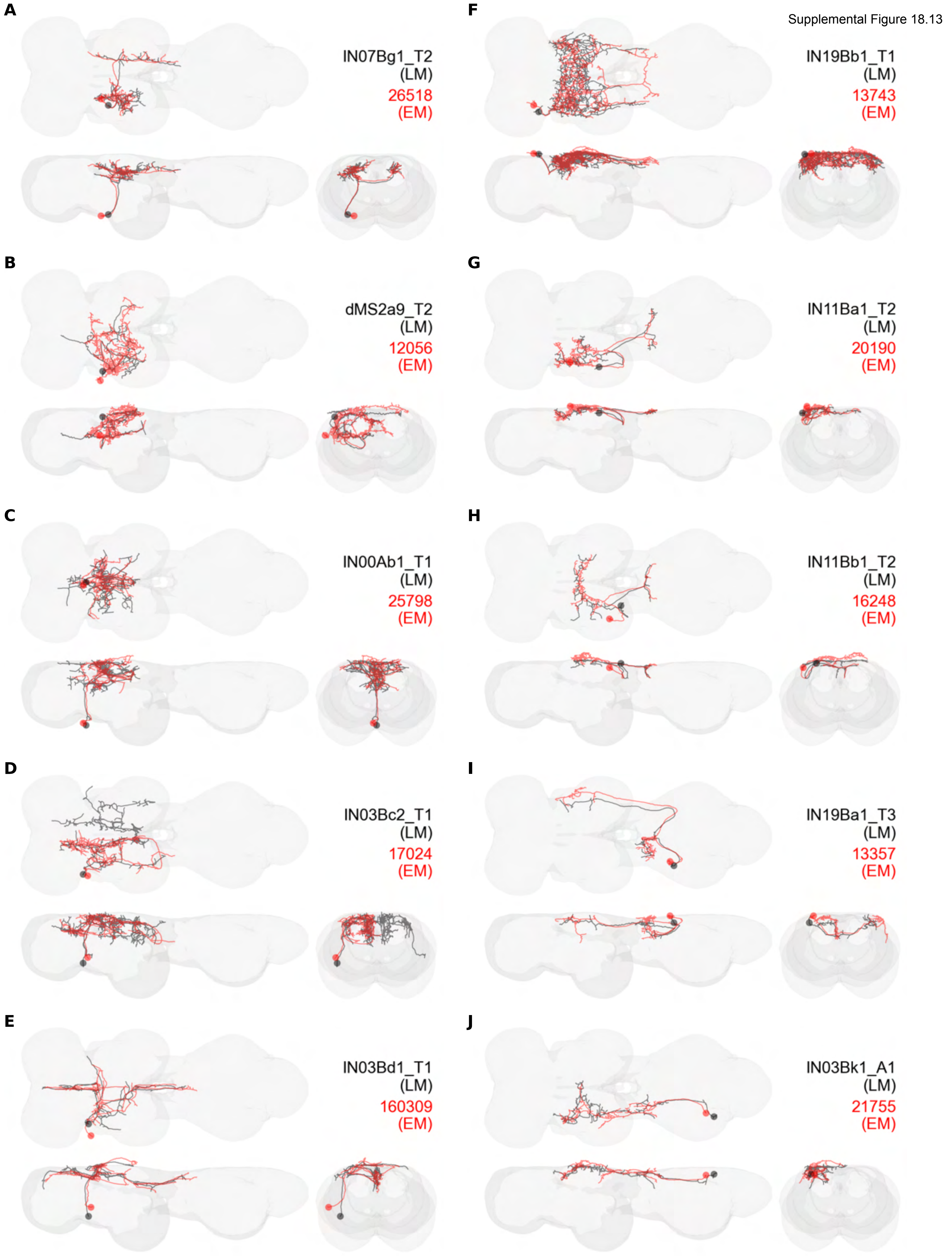

Supplemental Figure 18.14

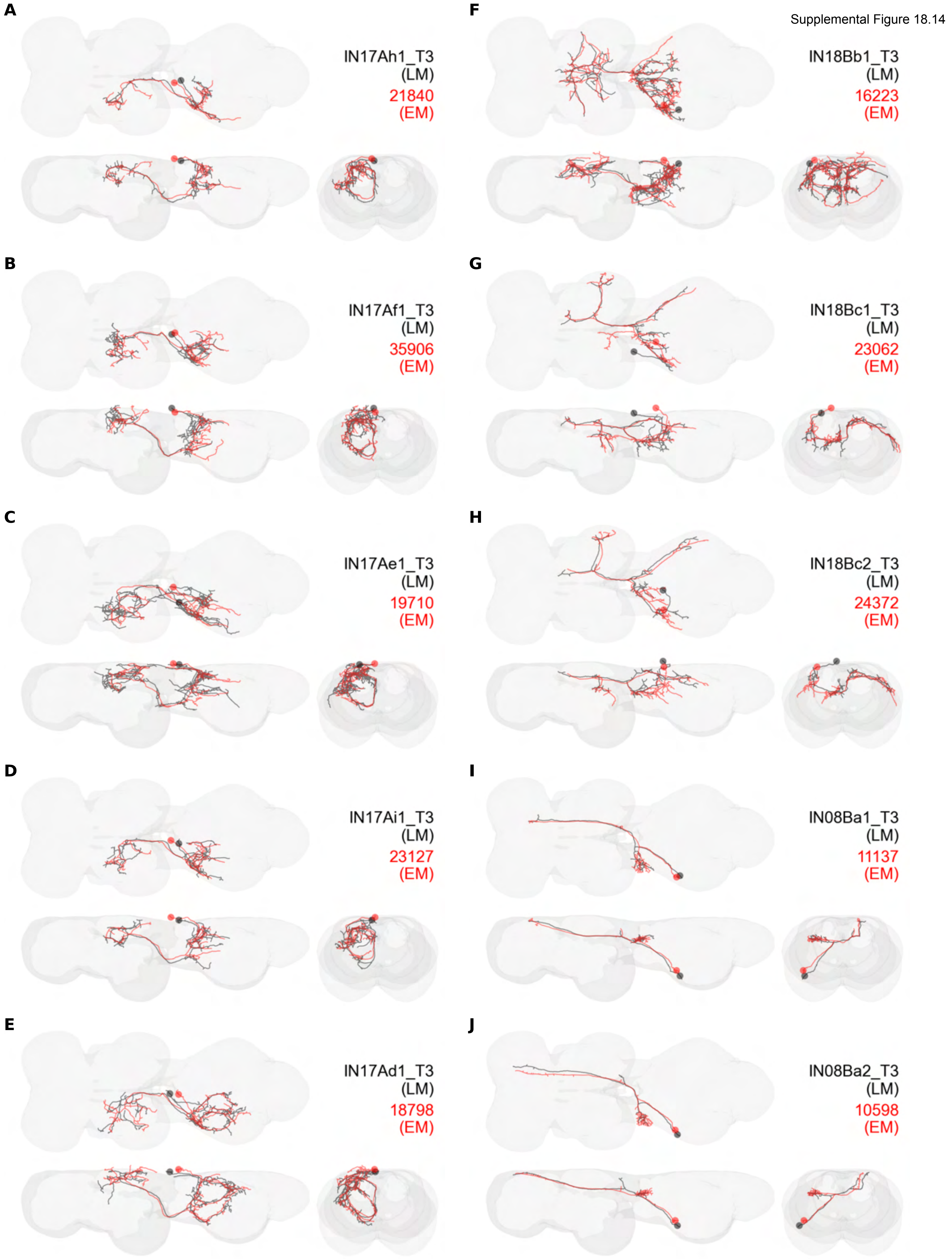

Supplemental Figure 18.15

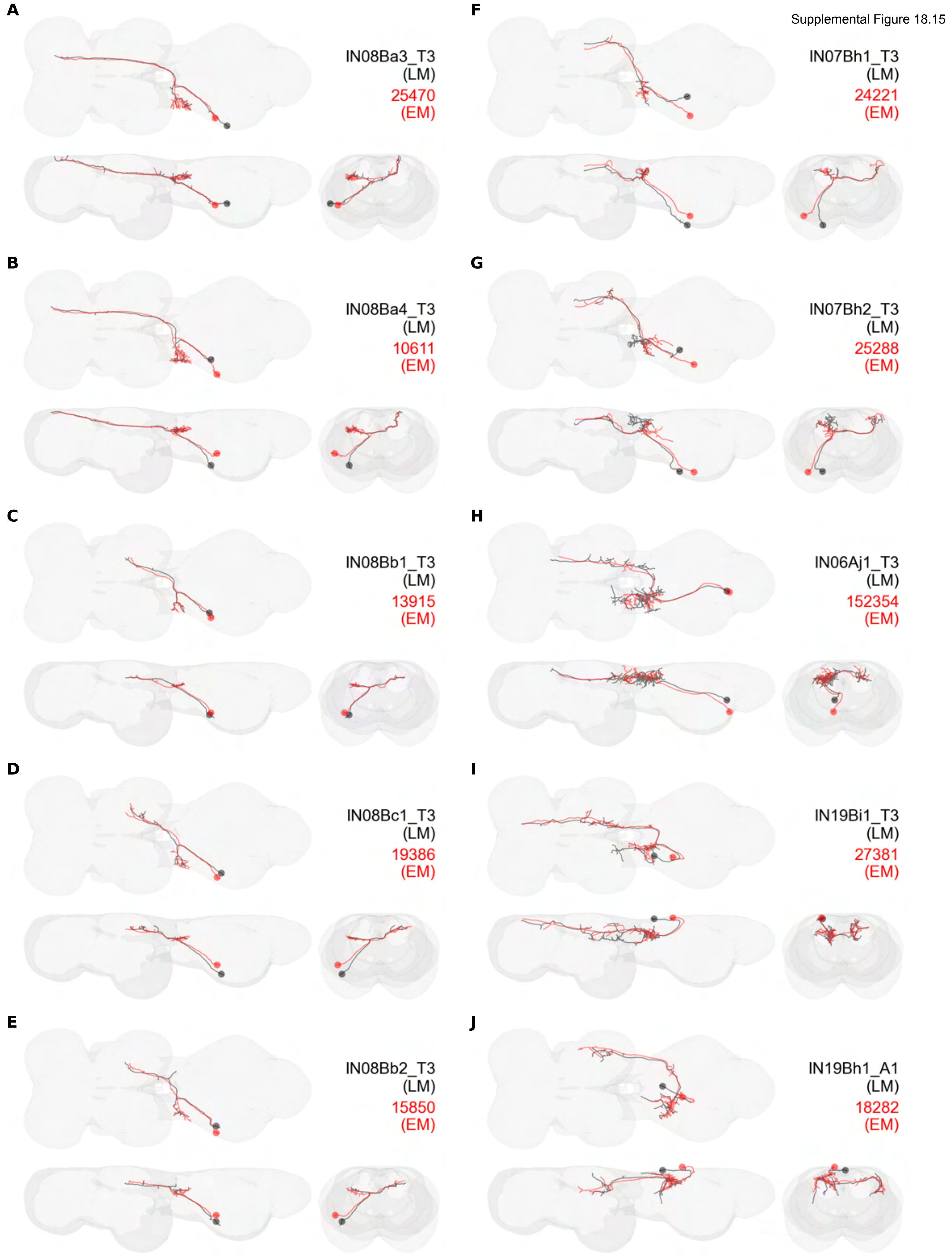

Supplemental Figure 18.16

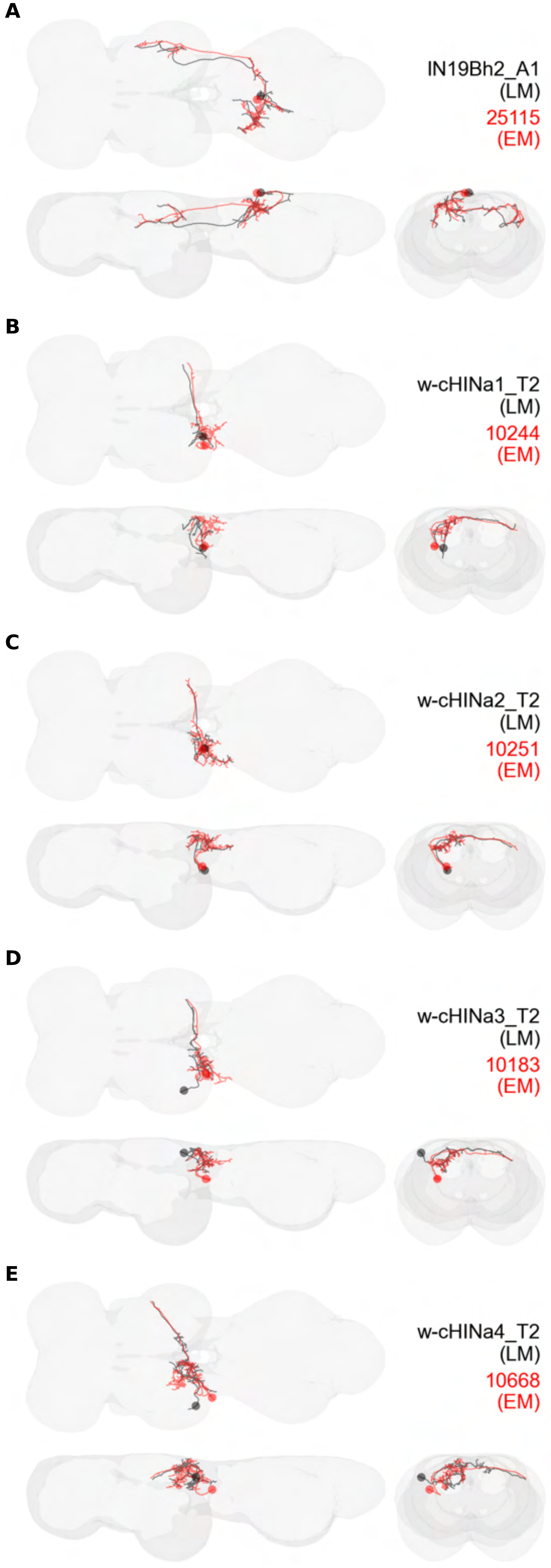

Supplemental Figure 18.17

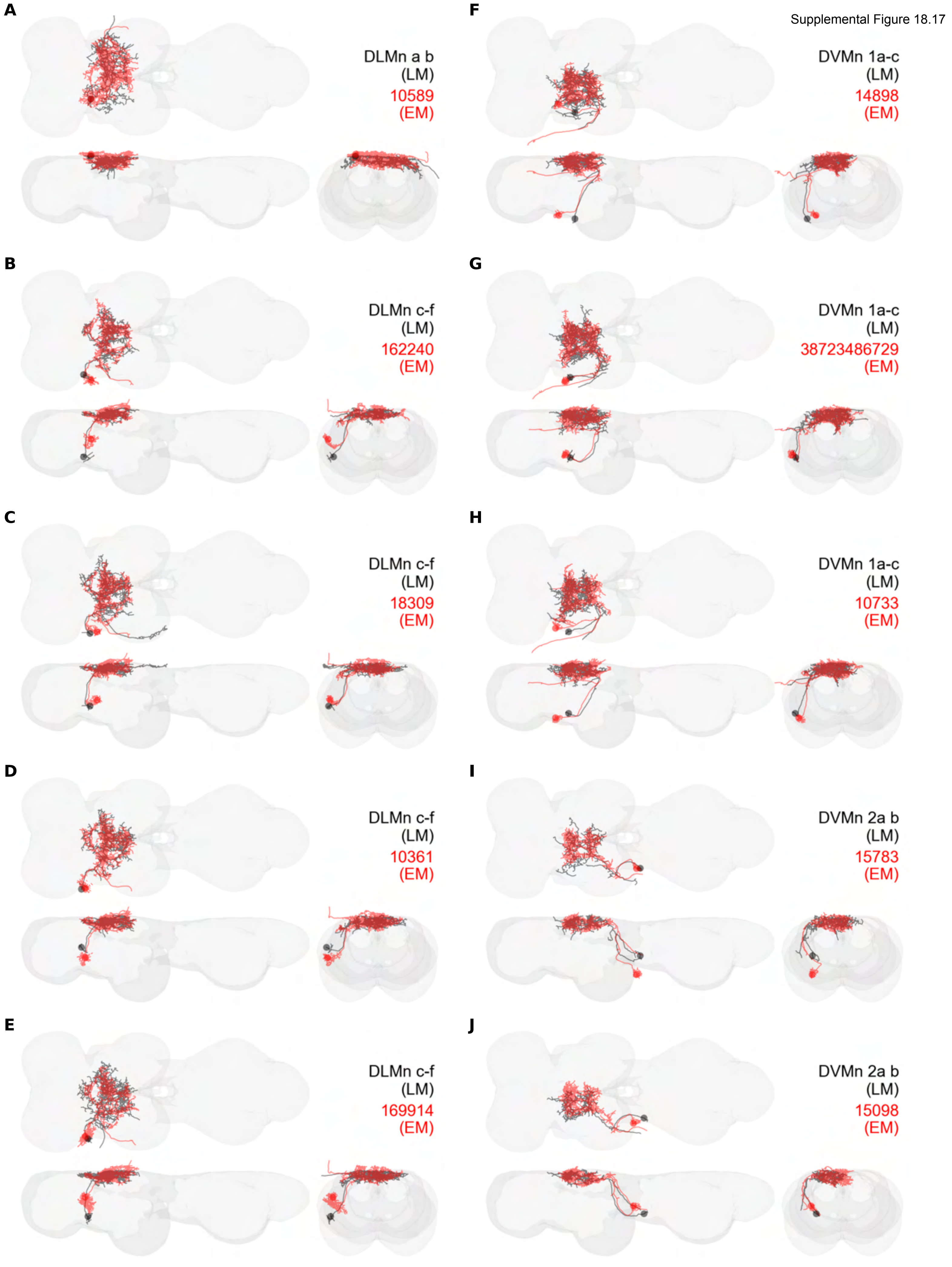

Supplemental Figure 18.18

**A**

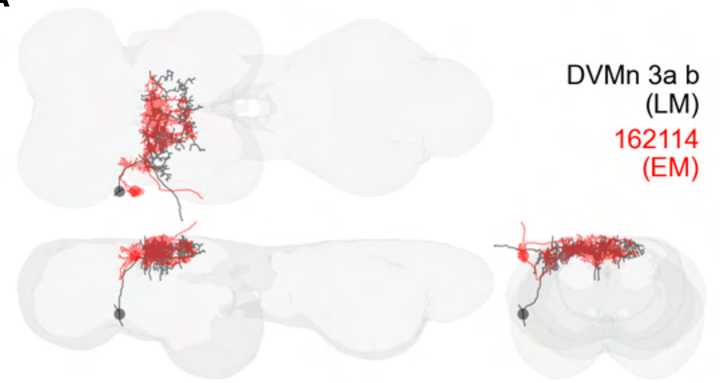


**B**

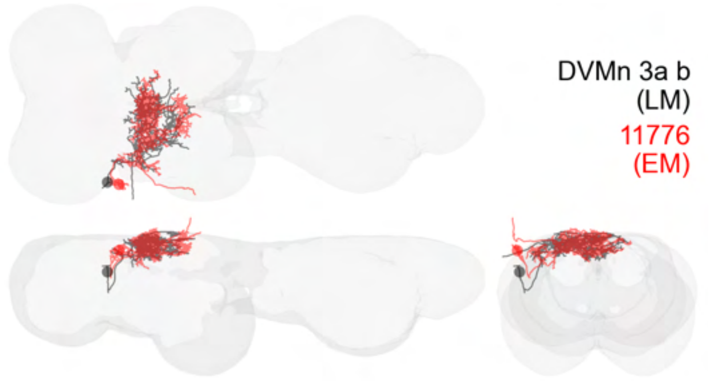

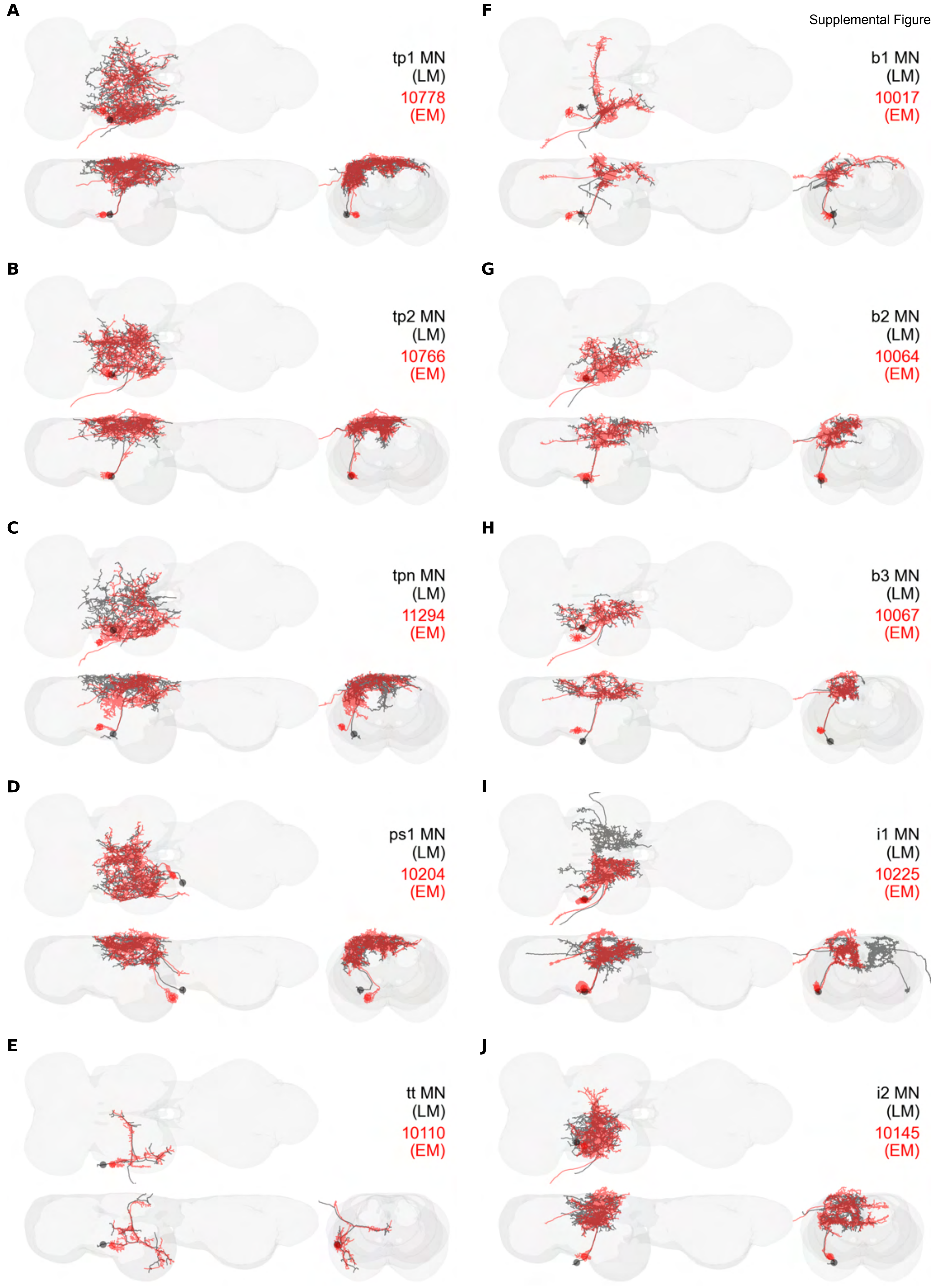
A
tp1 MN
(LM)
10778
(EM)
B
tp2 MN
(LM)
10766
(EM)
C
tpn MN
(LM)
11294
(EM)
D
ps1 MN
(LM)
10204
(EM)
E
tt MN
(LM)
10110
(EM)
F
b1 MN
(LM)
10017
(EM)
G
b2 MN
(LM)
10064
(EM)
H
b3 MN
(LM)
10067
(EM)
I
i1 MN
(LM)
10225
(EM)
J
i2 MN
(LM)
10145
(EM)

Supplemental Figure 18.20

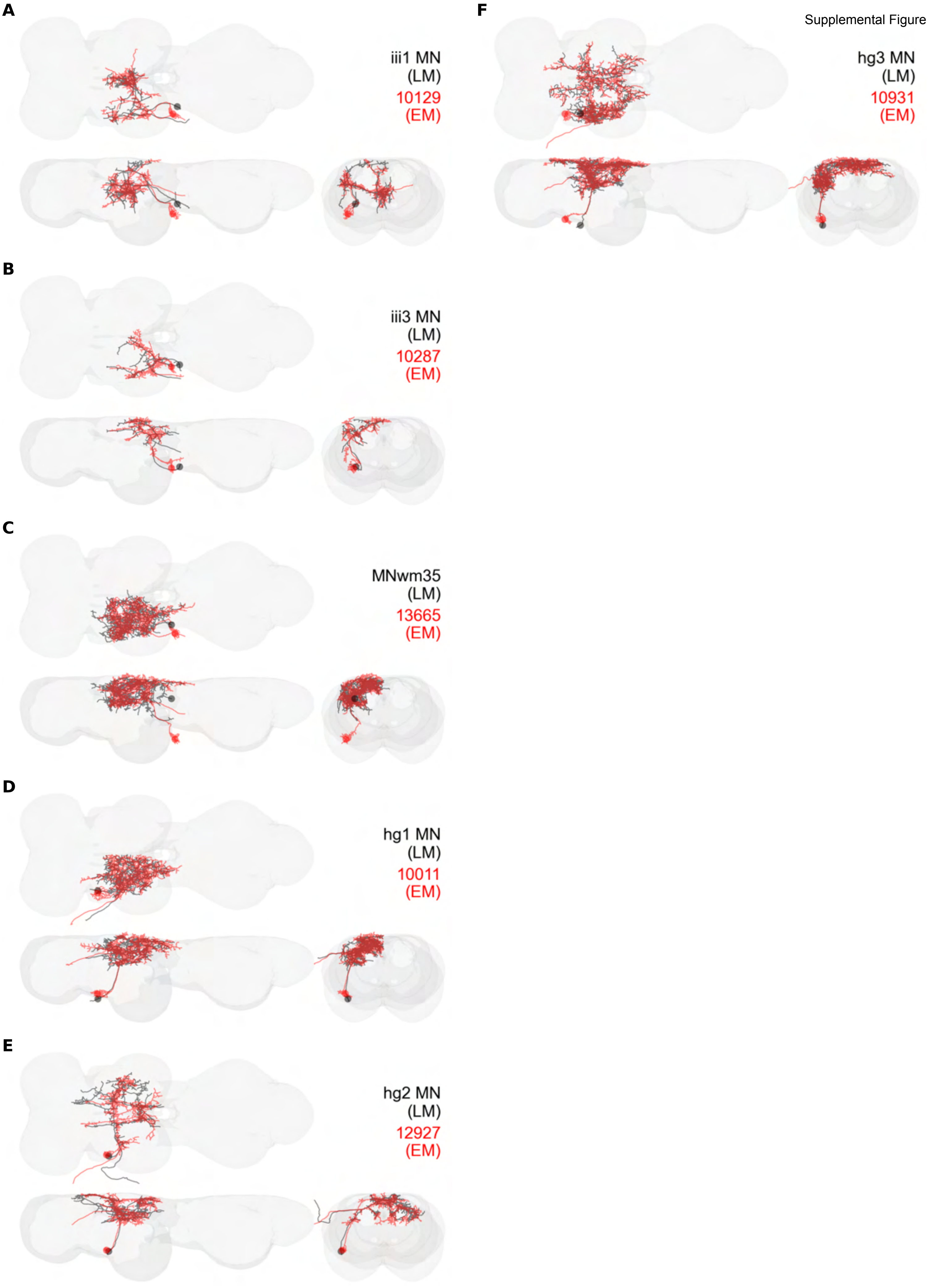

Supplemental Figure 18.21

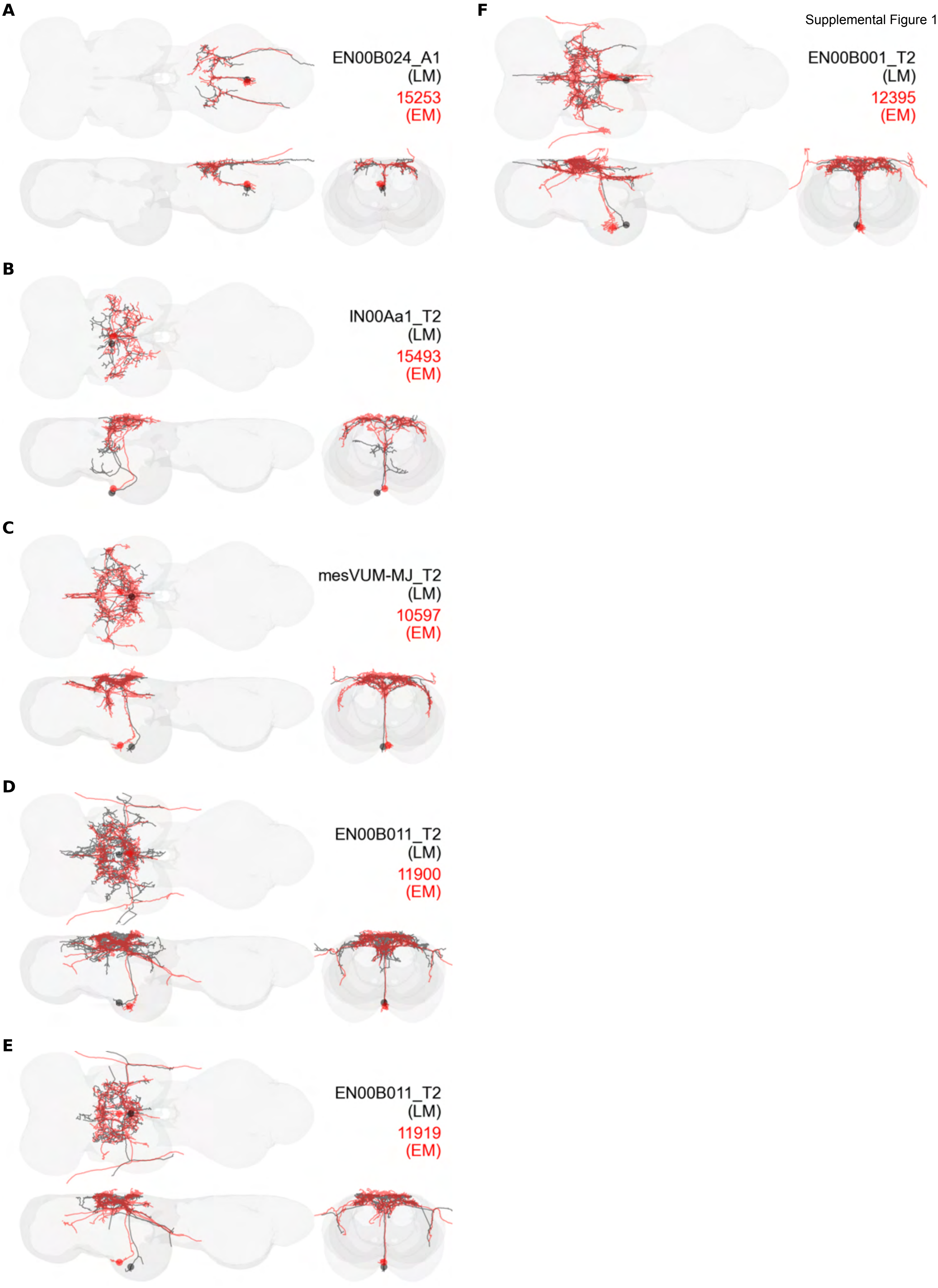
